# Decoding the Sequence Requirements for Translation Initiation

**DOI:** 10.64898/2026.05.12.723742

**Authors:** Bram M. P. Verhagen, David Liedtke, Lucía Barbadilla-Martínez, Carlos Alvarado, Valentyn Petrychenko, Michał Świrski, Micha Müller, Eivind Valen, Joseph D. Puglisi, Jeroen de Ridder, Niels Fischer, Marvin E. Tanenbaum

**Affiliations:** Oncode Institute, Hubrecht Institute–KNAW and University Medical Center Utrecht, Utrecht, the Netherlands; Oncode Institute, the Netherlands; Department of Bionanoscience, Kavli Institute of Nanoscience, Delft University of Technology (TU Delft), Delft, the Netherlands; Project Group Molecular Machines in Motion, Department of Physical Biochemistry, Max Planck Institute for Multidisciplinary Sciences, Göttingen, Germany; Center for Molecular Medicine (CMM), University Medical Center Utrecht, Utrecht, the Netherlands; Princess Máxima Center for Pediatric Oncology, Utrecht, the Netherlands; Department of Structural Biology, Stanford University School of Medicine, Stanford, CA, USA; Institute of Genetics and Biotechnology, Faculty of Biology, University of Warsaw, Warsaw, Poland; Computational Biology Unit, Department of Informatics, University of Bergen, Bergen, Norway; Michael Sars Centre, University of Bergen, Bergen, Norway; Department of Biosciences, University of Oslo, Oslo, Norway

## Abstract

Accurate selection of start codons by ribosomes is a fundamental determinant of proteome composition. Although the ‘Kozak sequence’—an 8-nucleotide sequence flanking the start codon—has long been viewed as the primary determinant of initiation in eukaryotes, it fails to explain the large diversity of start codon usage across transcripts. Here we combine massively parallel reporter assays, bioinformatics, machine learning, single-molecule imaging and cryo-electron microscopy to define the ‘extended translation initiation sequence (eTIS)’, an ∼80-nucleotide sequence surrounding the start codon that governs initiation efficiency. A deep-learning model trained on eTIS features accurately predicts translation initiation across transcripts. Unexpectedly, we find that the Kozak sequence is not optimal for initiation as is widely presumed, and we identify the origin of this discrepancy. eTIS nucleotides that promote efficient initiation are enriched in the human transcriptome and are evolutionarily conserved, underscoring their functional importance. Biophysical and structural analyses reveal that specific eTIS residues—including the key +6 position and residues in the mRNA entry and exit channel—engage ribosomal proteins, rRNA and initiation factors to promote start codon recognition by stabilizing the ribosome at the start codon and facilitating the structural transitions required for initiation. Finally, optimization of the eTIS markedly enhances translational fidelity and protein output from therapeutic mRNAs, highlighting its practical utility. Together, these findings redefine the sequence logic of translation initiation and establish a framework for precise control of protein expression.

## Introduction

Accurate and efficient recognition of start codons by the ribosome is essential for gene expression regulation and for defining proteome composition. In eukaryotes, start codon identification requires the ribosome to scan the 5′ untranslated region (UTR) of the mRNA in a 5’-to-3’ direction until it encounters a start codon, typically an AUG codon, where translation initiation can occur (Hinnebusch, 2014). Start codon selection by the ribosome determines the protein’s sequence by fixing both the reading frame and the N-terminus of the protein. Many mRNAs, however, contain multiple possible start codons, each potentially driving the synthesis of a distinct protein isoform that can have altered localization, stability or activity (Ingolia et al., 2011; Lee et al., 2012; Ly et al., 2025). Alternative start codons can also tune the expression levels of the protein encoded by the main open reading frame (ORF), for example through introduction of regulatory short ORFs, such as upstream ORFs (uORFs) (Dever et al., 2023). Alternative start codons can be either AUGs or near-cognate start codons (e.g. CUG, GUG). Such alternative start codons are abundantly present in most mRNAs, but only a small subset of them is used for translation initiation (Kearse and Wilusz, 2017).

An in-depth understanding of the mechanisms that control start codon selection would improve the molecular understanding of translation and facilitate identification of novel ORFs and potential microproteins in the genome (Chothani et al., 2023; Sandmann et al., 2023). In addition, such improved understanding would aid in the optimal design of therapeutic mRNAs, for which precisely controlled translation initiation is essential both for high level expression and for safety by minimizing the expression of ‘off-target’ proteins produced through initiation at alternative start codons.

A key sequence feature that is thought to mark potential start codons as sites of translation initiation is the Kozak consensus sequence (‘Kozak sequence’) — GCC(G/A)CC**<u>AUG</u>**GC—, discovered in the 1980’s through sequence comparison of a few hundred eukaryotic mRNAs (Kozak, 1987a). The Kozak sequence describes the most common nucleotide at each position in the region -6 to +5 relative to the start codon, known as the Translation Initiation Sequence (TIS) (with position +1 referring to the A of the AUG), and is widely assumed to be the most optimal TIS for start codon recognition. Indeed, high-throughput analysis confirmed that the Kozak sequence represents a potent enhancer of start codon recognition (Noderer et al., 2014), with the -3 and +4 positions representing especially important nucleotides (Kozak, 1986; Noderer *et al*., 2014). However, fewer than 1% of human genes harbor the Kozak sequence, and many bona fide start codons are embedded in TISs that deviate substantially from the Kozak sequence (Hernandez et al., 2019). Furthermore, we previously found that even when a start codon is flanked by the Kozak sequence that a substantial fraction of ribosomes initiate at alternative start codons (Boersma et al., 2019). These observations suggest that the Kozak sequence is not sufficient to explain the full spectrum of translation initiation efficiencies observed in the transcriptome, and suggest that additional mechanisms shape start codon selection.

Here, using a massively parallel reporter assay and bioinformatic analysis we show that start codon recognition is governed not only by the Kozak sequence, but by a conserved ‘extended TIS (eTIS)’ that spans approximately 40 nucleotides both upstream and downstream of the start codon. Using this data, we developed *RiboScanner*, a convolutional neural network model, that enables markedly improved start codon prediction and rational optimization of therapeutic mRNAs. Using single-molecule imaging and cryo-electron microscopy, we elucidate the molecular mechanism of eTIS-mediated translation initiation.

## Results

### Quantification of start codon recognition

To identify novel mRNA features that control start codon recognition efficiency, we developed ’RiboScan’, a massively parallel reporter assay that enables high-resolution exploration of mRNA sequence space. Earlier studies aimed at studying sequences that promote translation initiation have employed fluorescence reporters in which expression of a reporter gene sequence (e.g. GFP or a FLAG-tag) is driven by variable 5’UTRs, TISs or start codons (Akirtava et al., 2025; Diaz de Arce et al., 2018; Dvir et al., 2013; Jia et al., 2020; Noderer *et al*., 2014; Strayer et al., 2023). While such studies have successfully identified important regulatory elements of translation efficiency (i.e. ribosome recruitment to the mRNA), this reporter design is not ideal for identification of mRNA features that control start codon selection; if a particular sequence variant increases start codon selection efficiency, from, for example, 90% to 99% efficiency, the expected change in GFP expression is only 1.1-fold, an effect size that may escape detection. Indeed, in earlier studies even mutation of most TIS nucleotides did not result in detectable changes in reporter expression (Hernandez *et al*., 2019; Kozak, 1986; 1987b; 1997). To overcome this challenge, we aimed to design an alternative fluorescence reporter, in which GFP expression is exclusively driven by ribosomes that *fail* to recognize a start codon. In this design, if start codon recognition efficiency changes from 90% to 99%, the number of ribosomes synthesizing GFP will decrease from ∼10% to ∼1%, a 10-fold difference in GFP expression, providing greatly improved detection sensitivity. To design this reporter, we leveraged the fact that, if ribosomes fail to recognize a start codon during 5’UTR scanning (‘leaky scanning’), they continue scanning the mRNA and initiate at a downstream start codon (Hinnebusch, 2017). We engineered a fluorescence reporter in which GFP is encoded ∼100 nucleotides downstream of the first start codon (‘main start codon’) in an alternative reading frame (frame +1), such that ribosomes that initiate at the main start codon (frame 0) will translate GFP in the incorrect frame, producing a non-fluorescent polypeptide (Fig. 1a). To ‘capture’ ribosomes that undergo leaky scanning of the main start codon, three consecutive AUG sequences were placed immediately upstream and in frame with GFP (frame +1) (Fig. 1a). Thus, in this design GFP expression reports on leaky scanning of the main AUG and is inversely correlated with start codon recognition efficiency. We refer to this design as the ‘RiboScan reporter’.

**Figure 1.**
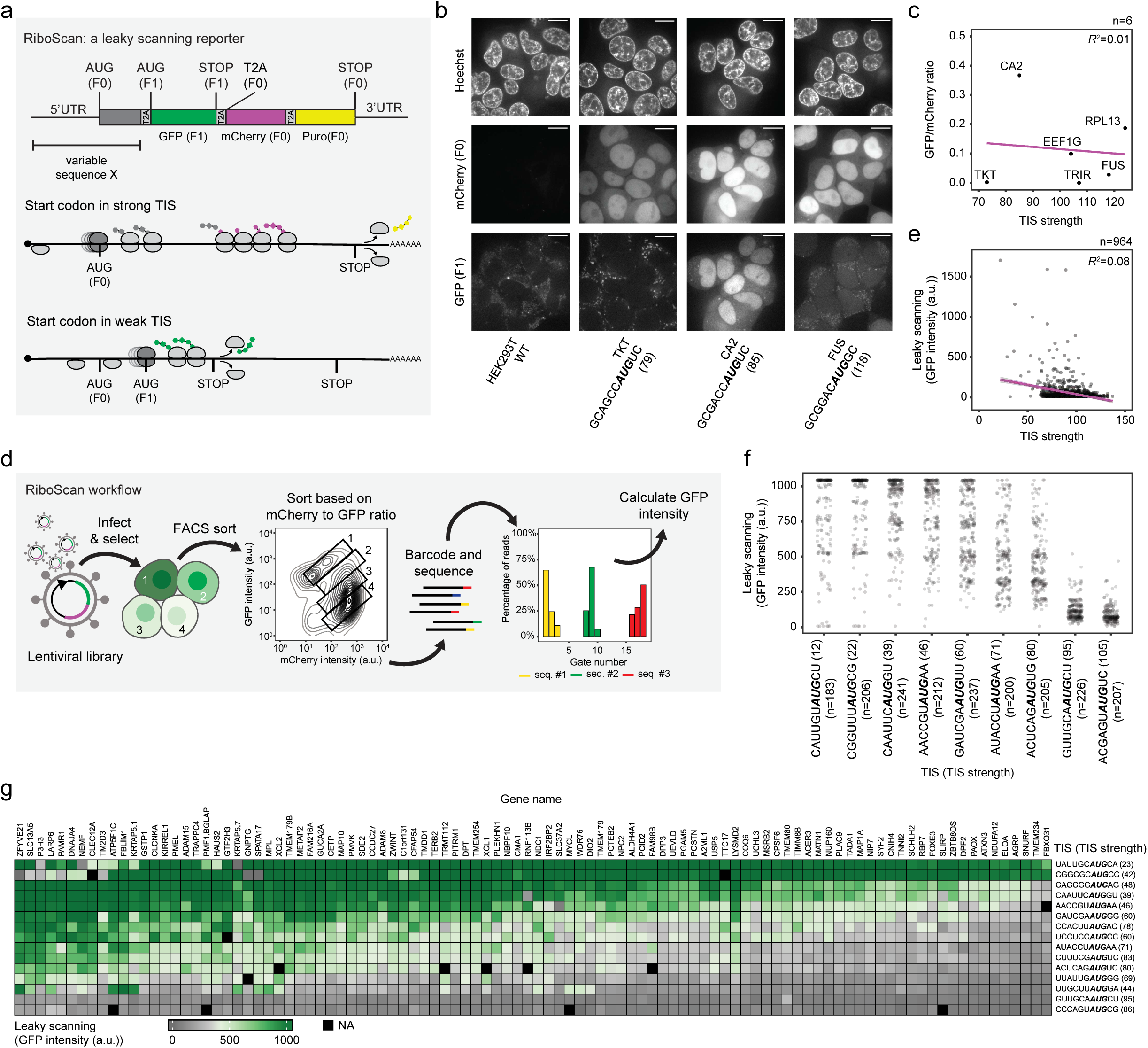
Gene sequences beyond the TIS control start codon recognition. **a**, Schematic of the RiboScan reporter mRNA. (Top) Reading frame is indicated for different mRNA features. T2A ‘Stop-Go’ sequences are indicated in grey. The first T2A sequence is in frame +1 (F1), the latter two in the main frame (F0). Middle, when the first start codon (main start codon, in F0) is positioned in a strong sequence context, ribosomes initiate on the first start codon and produce mCherry and the enzyme providing puromycin resistance. Bottom, when the first start codon is positioned in a poor sequence context, ribosomes continue scanning and initiate on the downstream start codon (F1) to produce GFP. Dark gray ribosome are undergoing translation initiation. **b**, Example microscopy images of endogenous genes in which the RiboScan reporter was inserted. The gene name, TIS and TIS strength (between brackets, obtained from Noderer *et al*. (Noderer *et al*., 2014)) are indicated. Scale bars, 10µm. **c**, The RiboScan reporter is inserted into 6 endogenous genes using CRISPR. For each targeted gene, the TIS strength and the ratio of F1 (GFP) and F0 (mCherry) intensities is depicted. A linear regression fit to the data is shown (purple line). **d**, Schematic of high-throughput analysis of the RiboScan reporter assay. Four hypothetical cells are shown. **e**, TIS strength is plotted against GFP intensity for 964 RiboScan reporters containing endogenous mRNA sequences. A linear regression fit to the data is shown in purple, with grey shaded region indicating the 95% confidence interval. **f**, Nine different TISs were introduced into indicated number of endogenous mRNA sequences (between brackets) and GFP expression of the RiboScan reporter was determined. Each dot presents a single mRNA sequence. **g**, Heatmap representing GFP expression of the RiboScan reporter of indicated mRNA sequences (columns) combined with indicated TISs (rows). The strength of each TIS is indicated between brackets. Black boxes represent missing data points. **c**, **e**, **f**, **g**, represent average values of two independent experiments.

To enable quantitative measurements of leaky scanning, and thus start codon recognition efficiency, using the RiboScan reporter, a number of additional features were introduced into the RiboScan reporter mRNA. First, a second fluorescent protein (mCherry) was introduced downstream of the GFP in frame 0 to normalize GFP expression for variations in reporter mRNA levels and overall translational efficiency. Second, a puromycin resistance gene was introduced to facilitate rapid selection of RiboScan reporter expressing cells. Third, T2A ‘stop-go’ sequences (Liu et al., 2017) were introduced upstream of GFP (frame +1) and both upstream and downstream of mCherry (frame 0) to ensure that GFP and mCherry are not fused to other amino acid sequences that could affect fluorescent protein stability and thereby impact the read-out of start codon recognition efficiency. For the same reason, a stop codon was introduced directly downstream of the GFP coding sequence in the GFP reading frame. Finally, to ensure that the RiboScan reporter reports exclusively on leaky scanning and not on upstream translation initiation in the 5’UTR (in the GFP frame), we introduced an additional stop codon in the GFP frame immediately upstream of the GFP sequence. Together, these modifications allow a highly sensitive and quantitative read-out of start codon recognition efficiency.

### The TIS alone does not explain start codon recognition

Using the RiboScan reporter, we first examined start codon recognition efficiency on endogenous mRNAs. The RiboScan reporter was introduced into 6 endogenous genes in human HEK293T cells using CRISPR, and GFP and mCherry expression levels were determined for each gene by microscopy. Clear GFP expression was observed in 4 of 6 targeted genes (Fig. 1b, Supplementary Fig. 1a), confirming that leaky scanning of the main start codon is pervasive, consistent with our previous work assessing start codon recognition by single-molecule imaging (Boersma *et al*., 2019). Interestingly, the level of leaky scanning (GFP expression) varied greatly between endogenous mRNAs. To assess if the sequence of the TIS determines the degree of leaky scanning, we made use of a previously-developed ‘TIS strength’ score ranging from 0-150 (Noderer *et al*., 2014). Surprisingly, we found that the efficiency of start codon recognition in endogenous mRNAs did not correlate well with the TIS strength (Fig. 1c), indicating that additional mRNA features must exist that determine start codon recognition efficiency.

To identify novel mRNA features that control start codon recognition efficiency, we inserted the RiboScan reporter into a lentiviral vector to allow high-throughput testing of the impact of different mRNA sequences on start codon recognition efficiency (Fig 1d). We introduced the 5’UTR sequence and the first 18nt of the coding sequence of 1000 endogenous genes into the lentiviral RiboScan reporter. Genes were included that had a 5’UTR length of between 6-90 nucleotides and that did not contain alternative AUG start codons in any of the frames, or stop codons in the mCherry and GFP frame to simplify data interpretation. HEK293 cells were transduced with the RiboScan reporter library at low MOI (<10% of cells infected, see Methods) to ensure each cell received only a single reporter variant. Cells transduced with the RiboScan reporter library were then FACS sorted into at least 12 different populations based on the GFP-mCherry intensity ratio and each population was analyzed by sequencing to determine the average normalized GFP expression associated with each mRNA sequence variant (Supplementary Fig. 1c, Supplementary Fig. 1a, see Methods). Data were highly reproducible between biological repeats (Supplementary Fig. 1d,e), and calculated GFP expression correlated well with the measured GFP expression for sequences tested individually (Supplementary Fig. 1f). Consistent with the analysis of endogenous genes, analysis of GFP expression levels from the lentiviral RiboScan reporter revealed that leaky scanning levels were highly variable between mRNA sequences, and that GFP expression levels correlated poorly with TIS strength (R^2^ = 0.08) (Fig. 1e), confirming that TIS strength alone does not well explain differences in start codon recognition efficiency among endogenous mRNA sequences. Indeed, even when the endogenous TIS was replaced by a single, constant TIS in hundreds of mRNA sequences, GFP expression was still highly variable among different mRNAs (Fig. 1f), further confirming that sequences beyond the TIS regulate start codon recognition. When different TISs were compared in this assay, highly variable start codon recognition was observed for all tested TISs. Nonetheless, TISs with a high TIS strength did, on average, show improved start codon recognition (Fig. 1f), indicating that TIS strength does contribute to start codon recognition efficiency somewhat, consistent with previous studies (Kozak, 1986; Noderer *et al*., 2014). To systematically assess the contribution of the TIS and non-TIS features to start codon recognition efficiency, we introduced 26 different TISs into a set of 300 endogenous gene sequences and determined GFP expression levels. While mRNAs with a strong TIS tended to show lower GFP expression on average, as before, GFP expression was also highly dependent on gene sequence beyond the TIS, with most genes showing consistently low, medium or high level of GFP expression, irrespectively of the incorporated TIS (Fig. 1g, Supplementary Fig. 1g-i). We confirmed that differences in GFP expression were due to differences in leaky scanning, rather than to initiation on alternative, upstream start codons (Supplementary Fig. 1j,k). Moreover, start codon recognition efficiency for different genes correlated well across cell types (Supplementary Fig. 1l, m), confirming the generality of these findings. Together, these results show that start codon recognition efficiency is heterogeneous among mRNA sequences, an effect that is not fully explained by variations in TIS strength.

### An extended TIS (eTIS) governs start codon recognition

To identify mRNA features that could explain variable start codon recognition efficiency, we examined potential differences in transcription start site (TSS) usage and RNA structure, but found that neither of these features explained GFP expression level variation (Supplementary Fig. 2a,b). Very short 5’UTR lengths (<12 nts) did affect start codon recognition efficiency, as was shown previously (Gu et al., 2021; Kozak, 1991), but no correlation between 5’UTR length and GFP expression was observed for mRNAs with 5’UTR lengths of ≥12 nts (Supplementary Fig. 2c,d). To locate the region within the mRNA that is most important for control of start codon recognition, we next performed a scanning mutagenesis analysis for >100 mRNAs, mutating blocks of 5 nts along the entire 5’UTR and first 93 nts of the coding sequence. Such mutations did not alter the TSS of the reporter, even when introduced very close to the TSS (Supplementary Fig. 3a). In cases where mutations introduced either AUG or stop codons in the GFP frame, altered GFP expression levels were observed as expected (Supplementary fig. 3b,c), and these sequences were therefore excluded from further analysis. Analysis of the remaining mutagenesis data revealed that the region from ∼40 nts upstream to ∼40 nts downstream of the start codon showed the greatest impact on start codon recognition efficiency, with observable effects up to 90 nts from the start codon (Fig. 2a and Supplementary Fig. 3d). To identify specific nucleotides that impact start codon recognition, we next mutated each nucleotide individually into all three other possible nucleotides within the region from -40 to +39 for 18 genes. This fine-grained scanning mutagenesis was performed for three different TISs to assess if the sequence of the TIS impacts the effect of distal sequences on start codon recognition. Mutation of numerous nucleotides outside the TIS caused robust changes in start codon recognition efficiency (Fig. 2b). To validate the impact of individual distal nucleotides on start codon recognition efficiency, we also computed the average contribution of each nucleotide to start codon recognition efficiency across 2000 endogenous mRNA sequences using two different TISs and two different 5’UTR lengths (See Methods). While the sensitivity of this second approach assay was somewhat limited due to the inherent high sequence complexity of large sequences, many of the preferred nucleotides were confirmed throughout all analyses, including most notably -40 to -27U; -14 to -9A; +6G/U; +11 to +13A; (Fig. 2b, Supplementary Fig. 4a-d). Validation experiments confirmed the importance of all four of these mRNA sequence regions (Supplementary Fig. 4e-k). We note that enrichment profiles downstream of nt +25 were not consistent between approaches, so we did not pursue further analysis of this region.

**Figure 2.**
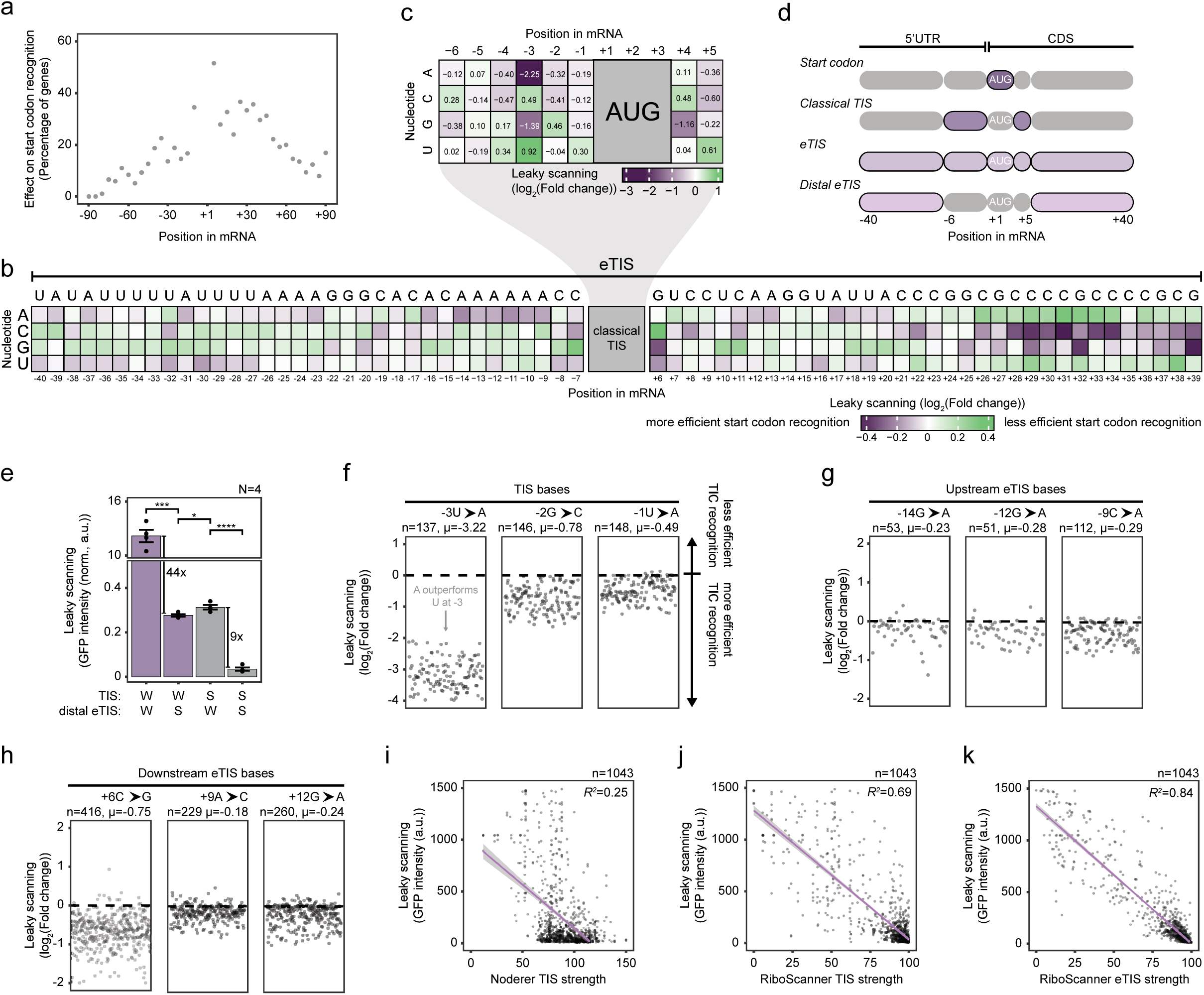
Start codon recognition is modulated by the eTIS. **a**, Scanning mutagenesis was performed on mRNAs with a 5’UTR length of 90 nts in which blocks of 5 consecutive nucleotides were mutated. The fraction of mRNAs in which a mutation caused a change in GFP expression of >1.25 fold is plotted for each position relative to the start codon. **b**, Heatmap representing the average log_2_(Fold Change) in GFP expression of the RiboScan reporter upon the introduction of the indicated nucleotide at the indicated position relative to the start codon. The sequence above the heatmap represents an example of a very strong eTIS. Note that in this strong eTIS introduction of (near-cognate) start codons was avoided to promote initiation at the main AUG start codon. **b**, **c**, Purple boxes indicate a reduction in GFP expression (enhanced start codon recognition), while green boxes indicate an increase in GFP expression (reduced start codon recognition). **d**, Schematic illustrating mRNA sequence regions that influence start codon recognition. The classical TIS comprises nucleotides −6 to +5 (excluding the start codon), whereas the eTIS refers to larger region of approximately -40 to +39. The term distal eTIS is used to refer to the eTIS region that flanks the classical TIS (-40 to -7 and +6 to +39). **e**, Quantification of GFP/mCherry ratios for RiboScan constructs containing combinations of strong (S) or weak (W) distal eTIS and TIS elements. Bars represent the mean GFP/mCherry ratio across four replicates, with individual replicates shown as dots. Fold changes in fluorescence ratios between sequences sharing the same TIS are indicated. Statistical significance was assessed using a Welch’s two-sample t-test. * denotes *P* < 0.05, *** *P* < 0.001, **** *P* < 0.0001. **f**-**h**, Log_2_(Fold Change) in GFP expression of the RiboScan reporter when comparing the optimal nucleotide with the least optimal nucleotide for indicated positions within the TIS (**f**) and distal eTIS (**g**-**h**). Each dot corresponds to an mRNA sequence carrying the specified substitution at the indicated position. **i**-**k**, GFP expression of the RiboScan reporter for 1043 mRNA sequences is plotted against TIS strength (Noderer *et al*. (Noderer *et al*., 2014)) (**i**) predicted TIS strength determined in this study (**j**) and predicted eTIS strength (**k**). Linear regression fits are shown in purple, with grey shaded regions indicating the 95% confidence intervals.

To determine the relative importance of the newly identified distal nucleotides to start codon recognition with that of the well-established classical TIS nucleotides spanning positions −6 to +5, we also assessed the effect on start codon recognition of individual mutations at each TIS position (Fig. 2c,d). Comparison of fold changes in GFP expression revealed that mutation of the -3 and +4 positions showed the largest change in GFP expression (Fig. 2c), consistent with the well-established importance of these two positions in start codon recognition (Hernandez *et al*., 2019; Kozak, 1986; Noderer *et al*., 2014). However, mutation of most other TIS nucleotides showed effect sizes that were in a similar range as mutation of many individual distal nucleotides (Fig. 2b,c). Furthermore, the summed positive effect of all distal nucleotides was even greater than that of the TIS nucleotides (11.63 vs. 5.18, respectively). Consistently, an mRNA encoding strong (S) distal nucleotides showed highly efficient start codon recognition even in the context of a weak (W) TIS. Notably, start codon recognition efficiency for this mRNA even exceeded that observed for an mRNA harboring a strong TIS and weak distal nucleotides (Fig. 2e). Together, these results identify key distal nucleotides that are important for start codon recognition and show that these distal nucleotides are at least as potent as TIS nucleotides in promoting start codon recognition. Since we find that nucleotides within the -40 to +39 region are potent modulators of start codon recognition, we refer to this region as the extended TIS (eTIS) and to the part of the eTIS that flanks the classical -6 to +5 TIS, as the ‘distal eTIS’ (Fig. 2d).

Distal eTIS nucleotides could affect start codon recognition through direct interactions with the ribosome and/or its associated translation factors, as was reported for several TIS nucleotides (Martin et al., 2016; Petrychenko et al., 2024; Simonetti et al., 2020; von Loeffelholz et al., 2025a). Alternatively, distal nucleotides could exert their impact on start codon recognition through interactions with RNA binding proteins (RBP) or through altering RNA structure. To discriminate between these possibilities, we examined the effect of each individual nucleotide substitution across all mRNA sequences. If distal nucleotides act through RBP interactions or RNA structure, the expectation is that the effect of mutations would be highly mRNA specific, as RBP binding motifs (which are typically 6-8 nts (Corley et al., 2020)) and RNA structures are dependent on a specific nucleotide flanking sequence. In contrast, if distal nucleotides act through direct interactions with the ribosome and its co-factors, mutations likely have the same effect on start codon recognition, irrespective of the surrounding mRNA sequence. We first examined TIS mutations, and found that TIS mutations had similar effects on start codon recognition, irrespective of mRNA sequence, as expected (Fig. 2e, Supplementary Fig. 4l). Analysis of distal eTIS nucleotides showed a very similar pattern to TIS nucleotides (Fig. 2f,g and Supplementary Fig. 4m-o), suggesting that distal nucleotides mainly act through direct interaction with the ribosome and its co-factors to impact start codon recognition efficiency.

### A machine learning model predicts start codon recognition

Since TIS strength does not correlate well with start codon recognition (See Fig. 1e), we wondered if the eTIS could predict start codon recognition more accurately. We therefore aimed to develop an eTIS strength score to test whether eTIS strength correlates well with start codon recognition efficiency. To this end, we trained a deep convolutional neural network (DCNN), termed RiboScanner, on 90% of our RiboScan dataset to predict GFP expression directly from mRNA sequence (Supplementary Fig. 5a). Based on the predicted GFP expression, we generated an eTIS score that ranged from 0-100, with higher values indicating more efficient start codon recognition. To determine whether the RiboScanner model could predict start codon recognition efficiency more accurately than the previous TIS model, we examined the 10% of the sequences withheld from training and validation. Indeed, the RiboScanner model dramatically improved prediction of start codon recognition efficiency (R^2^=0.25 vs 0.84 for the previous TIS and new eTIS models, respectively) (Fig 2h,i) (Note the slightly higher correlation for the previous TIS model (Noderer *et al*., 2014) compared to figure 1e, which is due to inclusion of non-native sequences in Fig. 2h-j, see Methods), indicating that the RiboScanner eTIS model had successfully identified many mRNA sequence features that determine start codon recognition.

Since our eTIS model includes analysis of both the TIS and distal eTIS regions, we asked whether improved start codon prediction of the eTIS model was due to inclusion of the distal eTIS in the analysis, due to improved prediction of the TIS, or both. We first tested whether the model had correctly learned the relative contributions of distal eTIS nucleotides for start codon recognition. For this, a set of reporters was created that contained a fixed TIS but variable distal eTISs. Comparison of the observed GFP values with those predicted by the RiboScanner model showed very good correlations for multiple reporter designs (R^2^=0.63-0.71) (Supplementary Fig. 5b). As the TIS was kept constant in these experiments, the high predictive power of the RiboScanner model demonstrates its ability to accurately interpret the effect of the distal eTIS on start codon recognition.

Next, we assessed whether the RiboScanner model also predicted TIS strength more accurately than the previous TIS model. For this, we generated a separate TIS score using the RiboScanner model (range 0-100, see Methods). Comparing TIS strength values from our model with those generated by the previously-developed TIS score (Noderer *et al*., 2014) showed a moderate correlation (R^2^=0.57), with both models emphasizing the importance of a -3 purine (Supplementary Fig. 5c-e). However, the RiboScanner TIS model was able to predict start codon recognition with substantially improved accuracy compared to the previous TIS model by Noderer and colleagues (R^2^ = 0.69 vs R^2^=0.25) (Fig. 2h,i and see Methods), suggesting that accurate prediction of start codon recognition by the eTIS models was also due to an improved interpretation of the classical TIS region. Importantly, the RiboScanner TIS model was not as good as the eTIS model in predicting start codon recognition (R^2^ = 0.69 vs R^2^=0.84), an effect that was especially notable for sequences with strong start codon recognition (Supplementary Fig. 5f,g), as is the case for most start codons present in endogenous genes. We conclude that both improved interpretation of the classical TIS region as well as inclusion of the distal eTIS allowed high accuracy prediction of start codon recognition by the RiboScanner model.

Since the RiboScanner eTIS model can predict start codon recognition efficiency on any mRNA sequence, we asked if it could aid in start codon identification in the human transcriptome. Using high quality ribosome profiling data (Mudge et al., 2022), we found that AUG start codons with strong experimental evidence of translation had significantly higher eTIS scores compared to AUG codons that showed weak evidence for translation (Supplementary Fig. 6a). Similar results were obtained when analyzing the near-cognate start codons CUG and GUG present in 5’UTRs (Supplementary Fig. 6b-e). These results show that eTIS strength correlates with transcriptome-wide start codon usage, further validating the eTIS model and suggesting that the eTIS model could help identify novel translation start codons, open reading frames and (micro-)proteins.

### Optimal eTIS residues are enriched and conserved

A key feature of the classical TIS is that favorable nucleotides (e.g. -3A or G) are enriched in the transcriptome (Hernandez *et al*., 2019; Kozak, 1986; 1987a), suggesting that evolution has shaped the sequence of this region for efficient start codon recognition. To determine if the distal eTIS has undergone similar evolutionary selection, we performed bioinformatic analysis of nucleotide enrichment in the distal eTIS regions. A complicating factor in this analysis is that GC content spikes at gene promoters (Zhang et al., 2004), which creates a gradient of increased GC content in 5’UTRs (Fig. 3a). As a consequence, 5’UTR length and presence of an intron in the 5’UTR impact GC content in the distal eTIS (Fig. 3a,b, Supplementary Fig. 7a,b). To overcome this challenge, we employed two parallel strategies; first, we normalized nucleotide abundance at each position relative to the start codon to local nucleotide abundance (See Methods). For nucleotides in the coding sequence, nucleotide abundance was additionally normalized to the position within the codon (i.e. position +6 was normalized to +9, +12, +15 etc.) to eliminate codon-based effects. In a parallel approach, we analyzed genes that encode an intron in the 5’UTR (at least 40 nts upstream of the start codon) that places the start codon beyond the GC spike originating from the promoter (See Fig. 3a, Supplementary Fig. 7b).

**Figure 3.**
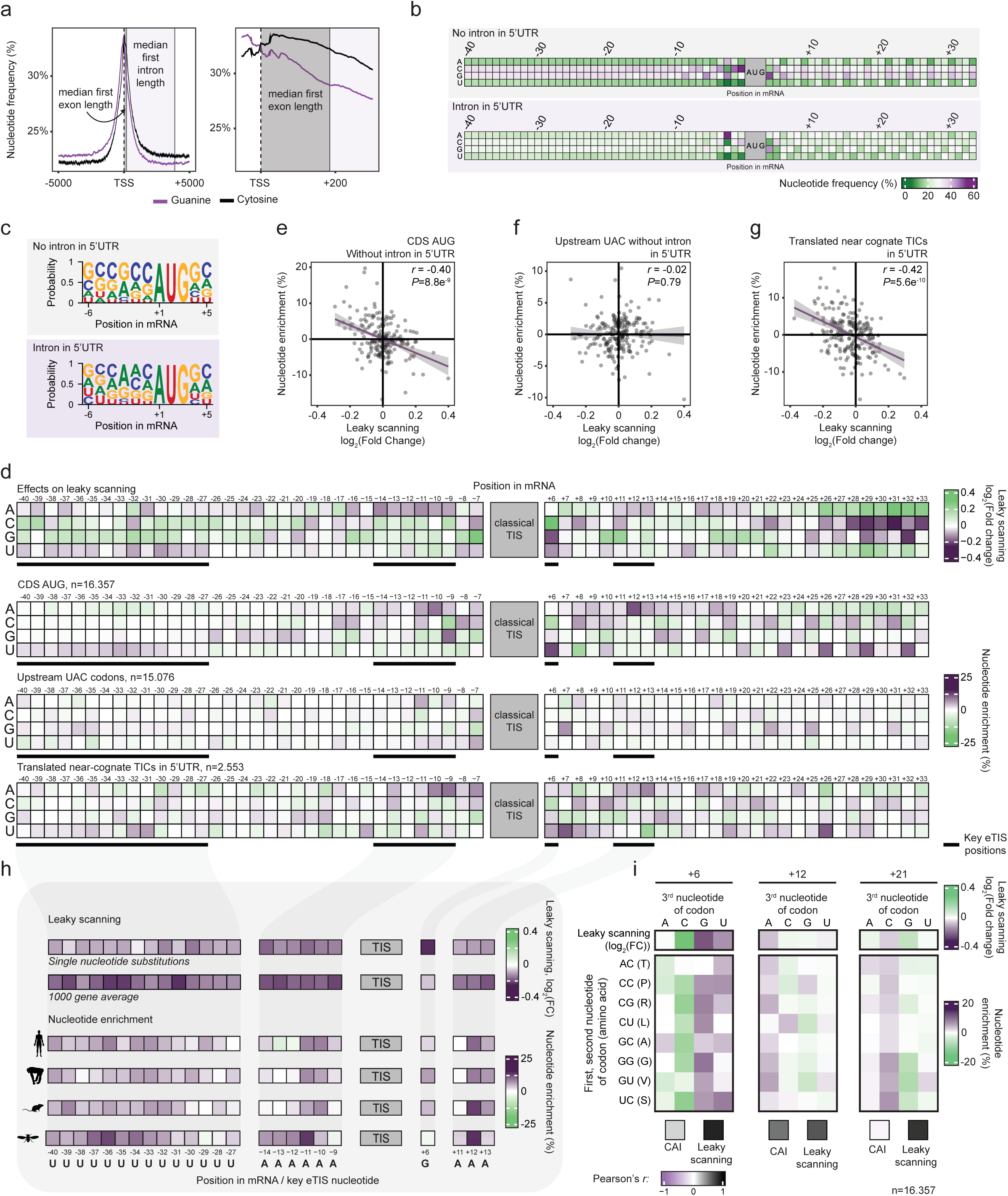
Genome-wide nucleotide enrichments in the eTIS. **a**, Genome-wide analysis of guanine and cytosine frequencies in the sense strand of the human genome, centered around the TSS. The average length of the first intron and exon is highlighted. Right graph shows a zoom-in of the region near the TSS. **b**, Heatmap representing the frequency of each nucleotide at indicated positions relative to the start codon. Nucleotide frequencies for genes without (Top) or with (Bottom) intron in the 5’UTR are shown. Note the difference in GC content. **c**, Nucleotide frequencies of the TIS for genes without (Top) or with (Bottom) intron in the 5’UTR. **d**, Heatmap representing the average log_2_(Fold Change) in GFP expression of the RiboScan reporter upon introduction of the indicated nucleotide at the indicated position relative to the start codon. Heatmap is replotted from Fig. 2b for comparison. Bottom three heatmaps show the enrichment of each nucleotide at indicated positions relative to the start codon. Different heatmaps represent data for canonical CDS AUG start codons (Top), upstream UAC codons as a negative control (middle) and upstream CUG or GUG start codons for which translation has experimentally been verified (Mudge *et al*., 2022) (Bottom). Key distal eTIS residues are underlined. **e-g** Comparison between the measured effect on leaky scanning of each nucleotide substitution and the genome-wide enrichment of the corresponding nucleotide. Data is shown for canonical CDS AUGs (**e**), upstream UAC codons as control (**f**), and translated upstream near-cognate CUG and GUG start codons (**g**). Linear regression fits are shown in purple, with grey shaded regions indicating the 95% confidence intervals. Pearson correlation coefficients (*r*) and corresponding *P* values are indicated for each panel to assess the significance of the linear relationships. **h**, Nucleotide sequences within the distal eTIS show consistent improvement of start codon recognition across experimental conditions, are preferentially utilized across the transcriptome and exhibit robust evolutionary conservation. Average log_2_(Fold changes) in leaky scanning (top three rows) upon the introduction of the indicated key distal eTIS residue at the indicated position relative to the start codon are replotted from Fig. 2b, and **Supplementary Fig. 4a,b,** respectively. Nucleotide enrichment of key distal eTIS residues (bottom four rows) were replotted from **Supplementary Fig. 8a**, for *H. sapiens*, *P. troglodytes*, *M. musculus* and *D. Melanogaster*, respectively. **i**, Heatmap depicting nucleotide enrichments for the third nucleotide of the codon (wobble base) for amino acids whose codons permit any of the four nucleotides at the wobble base. Position relative to the start codon is shown on top. The effect of each nucleotide on start codon recognition is shown in the top row (replotted from Fig. 2b). The two lower boxes report Pearson correlation coefficients between nucleotide enrichment and CAI (left), and between nucleotide enrichment and the effect on leaky scanning (right). Nucleotide enrichments correlate better with start codon recognition efficiency than with codon optimality.

Using these approaches, we first re-examined nucleotide enrichment within the classical TIS to ask if the Kozak sequence, which is highly GC-rich (GCC(^G^/_A_)CC**<u>AUG</u>**GC), is also affected by the promoter GC spike. When we examined nucleotide frequencies in the TIS for genes without an intron in the 5’UTR, the TIS consensus sequence was highly GC-rich and closely resembled the Kozak consensus sequence (Fig. 3c, top). In contrast, genes with an intron in the 5’UTR showed a TIS consensus that was more AU-rich (Fig. 3c, bottom, See Methods). Interestingly, when comparing the two TIS consensus sequences with our TIS mutagenesis data, we found that the AU-rich consensus sequence of 5’UTR intron-containing genes matched the *optimal* sequence for start codon recognition more closely (*R^2^*=0.71 vs *R^2^*=0.46). Specifically, a uridine at -5 and adenosines at -3, -2 and -1 were more strongly enriched in the intron-containing consensus sequences compared to intronless sequences, and these nucleotides performed very well in promoting start codon recognition in functional data (Fig. 2c). Of note, while both adenosine and cytosine at -1 and -2 positions are strong enhancers of start codon recognition, the optimal nucleotide at -1 depends on the nucleotide at the -2 position, and vice versa; with adenosine being optimal when the other position contains a purine (e.g. another adenosine), while a cytidine outperforming adenosine if the other position contained a pyrimidine (e.g. another cytidine) (Supplementary Fig. 7c,d). Such context-dependent strengths were only observed for the -1 and -2 positions within the TIS (Supplementary Fig. 7e). Taken together, these findings strongly suggest that the Kozak sequence was ‘polluted’ by the promoter GC spike, and does not represent the optimal TIS for start codon recognition. Rather, the optimal TIS is represented by the sequence GUCA(^AA^/_CC_)**<u>AUG</u>**GC, re-defining one of the most iconic and widely-used sequences in molecular biology and medicine.

Using the same approach as above, we found that distal eTIS nucleotides also showed strong enrichments for specific nucleotides at many positions (Fig. 3d, Supplementary Fig. 7f). Comparing these nucleotide enrichments to the functional effects of each nucleotide on start codon recognition revealed a significant correlation between the prevalence of a specific nucleotide and the impact of that nucleotide on start codon recognition (r = -0.40 for all transcripts and r = -0.36 for transcripts with an intron in the 5’UTR; note that correlations are negative as nucleotide enrichment was correlated with leaky scanning (Fig. 3e, and Supplementary Fig. 7g)). Notably, nucleotide sequences within the distal eTIS that showed consistent improvement of start codon recognition across experimental conditions often also showed strong enrichment in the genome (Fig. 3h). Key distal eTIS nucleotides that promote start codon recognition and are enriched include -40 to -27U; -11 to -9A; +6G/U; +11 to +13A. Other nucleotides (e.g. +26U) showed strong nucleotide enrichment, but lacked a corresponding increase in start codon recognition efficiency (Fig. 3d and Supplementary Fig. 7f), suggesting that these nucleotides may have experienced selective pressures other than start codon recognition. As a control, we also examined nucleotide enrichments at a non-start codon (UAC), which showed little global nucleotide enrichment and no significant correlation with start codon recognition (Fig. 3d, f, Supplementary Fig. 7h). A similar degree of nucleotide enrichment was observed surrounding near-cognate start codons (CUG and GUG) that showed strong experimental evidence of translation (Fig. 3d,g) (Mudge *et al*., 2022), but not for near-cognate start codons that showed no evidence of translation (Supplementary Fig. 7i), further confirming the importance of the eTIS in marking sites of translation initiation. Enrichment of specific nucleotides did not extend beyond the distal eTIS region, indicating that the eTIS region was uniquely relevant for translation (Supplementary Fig. 7j). Moreover, the key regions of the distal eTIS that showed strong activity and genome-wide enrichment, also showed robust evolutionary conservation (Fig. 3h, Supplementary Fig. 8a-d), further supporting their importance.

In addition to 5’UTRs, multiple nucleotides present within the coding sequence of genes, including the +6 position, showed genome-wide enrichment profiles. However, interpretation of nucleotide enrichment in the coding sequence is more complex, as these nucleotides are also under selective pressure from amino acid and codon usage biases (Hanson and Coller, 2018; Tuller et al., 2010; Verma et al., 2019). The +6 position is especially complex, as it may be under additional selective pressure from the fact that the N-terminal amino acid determines protein stability through the ‘N-end rule’ (Sriram et al., 2011). To disentangle amino acid- from nucleotide-based effects, we performed nucleotide-enrichment analyses specifically for the third nucleotide of the codon (wobble base) for amino acids whose codons permit any of the four nucleotides at the wobble base. We first zoomed-in on the first encoded amino acid (+4 to +6 nucleotide position). For almost all amino acids, a strong enrichment of the G or U—the optimal nucleotides for start codon recognition—was observed at the wobble position of this codon (+6 position) (Fig. 3i). A similar effect was observed for many other codons within the distal eTIS (Fig. 3i, Supplementary Fig. 7k), and these effects were conserved in many other metazoans (Supplementary Fig. 8e). Additionally, wobble base nucleotide enrichments correlated much better with start codon recognition efficiency than with codon optimality (as assessed by the codon adaptation index (CAI) (Sharp and Li, 1987) (Fig. 3i, Supplementary Fig. 7k, lower boxes), suggesting that optimal start codon recognition may have been a stronger selective pressure than codon optimality in shaping the 5’ region of coding sequences.

### eTIS optimization for mRNA therapeutics

We next aimed to leverage the discovery of the eTIS to optimize gene expression of a transgene. Most engineered transgenes, including those used in biomedical research and in clinical applications, are optimized for efficient start codon recognition using the Kozak sequence to increase expression levels and reduce ‘off-target’ protein production caused by downstream initiation. We reasoned that eTIS optimization could further improve transgene expression. To this end, we selected the nucleotide at each position that most strongly enhanced start codon recognition efficiency (See Fig. 2b). Introduction of an optimized eTIS resulted in an almost complete elimination of GFP expression in almost all endogenous mRNAs tested (Fig. 4a), demonstrating that eTIS optimization results in highly efficient start codon recognition. eTIS optimization even allowed highly efficient translation initiation on CUG or GUG start codons, an effect that was not observed with TIS optimization alone (Fig. 4b and Supplementary Fig. 9a), highlighting the potency of eTIS optimization in enhancing start codon recognition. Even for vaccine mRNAs which have already been extensively optimized, eTIS optimization substantially improved start codon recognition (Fig. 4c). Together, these results show that eTIS optimization results in near-perfect start codon recognition, eliminating off-target protein production caused by leaky scanning.

**Figure 4.**
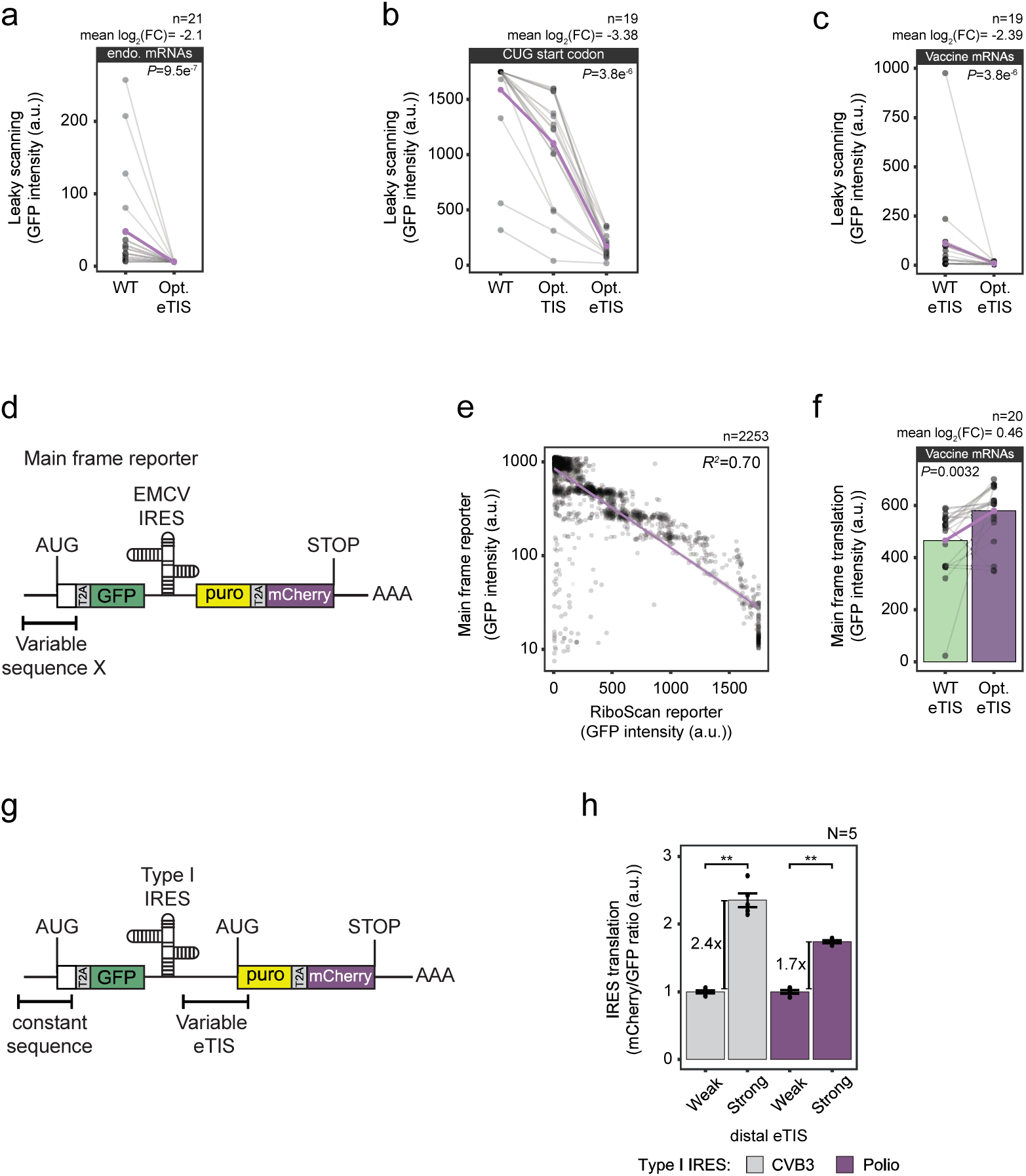
eTIS optimization for therapeutic mRNAs. **a**-**c**, Either the eTIS (**a**, **b**, **c**), or the TIS alone (**b**) was optimized for indicated mRNAs. The purple lines represent the mean changes in leaky scanning upon (e)TIS optimization. **d**, Schematic of the reporter used to assess the effects of the eTIS on expression levels of the protein encoded by the main ORF (GFP). Expression of mCherry was driven from the same mRNA downstream of an IRES to normalize GFP expression to mRNA levels. **e**, Main frame GFP expression plotted against RiboScan GFP expression (leaky scanning). A linear regression fit is shown in purple, with grey shaded region indicating the 95% confidence interval. **f**, the eTIS was optimized for vaccine mRNAs and main frame GFP expression was determined. The purple lines represent the mean changes in leaky scanning. *P* values in **a**, **b**, **c** and **f** were determined using a paired Wilcoxon signed-rank test. **g**, Schematic of the reporter used to assess the effects of the eTIS on IRES-driven translation. The expression of the puro-mCherry ORF is driven by a type I IRES in which the TIS is surrounded by strong or weak distal eTIS sequences. Expression of GFP was driven from the same mRNA to normalize mCherry expression to mRNA levels. **h**, Quantification of IRES-driven translation for constructs shown in **g**, containing weak or strong distal eTIS sequences. Data is represented as the mCherry/GFP fluorescence ratio. Bars represent the mean across five biological replicates (N = 5), with individual replicates shown as dots. Fold changes in fluorescence ratios between sequences sharing the same IRES are indicated. Statistical significance was assessed using a Welch’s two-sample t-test. ** denotes *P* < 0.01.

To determine whether improved start codon recognition upon eTIS optimization also resulted in increased expression of the main frame protein product, we engineered a reporter mRNA in which GFP was encoded in the main frame (frame 0). Expression of mCherry was driven by an IRES from the same mRNA for mRNA expression normalization (Fig 4d). We first compared GFP expression levels in this main frame GFP reporter to GFP expression from the RiboScan reporter for the same set of sequences, which revealed a strong inverse correlation between both values, indicating that efficient start codon recognition enhances main frame protein production, as expected (Fig. 4e). Consistent with this, eTIS optimization of vaccine mRNAs resulted in a significant increase in GFP expression in most cases (Fig. 4f). eTIS optimization also enhanced start codon recognition in Cos7 and CHO-K1 cells, cell types frequently used for recombinant protein production (Supplementary Fig. 9b,c), extending the potential applications of this approach.

Circular RNAs (circRNAs) are increasingly used in RNA-based therapeutics owing to their resistance to exonuclease-mediated degradation (Liu et al., 2022; Wesselhoeft et al., 2018). Translation of circRNAs typically relies on IRESs to recruit ribosomes to the RNA. To determine whether eTIS optimization also improves start codon recognition during IRES-driven circRNA translation, we measured protein output from mRNAs encoding type I IRESs—which are often used for therapeutic protein expression—with either a strong or weak distal eTIS (Fig. 4g). Strong distal eTIS sequences consistently increased total protein expression (Fig. 4h), indicating that distal eTIS residues promote start codon recognition in IRES-driven translation as well.

Taken together, these findings demonstrate that eTIS optimization improves translational precision and enhances transgene expression for therapeutic applications.

### Control of initiation kinetics by the eTIS

To uncover the mechanism by which the eTIS enhances start codon recognition efficiency, we employed a reconstituted *in vitro* system of human translation initiation, in which translation dynamics can be assessed by real-time single-molecule fluorescence imaging (Fig. 5a) (Grosely et al., 2025). In this system, recruitment of the small ribosomal subunit to the mRNA is visualized through fluorescent labeling of Met-tRNAi^Met^ within the eIF2-GTP-Met-tRNAi^Met^ ternary complex (which is part of the 43S pre-initiation complex). Initial start codon selection during mRNA scanning is marked by the release of fluorescently-labeled eIF1 from the scanning initiation complex, and late-stage initiation is assessed by following recruitment of fluorescent eIF5B (Fig. 5a) (Dever *et al*., 2023; Grosely *et al*., 2025). To specifically assess the effects of the distal eTIS residues on translation initiation kinetics, we designed two mRNAs with either a strong or weak distal eTIS, both containing the same weak TIS. As a control, a third mRNA was designed with a strong TIS and random distal eTIS (Fig. 5b, see Supplementary Table 1 for full mRNA sequences). All three sequences were first tested *in vivo* using the RiboScan reporter, which confirmed that distal eTIS optimization substantially reduced leaky scanning (Supplementary Fig. 10a). Start codon recognition was in a similar range for mRNAs containing a strong distal eTIS and weak TIS as for an mRNA containing a strong TIS and random eTIS, demonstrating that a strong distal eTIS can compensate for a weak TIS (Supplementary Fig. 10b). We then turned to the *in vitro* system to analyze translation initiation kinetics using the same mRNA sequences (Fig. 5c). First, we determined the fraction of ribosomes recruited to the mRNA that (eventually) progressed to late-stage initiation (eIF5B binding). Consistent with *in vivo* results, for both the strong TIS and strong distal eTIS mRNAs the large majority of ribosomes initiated translation during the experiment. In contrast, for the weak distal eTIS mRNA the majority of ribosomes failed to initiate (Fig. 5d). These results confirm the important role of the distal eTIS in start codon recognition, and further confirm that the distal eTIS functions independently of other trans-acting factors, such as RBPs.

**Figure 5.**
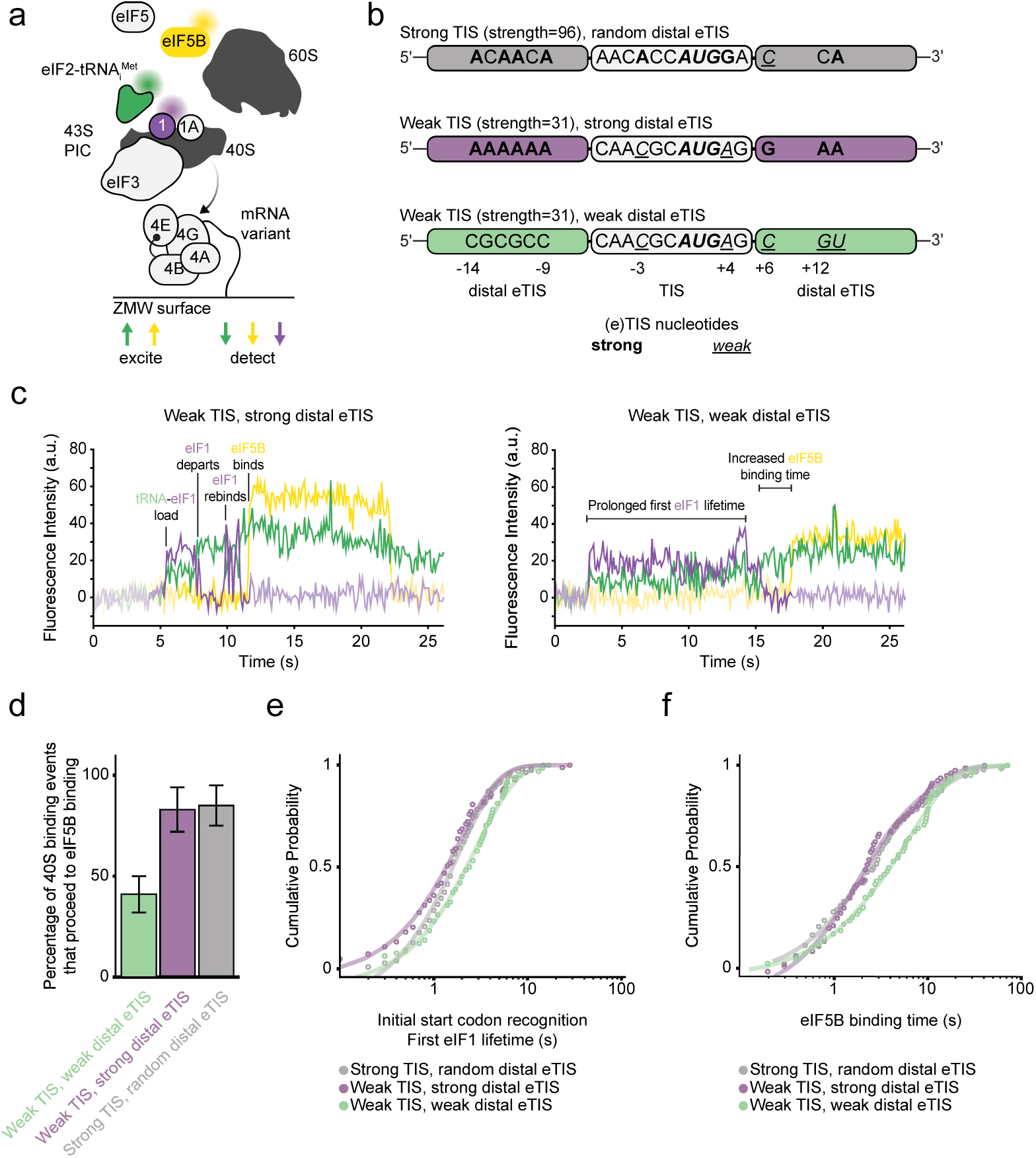
The eTIS modulates the kinetics of start codon recognition. **a**, Schematic of the *in vitro* single-molecule imaging approach (Grosely *et al*., 2025). mRNA variants are attached to the glass surface of zero-mode waveguides (ZMW). Proteins included in translation reaction are indicated, and fluorescently tagged proteins are indicated by colors. eIF1 (1) and eIF1A (1A) protein names are abbreviated. **b**, Schematic of mRNA sequences used in *in vitro* imaging experiments. (e)TIS residues that are either weak or strong are indicated. **c**, Example traces for mRNA with strong (left) and weak (right) distal eTIS nucleotides. Binding and departure of the 40S ribosome and individual translation factors is shown. Simultaneous appearance of tRNA and eIF1 indicates 40S binding. **d**, Percentage of 40S ribosome-mRNA binding events that result in eIF5B binding for indicated mRNAs. The error bars represent the 95% confidence interval derived from binomial bootstrapping of the observed events. **e**, Time from 40S recruitment to initial departure of eIF1 is show, which represent the time until initial start codon recognition. **f**, Time from the final eIF1 departure to eIF5B binding.

To assess if translation initiation dynamics were affected by the distal eTIS, we next zoomed into the subset of initiation events in which eIF5B was eventually recruited. We first examined the time from 43S mRNA recruitment to eIF1 release (‘first eIF1 lifetime’), which reflects the time required for initial start codon recognition. mRNAs containing a strong distal eTIS and a weak TIS showed similar initial start codon recognition kinetics as mRNAs with a strong TIS. In contrast, initial start codon recognition kinetics was 2-fold slower on mRNAs with a weak distal eTIS, indicating that the distal eTIS facilitates rapid start codon recognition (Fig. 5c,e). Previous work has shown that upon initial eIF1 release, eIF1 can rebind and release multiple times as the 48S translation initiation complex transitions back and forth between open and closed states, before committing to initiation (Grosely *et al*., 2025). The time to subsequent eIF1 release, as well as the overall time eIF1 could bind were increased for mRNAs with a weak distal eTIS (Supplementary Fig. 10c-e), indicating that ribosomes struggled to commit to initiation when the distal eTIS sequence was weak. Somewhat surprisingly, the time from final eIF1 release to eIF5B binding was also longer for the mRNA with weak distal nucleotides, indicating that even the steps in translation initiation post-start codon recognition depend on a strong distal eTIS (Fig. 5c,f). Together, these results indicate that initial start codon recognition, as well as subsequent steps in the initiation process are stimulated by the distal eTIS.

### Molecular basis of eTIS function

To understand how eTIS residues enhance initiation efficiency, we performed cryo-EM of *in vitro*–reconstituted human 48S translation initiation complexes (Petrychenko *et al*., 2024; Yi et al., 2022) assembled on two mRNAs that share an identical TIS with an optimal -3A and +4G, but have either a weak or strong distal eTIS sequence (Fig. 6, Supplementary Figs. 11, 12, Supplementary Tables 8, 9 and Methods). As expected, *in vivo* analysis of these two mRNA designs revealed that optimization of distal eTIS regions greatly improved start codon recognition efficiency (Fig. 6b,c).

**Figure 6.**
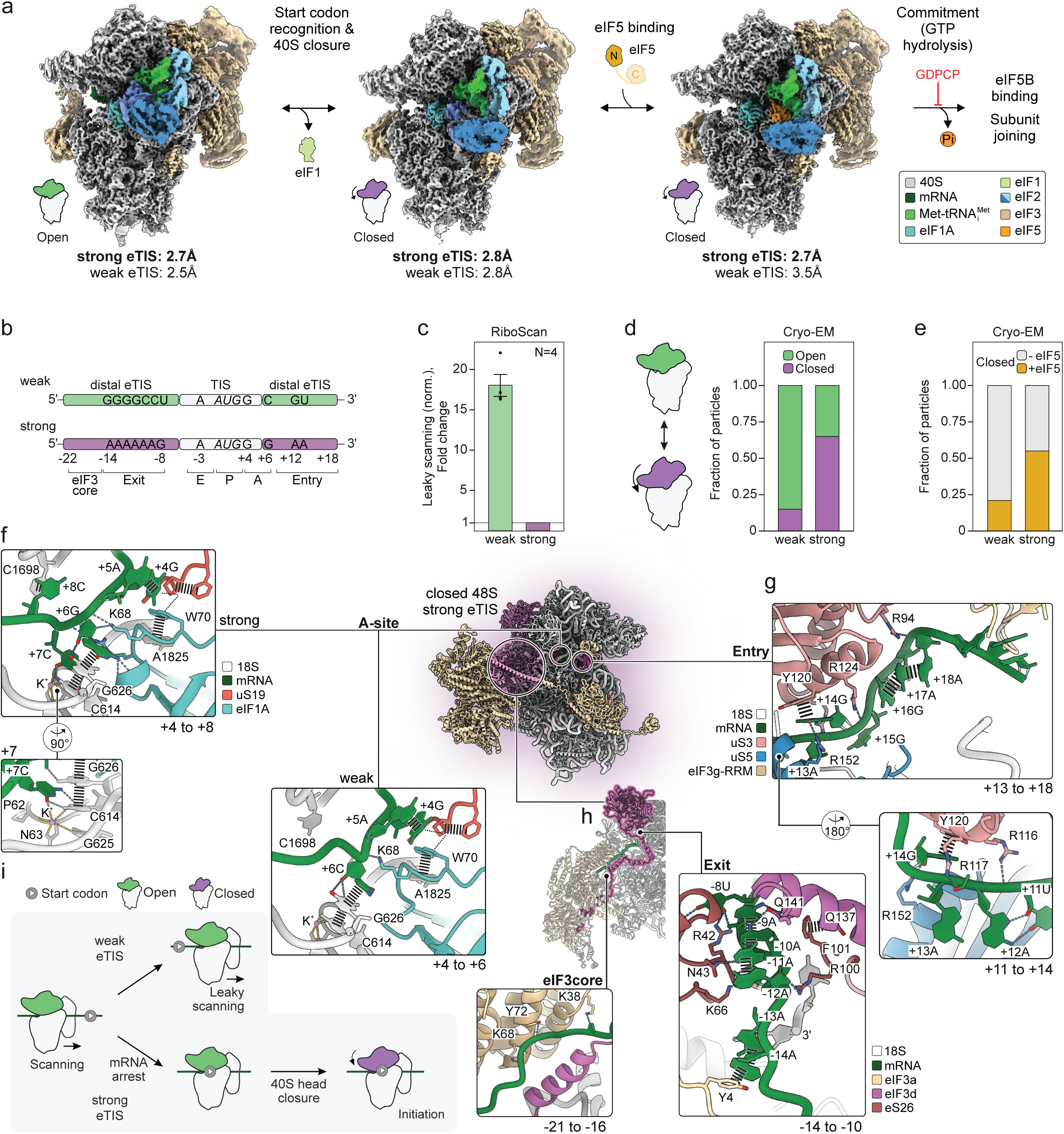
Distal eTIS residues promote initiation by stabilizing the 48S at the start codon and facilitating 40S head closure. Unless indicated otherwise, panels show 48S complexes assembled on mRNA with a strong distal eTIS. **a**, Cryo-EM analysis of 48S complexes assembled on distinct distal eTISs. Densities for strong distal eTIS were rendered using LocScale (Bharadwaj *et al*., 2025) with overall resolutions indicated in Å (bold). Resolutions for weak distal eTIS complexes are shown below. **b**, Schematics of mRNAs with identical TIS and distinct distal eTIS residues used for cryo-EM. Distal eTIS residues were designed for minimal (weak) or maximal (strong) start codon recognition (Methods). Key residues and interaction sites are indicated. **c**, Leaky scanning measured *in vivo* by RiboScan reporter in HEK293 cells, normalized to the strong distal eTIS construct. **d**, **e**, Effect of distal eTIS on 40S head closure (**d**) and eIF5 recruitment (**e**) as quantified by cryo-EM. eIF5 is the GTPase-activating protein of eIF2, promoting irreversible start codon commitment. **f**, Interactions of the key +6 residue in the A site of the decoding center for strong distal eTIS (left) and weak distal eTIS (bottom right). The extensive +6G network is further stabilized by +7C (close-up) and +8C. **g**, Distal eTIS interactions with uS3, uS5 and eIF3g-RRM at the mRNA entry channel. Close-up shows the interaction network of conserved nucleotides +12A and +13A, including uS3 residues R116 and R117. **h**, Distal eTIS interactions at the mRNA exit channel and eIF3 core. At the exit (bottom right), ribosomal protein uS26, 3’terminus of 18S rRNA (18S) and eIF3d-NTD clamp distal eTIS residues -9 to -14. Note the α-helix formed by eIF3d residues 125-142, typically unresolved in 48S structures. At the eIF3 core (bottom left), eIF3a interacts with the backbone of distal eTIS residues. **i**, Model for eTIS function in translation initiation. A weak eTIS disfavors start codon recognition, resulting in increased leaky scanning of the start codon and potential downstream initiation. In contrast, a strong eTIS promotes start codon recognition by stabilizing the scanning 48S complex at the start codon and facilitating 40S head closure.

Complexes were prepared for cryo-EM under steady-state equilibrium conditions in the presence of non-hydrolysable GDPPCP to prevent irreversible commitment to initiation. This allowed us to visualize how the different eTIS regions influence the critical checkpoints in start codon selection, in particular the transition of the 48S from the scanning-competent open state to the scanning-arrested closed state (Brito Querido et al., 2020; Llacer et al., 2015; Petrychenko *et al*., 2024). For each mRNA, we obtained three structures at 2.7–3.2 Å resolution capturing: (i) the scanning open 48S, (ii) 40S head closure upon start codon recognition and release of the fidelity factor eIF1 (Cheung et al., 2007; Hinnebusch, 2014), and (iii) subsequent recruitment of eIF5 to the closed 48S (Llacer et al., 2018) (Fig. 6a, Supplementary Fig. 11 and Methods).

We first quantified cryo-EM particle distributions among 48S states to assess distal eTIS-driven 48S dynamics and function of distal eTIS nucleotides. In line with efficient start codon recognition *in vivo*, the strong distal eTIS strongly promoted 40S head closure and eIF5 recruitment relative to the mRNA with the weak distal eTIS (Fig. 6c-e), indicating that eTIS elements increase initiation efficiency by promoting head closure and eIF5 binding. This is consistent with our single-molecule data showing accelerated eIF1 release, a prerequisite for head closure and eIF5 binding, with strong distal eTIS.

#### Key +6G stabilizes 48S at the start codon

Our high-resolution structures of the closed 48S complexes, trapped at the AUG start codon, provide insight into the molecular basis of these distal eTIS effects. In contrast, in the open 48S state the mRNA is highly flexible; despite extensive sorting trials the AUG start codon was not clearly discernable in the ribosomal P-site, indicating that these structures represent a mixture of scanning 48S complexes positioned at different locations along the mRNA. In the closed state, however, the AUG codon and surrounding mRNA are well resolved in the decoding center, enabling detailed structural analysis, and we therefore focused on the closed 48S structures.

In the closed 48S assembled on mRNA with a strong distal eTIS, the optimal +6G establishes an extensive interaction network in the A site of the 40S decoding center (Fig. 6f, left panels and Supplementary Fig. 12a bottom). The +6G engages the universally conserved G626 of 18S ribosomal RNA (rRNA) and, unexpectedly, eIF1A, a factor involved in initiation fidelity (Hinnebusch, 2017; Petrychenko *et al*., 2024). Specifically, the +6G base forms guanosine-specific hydrogen bonds with the backbone carbonyls of eIF1A residues K64 and R66; residues known to be important for efficient initiation (Battiste et al., 2000), but the underlying mechanism had remained elusive. In addition, the +6G backbone interacts with eIF1A K68 through hydrogen bonds involving the ribose and a salt bridge at the phosphate group, while its 2′-OH forms a hydrogen bond with the G626 carbonyl. Together, these contacts stabilize optimal sandwich π–π stacking of +6G onto the G626 base.

In contrast, in mRNA with a weak distal eTIS, the non-optimal +6C shows weaker stacking on G626, forming only limited non-specific interactions, primarily involving the phosphate with eIF1A K64 (Fig. 6f, lower panel and Supplementary Fig. 12a top). A non-optimal adenosine at +6 adopts distinct context-dependent configurations in the closed 48S: In a previous structure containing a +5U and +6A, the +6A stacks onto the Kozak residue +4G (Petrychenko *et al*., 2024). In a +5A context, +6A has been reported to participate instead in a stacking array extending from mRNA positions +4 to +7, but does not engage in an equivalent interaction network (von Loeffelholz et al., 2025b). These observations suggest a structural rationale for the hierarchy +6G > +6A > +6C in stabilizing mRNA at the start codon and promoting translation initiation, as reflected by *in vivo* RiboScan reporter measurements (Fig. 2b).

Downstream eTIS nucleotides may further reinforce the interaction network at the decoding site. In the weak eTIS mRNA, the +7A and +8A are poorly resolved, suggesting they are highly flexible. In contrast, +7C and +8C in the strong eTIS mRNA are well-resolved and extend the +6-interaction network (Fig. 6f). +7C interacts with a K^+^ ion and C614 of 18S rRNA, whereas +8C stacks onto C1698 of 18S rRNA. The +7C–C614 interaction is consistent with a non-canonical protonated C⁺–C base pair, potentially stabilized by the K^+^ ion.

Together, these data suggest that the +6 position may play a major and previously underappreciated role in stabilizing the scanning 48S at the start codon, consistent with our functional data identifying position +6 as an important determinant of start codon recognition efficiency (Fig. 2b, h, 3d and Supplementary Fig. 4e, n).

#### Distal eTIS interactions at the 48S periphery

Beyond the decoding center, distal eTIS residues establish extensive contacts with 40S ribosomal proteins and eIF3 subunits a, d and g at the 48S periphery. For the strong distal eTIS mRNA, focused classification of the cryo-EM data resolved these dynamic interactions at side-chain to secondary-structure detail, enabling assignment of the involved residues (Fig. 6g, h, Supplementary Figs. 11a, 12b-e and Methods).

At the mRNA entry channel (‘entry’), eTIS positions +19 to +22 interact with the RNA-recognition motif (RRM) of eIF3g, consistent with eIF3g-RRM’s general role in promoting initiation (Cuchalova et al., 2010). Conserved arginine residues of uS3 (R94, R116, R117 and R124) and uS5 (R152) contact the mRNA backbone of the purine stretch spanning from +12 to +18. Directly at the entry site, uS3 residue Y120 stacks onto +14G, while R116 and R117 coordinate nucleotides +12 and +13 (Fig. 6g, bottom right close-up). The optimal adenosines at +12 and +13 are highly enriched in the human transcriptome and broadly conserved across evolution (Fig. 3d and Supplementary Fig. 8a, e). Consistent with a functional role for these interactions, deletion of uS3 residues R116 and R117 alters start codon selection (Dong et al., 2017). In contrast, in the weak distal eTIS mRNA, residues in this region are poorly resolved already downstream of +6, impeding register assignment and detailed interpretation, and suggesting that optimal eTIS residues may reduce mRNA flexibility through stabilizing interactions with the 48S initiation complex.

At the mRNA exit channel (‘exit’), the poly-adenosine stretch (-14 to -9) of the strong distal eTIS mRNA forms two stacking arrays in the closed 48S that engage ribosomal protein eS26, the 3’ terminus of 18S rRNA and eIF3 subunits a and d (Fig. 6h, right panel, and Supplementary Fig. 12e). Nucleotides -14A and -13A stack onto eIF3a Y4, whereas -12A to - 9A are clamped between eS26, the 18S rRNA terminus, and the N-terminal domain (NTD) of eIF3d. eS26 residues I41, K66, R100 form non-specific contacts with the mRNA backbone at positions -11 and -10, while N43 directly engages the -11A base and R42 contacts the -8U base; Q113 of uS11 further recognizes uridine-specific features of the optimal -8U base (Supplementary Fig. 12e, top right). Collectively, these interactions stabilize the -9 to -12 stack for engagement by eIF3d. Remarkably, residues 125-145 of eIF3d–unresolved in previous structures–adopt an α-helical conformation in the closed state, spanning the mRNA and engaging the adenosine stack, with Q141 positioned to recognize adenosine-specific features of nucleotides -9 and -10 (Fig. 6h, bottom right, and Supplementary Fig. 12e). Recognition of the adenosine-rich sequences and -8U at the exit site suggests that optimal distal eTIS elements promote initiation not only by stabilizing the decoding center but also by specifically reinforcing mRNA interactions at the ribosomal periphery. Furthermore, these observations suggest a structural basis for how the eIF3d-NTD may contribute to start site selection. Previous functional studies implicated the eIF3d-NTD in start codon recognition at near-cognate start codons through an unknown mechanism (She et al., 2023). Our data indicate that eIF3d-NTD may modulate start codon recognition by engaging eTIS nucleotides at the mRNA exit channel. Finally, within the eIF3 core region upstream of the mRNA exit channel, additional contacts of eTIS residues ∼-20 to -17 with eIF3a and the NTD of eIF3d, including eIF3a sidechain interactions with the mRNA backbone (Fig. 6h, bottom left panel, and Supplementary Fig. 12d), suggest further contributions to mRNA stabilization at the periphery of the 48S complex.

In summary, our structural data reveal that distal eTIS residues form an interaction network across the entire 48S complex, extending from the eIF3 core region to the mRNA exit channel, over the decoding center to the mRNA entry channel. These interactions promote start codon recognition by anchoring the scanning 48S at the start codon and facilitating the structural rearrangements required for initiation commitment (Fig. 6i). Moreover, these findings provide a mechanistic framework for understanding previously identified roles of ribosomal proteins and initiation factors–in particular the enigmatic initiation factor eIF3d–in start codon recognition, linking their functional contributions to sequence-specific engagement of distal eTIS elements.

## Discussion

The position of start codons in the transcriptome determines proteome composition. However, many potential start codons exist in each mRNA and it has remained unclear which mRNA features determine which start codons are abundantly used for translation. Here, we identify a ∼80-nt sequence region extending across both the 5’UTR and coding sequence, which we term the extended TIS (eTIS), and show that the eTIS determines translation initiation efficiency at start codons.

### Mechanism of the eTIS

Our combined *in vivo* and *in vitro* biophysical and structural work establishes the eTIS as an mRNA-encoded regulatory module that operates across the full footprint of the scanning 48S complex. The eTIS engages the initiation machinery in a spatially coordinated manner along the mRNA trajectory, integrating interactions at the decoding center with mRNA engagement at the ribosomal periphery. At the decoding center, our structural and functional data suggest a major and previously underappreciated role for the +6 position, with additional roles of the +7 and +8 positions, in stabilizing the scanning 48S at the start codon. More distal elements include conserved poly-adenosine tracts at the mRNA entry (+12 to +18) and exit channel (-14 to -9) that form stacking arrays, stabilizing the mRNA along the dynamic 48S periphery and supporting both mRNA sequence-specific contacts and backbone interactions. Notably, only a subset of optimal eTIS residues engages in direct, nucleotide-specific interactions with the 48S initiation complex. This suggests that the eTIS functions through a distributed network of interactions, in which some optimal residues make direct nucleotide-specific contacts whereas others contribute more indirectly by stabilizing mRNA conformation and facilitating productive engagement. In addition, transient interactions, particularly in the scanning open 48S, likely contribute to initiation but are not readily captured by structural approaches.

In our 48S structure with strong eTIS mRNA, we resolve full-length eIF3d at side-chain to secondary structure resolution, revealing a basic α-helix that, together with eS26, engages the adenosine stack within the optimal eTIS at the mRNA exit channel. These findings suggest an explanation for the previously enigmatic role of eIF3d in start site selection and establish eIF3d as part of a coordinated eTIS recognition network that includes ribosomal proteins, 18S rRNA, and other initiation factors, including eIF1A and eIF3 subunits a and g. Together, our data support a model in which start codon selection emerges from a broad mRNA interaction landscape spanning the entire footprint of the 48S, with start codon recognition arising from cumulative contacts that anchor the scanning ribosome and promote commitment to initiation.

### Transcriptome-wide identification of start codons

With the advent of genome-wide methods for mapping translation initiation sites, it has become clear that most mRNAs contain multiple start codons, which drive expression of alternative protein isoforms and microproteins, and act as important translation regulators. However, identification of the complete set of start codons within the transcriptome has remained challenging, especially for start codons that are infrequently used for translation. Here, we introduce RiboScanner, a convolutional neural network model that improves start codon predictions compared to a previous model using TIS scores alone. We show that start codons with high eTIS scores are translated more heavily, and that favorable eTIS nucleotides are enriched around near-cognate start codons, suggesting that the eTIS signature can aid in identification of alternative start codons.

A complicating factor in sequence-based identification of start codons is that multiple types of evolutionary pressure, including the eTIS, promoters, amino acid preferences and codon usage, may have impacted mRNA sequence evolution near the start codon. As such, it is important to carefully deconvolve different mRNA features. For example, a previous study found that codon optimality was altered directly downstream of the start codon compared to the rest of the coding sequence (Tuller *et al*., 2010). Codon optimality scores may have been affected by evolutionary pressure to optimize the eTIS, and vice versa. Similarly, we show that a GC spike originating from promoters impacts the TIS and eTIS sequence composition, and that this GC spike affects the TIS consensus sequence. As such, the GC-rich Kozak sequence may represent the most *common* sequence among human genes, but likely not the most *optimal* sequence, which we show is more AU-rich. It therefore may be useful to replace the Kozak sequence with the optimal sequence GUCA(^AA^/_CC_)**<u>AUG</u>**GC in future mRNA designs. More generally, deconvolving the different process that have shaped mRNA sequence will be critical to better understand how translation initiation is regulated.

### Leveraging the eTIS for clinical application

The Covid-19 pandemic has highlighted the power of mRNA therapeutics and has resulted in an extensive effort to optimize mRNA sequence for improved translation (Metkar et al., 2024). Typically, high and specific expression of the encoded therapeutic protein is desired, both for *in vitro* and *in vivo* production of therapeutic proteins. We show that optimization of the eTIS not only increases protein expression yield, even in cases where the mRNA had already by extensively optimized (Fig. 4f), but also reduces the production of ‘off-target’ proteins (Fig. 4c, Supplementary Fig. 9b, c), caused by leaky scanning and downstream initiation, an important aspect that is often overlooked. Similarly, eTIS optimization will likely further improve production of recombinant proteins including antibodies for therapeutic use. The eTIS may therefore be useful for future optimization of clinical protein expression.

## Methods

### Cell lines

Human HEK293T, HeLa cells, and African green monkey kidney fibroblast-like cells (COS7) were cultured in DMEM (4.5 g/L glucose, GIBCO). Chinese Hamster Ovary cells (CHO-K1) were cultured in F-12 (GIBCO). Cells were supplemented with 5% (HEK293T) or 10% (CHO-K1, COS7, HeLa) fetal bovine serum (Sigma-Aldrich) and 1% penicillin/streptomycin (GIBCO) at 37°C and 5% CO_2_. Cell lines used in this study were routinely tested for the presence of mycoplasma.

### Lentiviral transduction

The various reporter libraries were introduced into cells using lentiviral transduction. For this, second generation, pHR-based lentiviral plasmids containing the various libraries were transfected into HEK293T cells together with pMD2.G and psPAX2 helper plasmids using PEI transfection (Polyethylenimine, Polysciences Inc.). One day after transfection, the medium was replaced by fresh cell culture medium. After 4 days, virus-containing supernatant was collected and passaged through a 0.33 μm filter to remove cellular debris and stored at -20°C. For lentiviral transduction, the virus-containing supernatant was added to HEK293T, CHO-K1, COS7 or HeLa cells. Six hours after virus addition, the cells were washed twice with PBS and fresh medium was added. Forty-eight hours after transduction, approximately 2% of the cells was set aside and maintained in puromycin-free medium to preserve an unselected population for MOI determination. The remaining 98% of cells were cultured under selective conditions by replacing the medium with fresh medium containing puromycin (final concentration: 1 µg/mL, ThermoFisher). A MOI was selected to achieve infection in approximately 10% of the cells, thereby minimizing the likelihood of individual cells expressing more than one reporter. After a minimum of one week of puromycin selection, cells were sorted by FACS.

### Design of the RiboScan reporters

The RiboScan reporter mRNA contained an test sequence (X), which typically included the full length 5’UTR sequence, an AUG start codon and a coding sequence of variable length (18 to 93 nts), followed by a GS linker, a single-nucleotide, which positioned the downstream GFP sequence out of frame with the start codon, and a T2A element (Liu *et al*., 2017), separating upstream translation products from GFP. Use of the T2A sequence ensured that GFP expression was not affected by the inserted coding sequence of the test sequence, which may affect protein stability. A stop codon was inserted downstream of the T2A to terminate all ongoing translation in the +1 frame (F+1) that was started at alternative translation initiation codons in the test sequence. Downstream of this stop codon, three tandem start codons with flanking Kozak consensus sequences (GCCACC***AUG***GC, x3) were introduced in the GFP reading frame to ‘capture’ all ribosomes undergoing leaky scanning of the main frame start codon. A stopless GFP-NLS sequence was introduced downstream of the three tandem AUG start codons, followed by a stop codon in the GFP frame. After the stop codon, a T2A–mCherry-NLS–T2A–puromycin resistance gene was inserted in the main frame (F0) to measure translation initiation at the main frame AUG (mCherry-NLS) and to select for cells expressing the reporter (puromycin resistance gene). See Supplementary Table 1, tab 2, for the fully annotated RiboScan reporter sequence. Where indicated (Fig. 2a, Supplementary Fig. 1j, 3b-d), the stop codon immediately upstream of GFP was absent in the RiboScan reporter. A CMV promoter was used to drive the expression of the RiboScan reporter, as the expression from this promoter is high in HEK293T cells and the TSS of this promoter had previously been identified (Isomura et al., 2008). The RiboScan reporter was cloned into a second-generation lentiviral vector (pHR). To enhance mRNA stability and facilitate nuclear export, the reporter construct was equipped with a woodchuck hepatitis virus post-transcriptional regulatory element (WPRE) (Zufferey et al., 1999). Polyadenylation was stimulated by the polyadenylation signal located in the 3′ long terminal repeat (LTR) of the lentiviral backbone. For the introduction of the different test sequences into the RiboScan reporter, two type IIS restriction sites (PaqCI/AarI, NEB) were introduced, one downstream of the CMV promoter and one upstream of the GS linker in the reporter. Type IIS restriction enzymes cut outside of their target sequence and thereby allow for scar-less integration of the test sequences. In this way, the 5’end of RiboScan mRNAs could be precisely controlled.

To monitor translation efficiency from the main start codon, a second reporter construct was generated (Main frame reporter). In this design, the GFP coding sequence was placed in frame with the main frame start codon and separated from the coding sequence of the inserted test sequence by a T2A sequence. Downstream of the GFP open reading frame (ORF), an additional ORF was introduced encoding a puromycin resistance gene and a second fluorescent protein (mCherry). The expression of the puro-mCherry ORF was driven by an EMCV internal ribosome entry site (IRES) to ensure its expression was independent of the main frame start codon driving expression of the test sequence and GFP. See Supplementary Table 1, tab 3 for the fully annotated main frame reporter sequence.

### Analysis of start codon recognition efficiency on endogenous mRNAs using the RiboScan reporter

#### Gene selection

To select genes for endogenous tagging with the RiboScan reporter, we applied the following criteria: 1. The gene should be highly expressed in HEK293T cells. 2. The gene should have a highly abundant transcript isoform. Transcript isoform expression data was obtained from the human Protein Atlas (Uhlen et al., 2005) and from Floor *et al*. (Floor and Doudna, 2016). 3. The transcript should have no AUG start codon other than the canonical start codon prior to the site of reporter integration

#### Production of the dsDNA repair template

The RiboScan reporter was introduced into the genome of HEK293T cells using CRISPR-Cas9-mediated homologous recombination. Double-stranded DNA (dsDNA) donor templates encoding the RiboScan reporter were generated by PCR amplification from the RiboScan reporter plasmid. Forward primers contained 75 nucleotide long homology arms corresponding to the target genomic locus and 15 nucleotides complementary to the sequence of the GS linker in the reporter plasmid (5′-GGCGGCAGCGGCTCA-3′). Reverse primers contained a homology arm of 70 nucleotides and 20 nucleotides complementarity to the reporter plasmid (5’-TGGGCCAGGATTCTCCTCGA-3’). A complete list of primers used to generate dsDNA repair templates is provided in Supplementary Table 1 (Tab 4). The linear dsDNA repair templates were excised from gel and purified with a DNA gel extraction kit (GeneJet, ThermoFisher). All dsDNA templates were eluted in nuclease free water.

#### gRNA design and synthesis

gRNAs were designed with CRISPick (Doench et al., 2016; Sanson et al., 2018) such that after correct integration of the reporter, the gRNA homology sequence is absent from the targeted locus. gRNAs were cloned as described in Hsu *et al*. (Hsu et al., 2013), using the PX459 plasmid (Addgene; #62988). The puromycin resistance cassette was removed from the plasmid by digestion with EcoRI (NEB) since the RiboScan reporter already contains a puromycin resistance gene for selection purposes.

Cells were transfected with 1µg of Cas9-guide construct and 3ug of linear donor template per 2 mL culture volume using FuGENE® HD (Promega), according to manufacturer’s protocol. 24 h after transfection cells were washed once with PBS and fresh medium was added. After 48 h, cells were treated with puromycin (final concentration: 1 µg/ml ThermoFisher). Puromycin-resistant cells were expanded until ready for analysis by microscopy. Correct integration of the RiboScan reporter at the targeted locus was verified using Sanger sequencing (Supplementary Fig. 1b, see Supplementary Table 1, tab 4, for applied primers).

#### Microscopy

Images were acquired using a Nikon TI inverted microscope with a perfect focus system equipped with a Yokagawa CSU-X1 spinning disc, a 100x, 1.49 NA oil immersion objective and an iXon Ultra 897 EM-CCD camera (Andor) using NIS elements (Nikon). The microscope was equipped with a temperature-controlled incubator and imaging was performed at 37°C. Cells were seeded in glass bottom 96-wells plates (Matriplates, Brooks) at a 20% confluency, two days prior to imaging. Prior to imaging, the medium was replaced with pre-warmed imaging medium (CO2-independent Leibovitz’s-15 medium (Gibco)) containing 5% fetal bovine serum (Sigma-Aldrich), 1% penicillin/streptomycin (Gibco) and 1:2000 Hoechst 33342 nuclear stain (ThermoFisher). Images were acquired with a 500ms exposure time for the 488 and 561 nm lasers and a 100ms exposure time for the 405 laser. A single Z-plane was imaged. For all imaging experiments, random non-overlapping positions in the imaging well were selected.

#### Image analysis

CellPose (v2.2.3) (Stringer et al., 2021) was used to perform nuclear segmentation based on the Hoechst signal. The resulting segmentation masks were employed to quantify nuclear GFP and mCherry intensities for each cell. Nuclei partially outside the field of view were excluded from further analysis. For each cell, the GFP to mCherry ratio was calculated to determine the efficiency of start codon recognition.

### Massively parallel reporter assay

#### Test sequence selection

Sequences of endogenous mRNAs were obtained from Ensembl (Release V110, July 2023). All transcripts with a Refseq ID and a 5’UTR length between 6 and 122 nucleotides were considered as input for the sequence libraries and a subset of these transcripts were randomly sampled. The complete 5’UTR and the first 18, 33, 39 or 93 nucleotides of the CDS of the transcripts were downloaded using BiomaRt (Smedley et al., 2009). In case multiple transcript isoforms were selected per gene, each of the transcripts was uniquely labeled by adding a unique number to the gene name (indicated as Gene_name.Number in the library). SNP information was obtained from the International Cancer Genomics Consortium (“simple_somatic_mutation.aggregated.vcf”, downloaded on 22-12-2022) and the “Human Somatic Short Variants” dataset from Ensembl (Release 110, July 2023) for the same transcripts. mRNA vaccine sequences were retrieved through a systematic search of patent databases. To verify reporter function, some test sequences included alternative AUG codons and in-frame stop codons. However, these sequences were excluded from downstream analysis because of interference with reporter readout.

#### Test sequences with alternative TISs

To investigate the impact of mRNA sequence on translation initiation beyond the TIS (Fig. 1g, Supplementary Fig. 1g), multiple sequence variants were generated for each gene. The endogenous TIS was replaced with a TIS of variable strength, with the same set of TISs applied across all genes. Sequences were constrained to exclude alternative AUG start codons and in-frame stop codons.

#### Test sequences for scanning mutagenesis

To locate the region within the mRNA that is most important for the control of start codon recognition, we performed scanning mutagenesis across entire test sequences. (Fig. 2a). The mRNA sequences were mutated in non-overlapping tiles of five nucleotides. Positions immediately adjacent to the start codon (e.g., −6 to –1, and +4, and +5) were excluded from analysis, as their alteration would introduce changes within the TIS itself. Consequently, the upstream tile spanned nucleotides −10 to −7, and the downstream tile spanned nucleotides +6 to +8. For the five nucleotide tiles, if the 5’UTR length was not a multiple of five, the most cap-proximal nucleotides were not mutated. Downstream of the start codon, nucleotides +6 to either +18, +33, +39 or +93 (Fig. 2a) were mutated. The tiles were always mutated in the same way; A was mutated to U and G to C, as well as the corresponding inverse substitutions.

#### Test sequences for establishment of the distal eTIS

To obtain the distal eTIS table (Fig. 2b) all possible single-nucleotide substitutions were introduced across the 34 nucleotides upstream and the first 34 nucleotides downstream of the TIS. These variants were included as individual elements in the library for 18 randomly selected human mRNAs with 5′UTRs longer than 70 nucleotides. Fixed TISs were incorporated for each gene to normalize for gene-specific TIS contributions.

#### Test sequences for the establishment of the TIS

To investigate the contribution of single nucleotides in the TIS to start codon recognition (Fig. 2c), each nucleotide of the endogenous TIS was systematically mutated to the three alternative nucleotides in 194 randomly selected transcripts with 5′UTR lengths between 26 and 35 nucleotides. The test sequences contained 18 nucleotides of coding sequence to preserve immediate downstream sequence features.

#### Test sequences without near-cognate start codons

To assess the contribution of near-cognate start codons to start codon recognition, all CUG and GUG start codons in test sequences derived from endogenous human mRNAs were mutated to CAG (Supplementary Fig. 1k). Test sequences with and without near-cognate start codons were included in the same sublibrary and quantified in the same experiment, enabling direct comparison under identical conditions.

#### Test sequences with homopolymeric nucleotide stretches

To assess the effect of homopolymeric nucleotides on start codon recognition (Supplementary Fig. 4f, h and j), mRNA transcripts lacking the nucleotides that composite these stretches were selected. Homopolymeric nucleotide stretches were then introduced into these transcripts to evaluate their contribution to start codon recognition. Fixed TISs were incorporated for each gene to normalize for gene-specific TIS contributions.

#### (Distal) eTIS optimization

To optimize the distal eTIS upstream of the start codon, all non-optimal distal eTIS residues were substituted for optimal nucleotides (Fig. 4a, c, f, and Supplementary Fig. 9b, c). In the CDS, non-optimal distal eTIS residues were only replaced if the original amino acid sequence could be preserved. When a nucleotide substitution would create an alternative start codon, the second-most favorable nucleotide was incorporated at that position. To evaluate how distal eTIS optimization influences start-codon recognition at near-cognate start codons (Fig. 4b, Supplementary Fig. 9a), the original AUG start codons were replaced with CUG and GUG start codons. When optimizing the entire eTIS region, the endogenous TIS was replaced with a strong TIS. The same TIS was applied consistently across all mRNA sequences.

#### Library design

Multiple sub-libraries were created based on 5’UTR length and/or experiment type, each equipped with a unique sequence for library specific PCR amplification. The final library elements were designed with the following structure: 5’- PCR adapter - PaqCI site (CACCTGC) – spacer – AACC – test sequence – GGCG – spacer – PaqCI site (GCAGGTG) – PCR adapter-3’ (one element per oligo). Or: 5’- PCR adapter - PaqCI site (CACCTGC) – spacer – AACC – Element#1 – GGCG – spacer – PaqCI site (GCAGGTG) – Spacer – PaqCI site (CACCTGC – spacer – AACC- Element#2 – GGCG – Spacer – PaqCI site (GCAGGTG) – PCR adapter-3’ (Two elements per oligo). The AACC sequence is part of the CMV promoter and the GGCG is part of the GS linker and was included for ligation. The libraries were synthesized as an oligopool (Twist Bioscience).

### Library cloning

Libraries were PCR amplified from the synthesized oligopool with Q5 DNA polymerase (NEB) according to the instructions provided by the manufacturer. The PCR products were purified using gel electrophoresis (GeneJET gel extraction kit, ThermoFisher) and digested with PaqCI (AarI, NEB) and finally purified using ethanol precipitation. The RiboScan reporter plasmid was digested with PaqCI, the main frame reporter with BsmBI (NEB). Digested backbones were purified using gel electrophoresis (GeneJET gel extraction kit, ThermoFisher). Library sequences were ligated into the reporter plasmid with T4 DNA ligase (NEB) with a molar ratio of 1:2 backbone over insert, for 16 hours at 16 °C. Ligated plasmids were ethanol precipitated and per library a test transformation was performed with DH5α bacteria as well as a transformation with the digested backbone to check for incomplete digestion. Single clones were isolated and sequence verified to ensure library integrity. Large-scale transformations were performed with MegaX DH10B electrocompetent cells (ThermoFisher). After recovery, cells were plated on selective agar and incubated overnight at 37 °C. Colonies were scraped, pooled to maintain library complexity, and plasmid DNA was isolated using a Maxiprep kit (PureLink™ HiPure Plasmid Maxiprep Kit, ThermoFisher).

### Reporter construction for single-variant validation

For validation purposes, single test sequences were cloned into the RiboScan and Main Frame reporters (Fig. 2e, Supplementary Fig. 1c, f, Supplementary Fig. 4e). DNA oligo’s containing the test sequence were synthesized (IDT) with the overhangs required for ligation into the digested reporter backbone (Supplementary Table 1, tab 5). The oligos were hybridized and ligated into the digested RiboScan reporter using Instant Sticky End Master Mix (NEB) with a 1:1 molar ratio of insert and backbone.

Sequences used for single-molecule imaging experiments and for Cryo-EM structure determination were first cloned into the RiboScan reporter using the same cloning strategy, followed by validation of their expression in HEK293T cells (Fig. 6c, Supplementary Fig. 10a, b).

### eTIS optimization of type I IRESs

To assess whether the eTIS contributes to start codon recognition during translation driven by type I IRES elements, we adapted the main-frame reporter. In the reporter, GFP serves as a normalization control and is expressed via cap-dependent translation. mCherry expression was driven from the same mRNA downstream of a variable type I IRES containing either weak or strong distal eTIS sequences (Fig. 4g). Type I IRES elements from Coxsackievirus B3 (CVB3) and poliovirus were PCR amplified from synthetic gene fragments (Twist Bioscience) using primers encoding strong or weak distal eTIS variants (Supplementary Table 1, tab 6). The distal eTIS sequences were inserted downstream of the sixth stem–loop domain of each IRES to preserve IRES activity. Lentiviruses were produced and used to transduce cells, followed by puromycin selection to establish stable reporter populations.

### FACS sample preparation and FACS analysis

Cells were grown to a confluency of 70 %, washed twice with PBS, dissociated with trypsin and resuspended in PBS. Cells were pelleted by centrifugation at 1200 rpm and subsequently resuspended in PBS containing 20 % (v/v) fetal bovine serum (FBS; Sigma-Aldrich) for FACS sorting. All FACS sorts were carried out using a FACS Aria Fusion (BD Biosciences) using a 488 nm and 532 nm laser to measure GFP and mCherry levels, respectively. Cells transduced with the RiboScan and Main frame reporter libraries were sorted into different gates based on the GFP/mCherry ratio (see Supporting Fig. 1a,b). To set the gates for the RiboScan library, cell lines expressing single RiboScan test sequences were created (see Supplementary Fig. 1c). Gates were set with equal spacing, and the same number of cells were sorted per gate. The coverage of each library element was approximately 400-fold. Cells were collected in 2 mL Eppendorf tubes and pelleted directly after sorting by centrifugation at 1200 rpm for 5 min at 4 °C, the supernatant was removed, and the samples were stored at -20 °C until library preparation, typically within 48 hours. Two biological repeats were sorted per library.

GFP and mCherry fluorescence levels were measured for single RiboScan and type I IRES reporter constructs. The GFP-to-mCherry ratio was calculated for the RiboScan reporters (Fig. 2e) and the mCherry-to-GFP ratio for the IRES reporters (Fig. 4h).

### FACS-seq library preparation and sequencing

For each biological replicate of each library, the test sequences from each sorted FACS gate were PCR amplified, barcoded, and sequenced. To achieve this, genomic DNA from the cells was extracted using a genomic DNA extraction kit (GeneJET gDNA extraction kit, ThermoFisher). PCR amplification was performed with a high-fidelity DNA polymerase (Q5, NEB) in the presence of high GC-buffer for 20 to 25 cycles. The reverse primer (5’-GTGACTGGAGTTCCTTGGCACCCGAGAATTCCANNNNNNNNGTGCCCGTTGACGTCGC-3’, for RiboScan reporter, 5’-GTGACTGGAGTTCCTTGGCACCCGAGAATTCCANNNNNNNNTGTGTCCGTTGACGTCTC -‘3 for the main frame reporter) contains an eight nucleotide long barcode corresponding to the FACS gate ID (See Supplementary Table 1, tab 7). The forward primer is complementary to the 3’ end of the CMV promoter (5’-TTCAGAGTTCTACAGTCCGACGATCGCAGAGCTGGTTTAGTGAACC-3’). PCR products were purified from an agarose gel (GeneJET gel extraction kit, ThermoFisher) and DNA concentrations were measured using a Qubit high sensitivity dsDNA assay (ThermoFisher). Purified PCR products for each library were pooled at equimolar ratios. TruSeq small RNA primers (Illumina) were used for the preparation of the Illumina sequencing libraries in a final PCR reaction ran for eight cycles. Each individual repeat of each library in a sequencing run was equipped with a unique TruSeq index. Sequencing libraries were sequenced on an Illumina NextSeq2000 platform (High output, 2 x 100 bp or 2 x 150 bp) or an Illumina NextSeq500 platform (Mid output, 2 x 150 bp).

### 5’RACE for TSS mapping

5′ Rapid Amplification of cDNA Ends (5′RACE) was performed to determine transcription start sites (TSSs) of RiboScan reporters. Total RNA was extracted from cells expressing the RiboScan reporter library using the RNeasy Kit (Qiagen). First-strand cDNA synthesis was carried out with a primer complementary to the GFP sequence in the RiboScan reporter construct (5′-TCTTGTAGGTCCCGTCGTCT-3′), using SuperScript III reverse transcriptase (Invitrogen) for 45 minutes at 55 °C. RNA was subsequently removed by adding 1 µL of RNase Cocktail Enzyme Mix (ThermoFisher) per 20 µL of reaction volume. cDNA was purified using AMPure XP beads (Beckman Coulter) at a 0.8x bead-to-sample ratio. The purified cDNA was A-tailed using terminal deoxynucleotidyl transferase (TdT; NEB) in the presence of a 1000-fold molar excess of dATP over the 3′ ends, resulting in A-tails of approximately 10–20 nucleotides. After a second bead cleanup (0.8x ratio), second-strand synthesis was performed using primer 5′-TTCAGAGTTCTACAGTCCGACGATCNNNNNNCCTTTTTTTTTTTTTTTTTV-3′, containing a six nucleotide long unique molecular identifier (UMI). Following another bead cleanup, PCR amplification (20 cycles) was performed using forward primer 5′-TTCAGAGTTCTACAGTCCGACGATC-3′ and reverse primer 5′-GTGACTGGAGTTCCTTGGCACCCGAGAATTCCAATGTTCAGGTGCCCGTTGACGTCGC-3′, with complementarity to the GFP sequence in the RiboScan reporter. Illumina index 1 and index 2 adapters were added in a final PCR (8 cycles). Libraries were sequenced on an Illumina NextSeq500 platform (Mid Output, 2 x 150 bp).

### Raw data processing and sequence alignment

Sequencing reads were trimmed using CutAdapt (Martin, 2011) (version 3.5). For TSS determination with 5’RACE, read pairs were filtered on the presence of the sequence motif NNNNNNNNCCTTTTTT in read one and a sequence that is complementary to the RiboScan reporter in read two (5’-AGAAGAAGCGCAAGGTTTCAAAAGGAGAAGAACTTTTCACCGGAGTTGTCCCAATTCTTGTCGAACTCGACGGCGACGTCAACGGGCACAAATTTTCCGTC-3’). The UMI was extracted and appended to the read name and the poly-T sequence was trimmed from read one. The trimmed reads were aligned to a custom build reference genome containing the test sequences using Bowtie2 in the sensitive local alignment mode (Langmead and Salzberg, 2012). To enable mapping of reads initiating within the CMV promoter, the sequence 5′-tctatataagcagagctggtttagtgaacc-3′, corresponding to the 3’end of the CMV promoter, was appended upstream of each test sequence in the reference genome.

For the FACS-seq sequencing libraries, the adapters, matching the CMV promoter (5’-GCAGAGCTGGTTTAGTGAACC-3’), and matching the GS linker of the RiboScan reporter or the main frame reporter (5’-GGCGGCAGCGGCTCAGGAA-3’) were removed from read one to obtain the complete test sequence. Untrimmed reads were removed. The FACS gate barcodes were obtained by obtaining the first eight bases from read two.

### TSS mapping

The TSS of the CMV promoter was mapped using 5′ RACE. Reads were first uniquely assigned to individual test sequences based on alignment to the custom reference genome. The position of the first matched nucleotide for each read was determined from the POS column in the SAM file generated by Bowtie2, corresponding to the leftmost aligned position. Only test sequences with >100 uniquely assigned reads were included in downstream analyses. To generate the histogram shown in Supplementary Fig. 2a, for each test sequence the nucleotide position relative to the 5′ end with the highest number of assigned reads was identified and included in the plot. To confirm that mutations introduced near the 5′ end of the mRNA do not alter TSS selection, representative 5′ RACE profiles were generated for several sets of mRNA variants (Supplementary Fig. 3a).

### Calculating GFP expression for each test sequence

The processed reads were analyzed using a custom R script to assign average GFP expression to individual test sequences. Trimmed read pairs were first filtered and grouped according to FACS gate barcodes. Read pairs were compared with the expected test (Read 1) and FACS gate barcode (Read 2) sequences, and pairs without an exact match to both were excluded from subsequent analyses. Test sequences with less than 50 assigned reads were excluded from further analysis. Read counts were normalized by gate barcode such that all gates contributed equally to the total read population. Subsequently, normalization was applied per test sequence to equalize the total number of reads across gates for each test sequence. Because the expression distribution of individual test sequences across FACS gates was anticipated to be confined to a limited fluorescence range (Supplementary Fig. 1c), peak calling was performed using a custom algorithm. Briefly, the algorithm identified a region comprising three consecutive FACS gates with the highest cumulative read abundance. Read numbers in gates outside this peak region were set to zero. The data were re-normalized such that the summed normalized read fraction for each element equaled one. The GFP intensity for each library element was then calculated by multiplying the normalized read fraction in each FACS gate by the corresponding mean GFP fluorescence per gate, and summing the resulting values across all gates. GFP intensities shown in each graph represent the average across two independent biological replicates. For each figure panel based on FACS-seq experiments, GFP intensity values from both biological replicates are provided in Supplementary Table 2.

### Assessment of differential start codon recognition through scanning mutagenesis

Differential expression analysis was performed to identify the percentage of genes experiencing alternative start codon recognition as a result of the mutated tile in the scanning mutagenesis screen (Fig. 2a). Differential expression analysis was performed using Limma (Ritchie et al., 2015). GFP intensity values from the RiboScan FACS-seq experiment were log10-transformed prior to analysis. Fold changes in RiboScan reporter expression (i.e. GFP) were calculated on a log_2_-transformed scale, and *P* values were adjusted for multiple testing using the Benjamini–Hochberg procedure. Mutations with a log_2_(Fold change) corresponding to >1.25-fold linear change and an adjusted *P* value <0.05 were classified as significantly altering start codon recognition. For each mutation tile, the percentage of genes exhibiting a significant change in leaky scanning relative to the corresponding WT sequence was computed and plotted as a function of mRNA position relative to the start codon. No distinction was made between increases and decreases in reporter expression, as the direction of the effect depends on the specific mutation introduced within each tile.

To generate the graph shown in Fig. 2a, data from two libraries were combined. For the upstream region, we used a library comprising genes in which 5′ untranslated regions (5′UTRs) of 90 nucleotides in length were mutagenized. For the region downstream of the start codon, we used a separate library of sequences with 5′UTRs of 30 nucleotides and 90 nts downstream of the start codon. In this library, only the downstream sequence was mutated.

### Calculation of the effects of (e)TIS mutation on start codon recognition efficiency

To generate the (e)TIS heatmaps shown in Figure 2b and 2c (distal eTIS and TIS, respectively), which represent the effects of single-nucleotide substitutions on start codon recognition efficiency, only those genes were included for which the GFP expression could be quantified for all four nucleotides at a given nucleotide position. For each gene, the mean GFP expression across the four nucleotide variants at each position was calculated, and log₂(Fold change) values were then derived for each nucleotide relative to this positional mean. The table represents the mean log_2_(Fold change) for each nucleotide at each position across all genes analyzed. Substitutions that created a start codon or a downstream in-frame stop codon were excluded from the analysis at that nucleotide position, as these events confound interpretation of the reporter signal.

To generate the distal eTIS tables shown in Supplementary Fig. 4a, b, transcripts with long (a) or short (b) 5′UTRs were grouped according to the introduced TIS. For each group, the average GFP intensity associated with each TIS was calculated. At every nucleotide position, the mean GFP intensity was then computed separately for each nucleotide. Nucleotide-specific effects were quantified as log₂(Fold change) values by comparing the nucleotide-specific positional intensity to the mean intensity of each position. The log_2_(Fold Change) values are provided in Supplementary Table 3.

The effect of individual distal eTIS positions on start codon recognition was visualized using dot plots. For each gene, we identified the two nucleotides exhibiting the largest difference in start codon recognition efficiency (the most optimal and least optimal) and calculated the log₂(Fold change) in GFP expression between these nucleotides (Fig. 2f–h and Supplementary Fig. 4l–o). To improve visualization outliers were removed from the graphs (the 2% of genes showing the weakest and strongest changes in start codon recognition were excluded from the plots).

### Modeling (e)TIS strengths

#### Development of RiboScanner; a deep convoluted neural network for eTIS strength determination

##### Model input

The RiboScanner model receives RNA sequences from the experimental libraries and predicts their associated eTIS strengths. Each sequence is one-hot encoded as a matrix of shape (156, 4), corresponding to the RNA nucleotides (A = [1, 0, 0, 0]; C = [0, 1, 0, 0]; G = [0, 0, 1, 0]; U = [0, 0, 0, 1]; N = [0, 0, 0, 0]). The input RNA fragments range in length from 26 to 129 nucleotides, and all sequences were padded to a fixed length of 156 nucleotides using N bases. The number of N bases added to the 5′ and 3′ ends of each fragment was randomly assigned following a uniform distribution to minimize positional bias. To account for potential motif formation at fragment boundaries, we appended a flanking adaptor sequences derived from the reporter backbone: AGTGAACC at the 5′ end and GGCGGCAG at the 3′ end of the sequence. Sequences with additional AUGs and in-frame stop codons downstream of the main start codon were retained in the training set to allow the model to learn from these features.

##### Deep-learning model architecture

The eTIS model combines CNN and RNN layers inspired by Zheng *et al*. (Zheng et al., 2023). The convolutional encoder was implemented as a 1D convolutional block composed of four sequential convolutional layers with output channel sizes of 128, 256, 256, and 256, respectively. Each convolution used a kernel size of 3 and a stride pattern of [1, 1, 1, 3], with padding of 1 to preserve sequence length prior to down sampling. All convolutional layers were followed by batch normalization and a Mish non-linear activation function. This convolutional tower transformed the one-hot–encoded DNA input (4 input channels) into a dense 256-dimensional representation capturing local sequence features. The resulting feature maps were then transposed and passed to a two-layer gated recurrent unit (GRU) network with a hidden size of 80, enabling the model to capture long-range dependencies within the encoded sequence. The last hidden state of the GRU was fed into a fully connected layer with a single output neuron to produce the final prediction. The model was implemented in PyTorch (v2.1.1).

##### Model training and evaluation

We split our 70,777 sequences into 11 equally sized folds, ensuring that sequences originating from the same gene were assigned to the same fold. One-fold was reserved as a test set for the final model assessment, while the remaining ten folds were used in a cross-validation scheme. For an overview of the training and validation dataset see Supplementary Table 4. Specifically, nine folds were used for training and one-fold for validation to determine the optimal hyperparameters. The final model is an ensemble of 10 models, each trained on a unique combination of nine folds.

We employed a mean squared error (MSE) loss function. The deep learning models were trained for 25 epochs with a batch size of 32. To optimize training, we used the stochastic gradient descent optimizer with a momentum of 0.9, weight decay of 0.0001, and a learning rate of 0.0005. L1 and L2 regularization terms (both set to 0.05) were applied to the loss function. Model training was performed on an NVIDIA Quadro RTX 6000 GPU.

#### Conversion of RiboScanner predicted leaky scanning values to an eTIS strength

RiboScanner was developed to predict GFP expression levels from RiboScan reporters for a given test sequence. To derive an eTIS strength score—representing the efficiency with which the start codon in the test sequence is recognized by the translation machinery—from these predictions, the predicted leaky scanning values were rescaled relative to the maximum predicted leaky scanning value observed within the test set. The transformation was performed using the following formula:

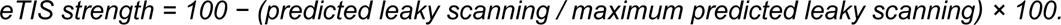

In our dataset, the maximum predicted leaky scanning value corresponded to a value of 16.010647.

#### eTIS strength calculation

Coding sequence AUGs and upstream AUGs were retrieved from the human transcriptome using BioMart (Ensembl release V115, September 2025). For each canonical mRNA transcript, a 73-nucleotide window (−40 to +33 nucleotides relative to the start codon) was extracted around the AUG for eTIS strength prediction using RiboScanner. Sequences containing additional AUG codons, in-frame stop codons, or 5′UTRs shorter than 12 nucleotides were excluded, as these features could confound eTIS strength estimation (Supplementary Fig. 2d, Supplementary Fig. 3b,c). Upstream AUGs were classified as translated or non-translated based on a previously published dataset (Mudge *et al*., 2022).

To estimate eTIS strengths for near-cognate start codons within 5′UTRs, nucleotide sequences encompassing at most 40 nucleotides upstream and 30 nucleotides downstream of all CUG and GUG codons in human 5’UTRs were obtained. Because RiboScanner was not trained on mRNA sequences containing the near-cognate start codons CUG and GUG, these codons were computationally substituted with AUG to enable eTIS strength prediction. As above, sequences containing additional AUGs, in-frame stop codons, or 5′UTRs shorter than 12 nucleotides were excluded to avoid interference with eTIS strength predictions. Near-cognate start codons were classified as translated or non-translated based on a previously published dataset (Mudge *et al*., 2022).

#### RiboScanner TIS strength calculation

To determine the relative strength of each TIS in promoting start codon recognition, we randomly selected 100 representative distal eTIS contexts from human mRNAs, spanning nucleotides −40 to −7 and +6 to +33 relative to the start codon. All selected distal eTISs contained a single AUG codon. Each of the 65,536 possible TIS variants was computationally introduced into all 100 distal eTIS contexts, and eTIS strengths were predicted using RiboScanner. For each TIS variant, the TIS strength was defined as the mean predicted eTIS strength across all distal eTIS contexts. The RiboScanner TIS strengths are provided in Supplementary Table 4, tab 5.

Noderer TIS strength scores (Noderer *et al*., 2014), RiboScanner TIS strength scores, and RiboScanner eTIS strength scores were compared with GFP expression levels from reporter mRNA constructs containing either endogenous mRNA sequences (Fig. 1e) or mRNA sequences from the test set (Fig. 2i–k and Supplementary Fig. 5f, g). Correlations between GFP expression and Noderer TIS strength scores were higher for all tested mRNA sequences (which included both endogenous sequences, random sequence and various mutated sequences) than for endogenous mRNA sequences only. The likely reason for this is that the larger synthetic sequence libraries contained sequences with very poor TIS strengths, which were predicted more accurately by the Noderer TIS score, yielding a slightly higher overall correlation coefficient.

### Ribosomal profiling analysis

Human translation initiation profiling datasets (see Supplementary Table 6) were obtained from GWIPS-viz (Michel et al., 2018) and concatenated into one dataset. The eTIS regions for all upstream CUG and GUG codons were defined as described in the section “*eTIS strength calculation”.* Reads mapping to each CUG or GUG start codon and the corresponding AUG start codon from the main start codon were summed, and only start codon pairs with ≥100 total assigned reads were retained for further analysis. Upstream start codons were classified as translated when reads mapping to the upstream codon represented ≥10% of those mapping to the canonical AUG codon. Representative examples of translated near-cognate start codons are shown in Supplementary Fig. 6e, f.

### mRNA secondary structure prediction

The RNAfold function of the ViennaRNA package (Version 2.5, 11-8-2021) (Lorenz et al., 2011) was used to calculate the minimal free energy. Sequences spanning 15 nucleotides upstream and downstream of the start codon were used for mRNA secondary structure determination.

### Transcriptome-wide analysis of nucleotide enrichment in the eTIS

To analyze enrichment of specific nucleotides at each position within the eTIS, all mRNA sequences for each investigated organism were obtained from Ensembl using BiomaRt (Smedley *et al*., 2009) (datasets listed in Supplementary Table 6). For Saccharomyces cerevisiae, 5′UTR sequences were derived from mRNA sequencing data reported by Pelechano *et al*. (Pelechano et al., 2013). Transcripts were filtered on canonical status, only Ensembl canonical transcripts were considered, to ensure equal weighing of all genes.

We initiated the analysis by comparing non-normalized nucleotide frequencies within the eTIS region between transcripts containing or lacking an intron in the 5′ untranslated region (5′UTR) (Fig. 3b; Supplementary Fig. 7a). A similar comparison was performed for the TIS region. For the latter analyses, transcripts with a 5′UTR intron were additionally required to have the second exon begin at least 1.000 nucleotides downstream of the transcription start site (TSS), thereby minimizing biases arising from promoter-associated GC enrichment. Nucleotide frequencies at TIS positions for either genes with or without introns (Fig. 3c) were subsequently compared with log₂(Fold changes) in start codon recognition (Fig. 2c).

To assess enrichment of specific nucleotides in the distal eTIS across the transcriptome, analyses were performed either on all mRNAs collectively (Fig. 3d) or restricted to transcripts with (Supplementary Fig. 7e) or without (Supplementary Fig. 8a) a 5’UTR intron. Only transcripts with 5′UTRs of at least 40 nucleotides were retained to ensure complete representation of the 5′UTR in the heatmap representing nucleotide enrichment of the eTIS. For each transcript, the 34 nucleotides upstream of the TIS were extracted. For transcripts containing a 5′UTR intron, an additional filter was applied to ensure that all 34 nucleotides were located within the second exon. Nucleotide frequencies were calculated across all 34 nucleotide positions for each of the transcripts groups. Percentage enrichment at each position for each nucleotide was determined by dividing the nucleotide frequency at that position by the average nucleotide frequency in the genome over all 34 upstream distal eTIS nucleotides.

Downstream of the TIS, nucleotides +6 to +33 were analyzed for each group of transcripts. For transcripts lacking a 5′UTR intron, an additional filter was applied to ensure that all 33 nucleotides were contained within the first exon. Because nucleotide frequencies vary across the three codon positions owing to codon usage and amino acid composition, average frequencies were calculated separately for each codon position rather than across the region as a whole. Percentage enrichment at each position was then obtained by dividing the nucleotide frequency at that position by the average nucleotide frequency of the corresponding codon position measured over the 10 codons under analysis.

A similar approach was applied to calculate nucleotide enrichment around upstream UAC codons (Fig 3d, f) and upstream near-cognate codons (CUG/GUG), which had previously been classified as translated (Fig. 3d, g) or non-translated (Supplementary Fig. 7h) (Mudge *et al*., 2022). For the latter analysis, sequences containing additional start codons or downstream in-frame stop codons were excluded, as these features may impair initiation at the candidate start codon and thereby weaken selection on eTIS bases. For a list of included sequences see Supplementary Table 5.

To compare the nucleotide enrichment of eTIS residues among different species (Supplementary Fig. 8a-d), we limited the analysis to all mRNA sequences without a 5’UTR intron and with a minimal 5’UTR length of 120 nucleotides. Nucleotide enrichment was calculated as described above, but here for two regions within the 5′UTR: a control region, spanning nucleotides -100 to -61 and the eTIS region (-40 to -7). The 120-nucleotide cutoff for 5′UTR length was applied to mitigate potential nucleotide biases associated with sequences proximal to the 5′ cap of the mRNA. The obtained enrichment values for each region were compared between *H. sapiens* and nine other model organisms (Supplementary Fig. 8d).

### Nucleotide enrichment per type of amino acid

Nucleotide enrichment analysis was performed on a per-codon basis (Fig. 3i, Supplementary Fig. 7k and Supplementary Fig. 8e), to assess whether nucleotides favored for efficient start codon recognition are enriched independently of the encoded amino acid. Codons were first grouped by the nucleotide composition of the first and second nucleotide and by the identity of the encoded amino acid. Then a global nucleotide frequency was calculated for each group across the first 300 codons of all coding sequences. Nucleotide enrichment at each position was then determined by subtracting the global nucleotide frequency from the position-specific frequency. Finally, position-specific nucleotide enrichments, for example, at the +6 nucleotide, were calculated by subtracting the mean nucleotide enrichment of the first ten codons, excluding the position under analysis, from the nucleotide frequency of the position being accessed. The same procedure was performed for other positions (e.g., +9, +12, +15, up to +30). Nucleotide enrichments were compared to codon optimality values as determined by the codon adaptation index (CAI) (Sharp and Li, 1987) and with the measured effect of each nucleotide on start codon recognition (taken from Fig. 2b). A similar analysis was performed for additional organisms (Supplementary Fig. 8e). For *S. cerevisiae*, all transcripts were included.

### Real-time single-molecule imaging assay

To produce the short model mRNA, DNA encoding the reverse complement of the mRNA were purchased from IDT as single-stranded oligos (See Supplementary Table 1, tab 8, “Sequences for single-molecule imaging”, for a list of sequences). The RNAs were transcribed *in vitro* and modified to contain a 5’-m7G cap and 3’-biotin as previously described (Grosely *et al*., 2025). All real-time single-molecule experiments were performed on a PacBio RS II and experimental movies analyzed using MATLAB R2024b as described previously (Grosely *et al*., 2025). The rates and rate constants are summarized in Supplementary Table 7.

### Cryo-electron Microscopy

#### Cryo-EM sample preparation

Native human 40S ribosomal subunits and initiation factors eIF2, eIF3, eIF4G and eIF4E were purified from HeLa cell lysate (Pisarev et al., 2007; Yi *et al*., 2022), and recombinant factors eIF1, eIF1A, eIF5, eIF4A, eIF4B and [^3^H]Met-tRNA ^Met^ were produced as described (Yi *et al*., 2022). The two mRNAs for cryo-EM analysis were derived from a model mRNA with an unstructured 5’ untranslated-region (Kumar et al., 2016; Yi *et al*., 2022), and each contained an identical suboptimal TIS (*CGCACC**AUG**GA*), but distinct distal eTIS sequences (Supplementary Table 1, tab 9, “Sequences for Cryo-EM structures”). Distal eTIS sequences were designed to yield optimal or non-optimal start codon recognition efficiencies. mRNAs were transcribed *in vitro* (Yi *et al*., 2022), capped using the Vaccinia Capping System (New England Biolabs), and purified with the Monarch® Spin RNA Cleanup Kit (New England Biolabs).

To assemble 48S initiation complexes, 40S subunits (0.3 μM) were incubated for 5 min at 37 °C with eIF1 (2.7 μM), eIF1A (2.7 μM), eIF2 (0.6 μM), eIF3 (0.7 μM), eIF4E (0.2 μM), eIF4G (0.3 μM), eIF4A (1.2 μM), eIF4B (0.5 μM), eIF5 (2.7 μM), [^3^H]Met-tRNA ^Met^ (1.6 μM) and mRNA (1.2–1.3 μM) in reaction buffer (20 mM HEPES pH 7.5, 95 mM potassium acetate, 3.75 mM magnesium acetate, 1 mM ATP, 0.5 mM GDPCP, 2 mM DTT, 0.25 mM spermidine). Complexes were subsequently stabilized by crosslinking with 3.5 mM bis(sulfosuccinimidyl)suberate (BS3; Sigma Aldrich) for 10 min at room temperature.

#### Cryo-EM analysis

For grid preparation, 4.5 μL of 48S reaction mixture were applied to glow-discharged Quantifoil 3.5/1 grids (Jena Bioscience) coated with prefloated continuous carbon. Grids were manually blotted with pre-wetted Whatman #1 filter paper and plunge-frozen in liquid ethane at 4 °C and 95% humidity using a custom-built device.

Cryo-EM data were collected on a Titan Krios G2 transmission electron microscope (Thermo Fisher Scientific) operated at 300 kV in EFTEM mode and equipped with a Quantum LS energy filter (Gatan) using a 20-eV slit width. Automated data acquisition was performed using SerialEM (Mastronarde, 2005) on a K3 detector (Gatan) in counting mode at a nominal magnification of 81,000× (1.05 Å/pixel), with a defocus range of 0.5–2 μm and a total exposure of 40 e/Å² fractionated over 40 movie frames.

#### Image processing

Image processing for each dataset was carried out independently using the same workflow in Relion 5.0 (Scheres, 2012) (Supplementary Fig. 11a), unless stated otherwise. Movies were motion-corrected, dose-weighted, and averaged, and contrast-transfer function (CTF) parameters were estimated. Particle picking was performed in Warp (Tegunov and Cramer, 2019). After fourfold-binned extraction, particles were subjected to 2D classification and 3D sorting to assess particle quality and eIF3 occupancy. Particles containing eIF3 were then re-extracted at full resolution (1.05 Å/pixel), refined to high-resolution, and further sorted for global quality, and eIF3 occupancy using alignment-free focused classification with signal subtraction (FCwSS) (Scheres, 2016). High-quality particles were subsequently subjected to CTF refinement (Zivanov et al., 2020) and Bayesian polishing (Zivanov et al., 2019), followed by further refinement. Particles were then classified by FCwSS focused on the 40S head, yielding two major 48S populations corresponding to open and closed conformations (Fig. 6d). Each population was further classified by FCwSS to assess ternary complex occupancy; open 48S particles were additionally sorted according to mRNA configuration, whereas closed 48S particles were sorted based on the presence or absence of eIF5 (Fig. 6e). In open 48S structures assembled on weak or strong eTIS mRNAs, the mRNA density was poorly resolved throughout, including at the decoding center, despite high overall resolution. Extensive classification using FCwSS did not resolve codon-anticodon interactions, or a defined mRNA path, indicating substantial flexibility and heterogeneity of mRNA positioning in the open state and precluding detailed structural analysis. In contrast, in closed 48S structures the start codon and surrounding mRNA in the decoding center were well resolved. Notably, closed 48S particles with or without eIF5 displayed no discernible differences in mRNA configuration; therefore, subsequent analyses were performed on merged closed 48S datasets combining eIF5-bound and -unbound particles.

To resolve distal eTIS interactions in the closed 48S at the dynamic periphery, particularly at the mRNA entry and exit channels, targeted classifications were performed (Supplementary Fig. 11a). At the mRNA entry site, particles were sorted by FCwSS focusing on distal mRNA elements including eIF3g. For the mRNA exit-eIF3d region, FCwSS was first used to classify particles according to the position of the eIF3d central domain and then according to mRNA-exit conformation. For the closed 48S particles assembled on the strong eTIS mRNA, these additional sorting steps enabled detailed interpretation of distal eTIS contacts at the mRNA entry, mRNA exit and eIF3 core region. In contrast, analogous sorting of particles with weak eTIS did not improve density sufficiently for comparable analysis. Final particle subsets, including those obtained by local classification, were refined to high resolution using the full particle box and subsequently post-processed by sharpening or low-pass filtering, as appropriate (Supplementary Fig. 11b, c and Supplementary Tables 8, 9). Overall maps were visualized using LocScale 2.0 rendering (Bharadwaj et al., 2025). Figures were prepared in ChimeraX 1.10 (Meng et al., 2023).

#### Atomic model building and refinement

Atomic model building was performed for the closed 48S complexes assembled on mRNAs with weak and strong eTIS sequences. Initial models were based on the previously reported closed “48S-2” structure (PDB 8PJ2; Petrychenko et al. (Petrychenko *et al*., 2024)), with modifications and ion placements guided by the 1.9 Å human 80S ribosome structure (PDB 8QOI; Holvec et al. (Holvec et al., 2024)). eIF3d residues 2–20 and eIF3e positions 110-121 were modelled based on previous work (PDB: 7A09 Kratzat et al. (Kratzat et al., 2021)). To guide modelling of eIF3d, a full-length structure prediction was generated using the AlphaFold3 web server (Abramson et al., 2024) (random seed 131217477) with the canonical human eIF3d sequence (UniProt ID O15371; (UniProt, 2023)). Initial models were rigid-body fitted into the maps using ChimeraX 1.10 (Meng *et al*., 2023), followed by flexible fitting and manual adjustment in WinCoot 0.9.8.96 and 1.1.17 (Emsley et al., 2010).

Manual model building was primarily performed using sharpened maps at the final resolution, with the following exceptions. For the closed 48S with weak eTIS mRNA, residues -27 to -4 and +7 to +11 were built into maps low-pass filtered to 6 Å. For the closed 48S with strong eTIS mRNA, residues −6 to −4 were built using maps filtered to 6 Å, whereas residues +8 to +13 at the entry site were built at 4.5 Å. Locally classified maps of the closed 48S with strong eTIS mRNA were aligned to the overall map based on the respective region of interest. For the peripheral mRNA entry region, the locally classified “entry” map filtered to 4 Å was used to model interactions and residues −14 and −15, and to 4.5 Å for residues +16 to +21 and eIF3g. The mRNA exit region and eIF3 core, including full-length eIF3d, were modelled using the “exit–eIF3d” map obtained by local classification. In this region, mRNA residues -14 to -7 and the previously unresolved eIF3d α-helix (residues 125–145) were interpreted at 4.5 Å, whereas 18S rRNA, eS26 and eIF3a were modelled at 3.5 Å. The remaining eIF3 core was modelled at 3.5–7 Å resolution, including eTIS residues -27 to -15 and additional regions of eIF3d (residues 77–124 and 146–161).

Final models were refined iteratively using phenix.real_space_refine in Phenix 1.21.2-5419 (Liebschner et al., 2019) (Supplementary Table 8). Refinement consisted of 3–5 macrocycles with 200–500 iterations each, including ligand restraints generated using the Grade2 web server (Global Phasing), metal coordination restraints generated with phenix.ready_set, and secondary structure restraints. Manual corrections were performed in WinCoot between refinement runs to improve geometry and map fit.

## Data availability

A selection of raw images that support the findings of this study are openly available (https://doi.org/10.6084/m9.figshare.31730332). All RiboScan data generated in this study have been deposited at the Gene Expression Omnibus (GEO) under accession number (GSE324577). Cryo-EM maps and associated coordinates of atomic models were deposited to the EMDB and PDB with the following accession codes: (XXXX). Cryo-EM micrographs and particle images were deposited to the Electron Microscopy Public Image Archive (EMPIAR) with accession code (XXXX). Plasmids generated for this study can be requested from Addgene (Marvin Tanenbaum Lab).

## Code availability

Analysis scripts and the script for the RiboScanner model are available on Github (https://github.com/TanenbaumLab/RiboScan, and https://github.com/luciabarb/RiboScanner).

## Acknowledgements

We thank members of the Tanenbaum lab for helpful discussions. We thank M. Rodnina for providing access to the infrastructure of her department, as well as generous support, C. Dienemann and U. Steuerwald for maintenance of the cryo-EM facility at MPI-NAT, O. Geintzer, V. Herold, F. Hummel, S.Kappler, C. Kothe, A. Pfeifer, and M. Zimmermann for expert technical assistance, and S. Urlenka, M. Klein, M. Fechner and A. Esztermann for support in high-performance computing. In addition, we would like to thank the Hubrecht Sorting Facility as well as the Utrecht Sequencing Facility, subsidized by the University Medical Center Utrecht and the Netherlands X-omics Initiative (NWO project no. 184.034.019). This research was supported by the Max Planck Society (N.F.). B.M.P.V., L.B., M.D.M., J.d.R. and M.E.T. were supported by Oncode Institute, which is partly funded by the Dutch Cancer Society (KWF). M.E.T. and B.M.P.V. were supported by a grant from the Dutch Research Council (NWO/016.VIDI.189.005).

## Author contributions

B.M.P.V and M.E.T. conceptualized the study. B.M.P.V., D.L. and C.A. performed the experimental investigation. B.M.P.V. acquired the data for the *in vivo* measurements of start codon recognition and sequencing experiments. D.L. acquired the Cryo-EM data. C.A. acquired the single-molecule imaging data, supervised by J.D.P.. L.B-M. trained the convolutional neural network models and wrote the RiboScanner GitHub repository, supervised by J.d.R. M.S. performed analysis of ribosome profiling data, supervised by E.V.. B.M.P.V., D.L., L.B.-M., C.A., V.P., M.S. and N.F. conducted formal analysis. B.M.P.V., D.L., V.P., M.D.M. and N.F. performed validation. B.M.P.V, D.L., N.F. and M.E.T. wrote the original draft of the manuscript. All authors reviewed and edited the manuscript. N.F. and M.E.T. supervised the study and acquired funding.

## Supplementary Figure Legends

**Supplementary Figure 1.**
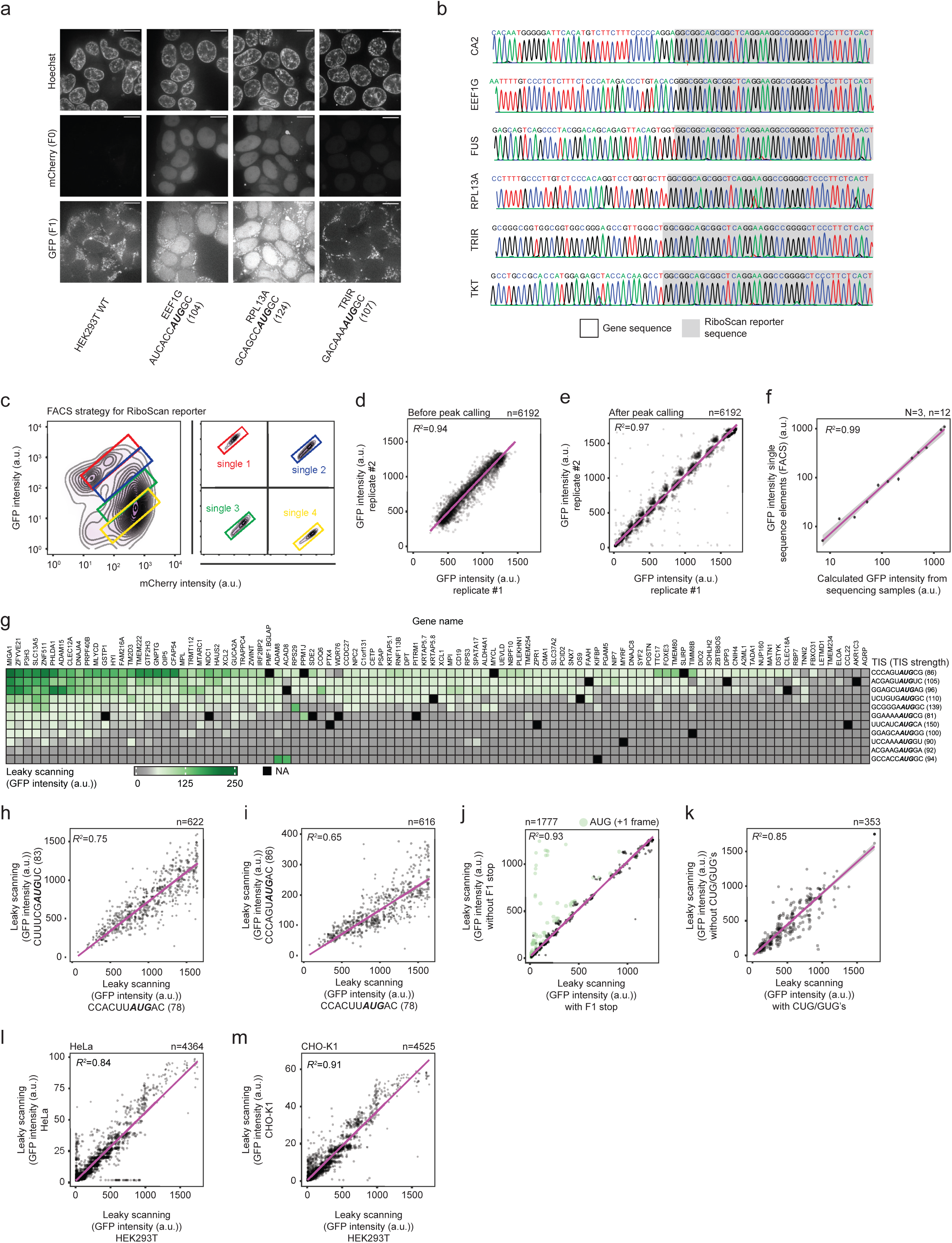
Gene sequences beyond the TIS control start codon recognition. **a,** Example microscopy images of endogenous genes in which the RiboScan reporter was inserted. The gene name, TIS and TIS strength (between brackets) are indicated. Scale bars, 10 µm. **b**, Genotyping results. PCR fragments were subjected to Sanger sequencing to confirm correct integration of the RiboScan reporter into the respective genes. **c**, GFP and mCherry expression levels for four representative single RiboScan reporters were determined by FACS (right set of panels). FACS plots are shown as examples. The gating angle used to sort cells from the libraries (left panel) was defined based on multiple single reporters. **d**, **e**, Analysis of experimental reproducibility. Either GFP intensities (**d**) or GFP intensities after peak calling (**e**) (See Methods) were compared between two biological replicates. **f**, Calculated GFP intensities for individual RiboScan reporters as determined by the sequencing-based approach were compared with GFP intensities of single RiboScan reporters tested by FACS individually. **g**, Heatmap representing GFP expression of the RiboScan reporter of indicated mRNA sequences (columns) combined with indicated TISs (rows). The strength of each TIS is indicated between brackets. Black boxes represent missing data points. Same as Fig. 1g for strong TISs. **h**, **i**, Comparison of GFP intensities from RiboScan reporters containing different TISs. **j**, Comparison of GFP intensities for RiboScan reporters with or without a stop codon in frame 1 downstream of the main start codon. Note that a few mRNAs show strongly reduced GFP expression upon introduction of the stop codon. These mRNAs contained an AUG codon in the GFP frame (F1). **k**, Comparison of GFP intensities for RiboScan reporters with or without CUG and GUG codons in frame 1. **l**, **m**, Comparison of GFP expression of the RiboScan reporters in Hela cells (**l**) or CHO-K1 cells (**m**) with the RiboScan GFP values in HEK293 cells. The observed differences in GFP expression likely reflect cell-type-specific differences in reporter construct expression. Each dot in **d**-**f**, **h**-**m** represents a single mRNA sequence. Linear regression fits are shown in purple, with grey shaded regions indicating the 95% confidence intervals.

**Supplementary Figure 2.**
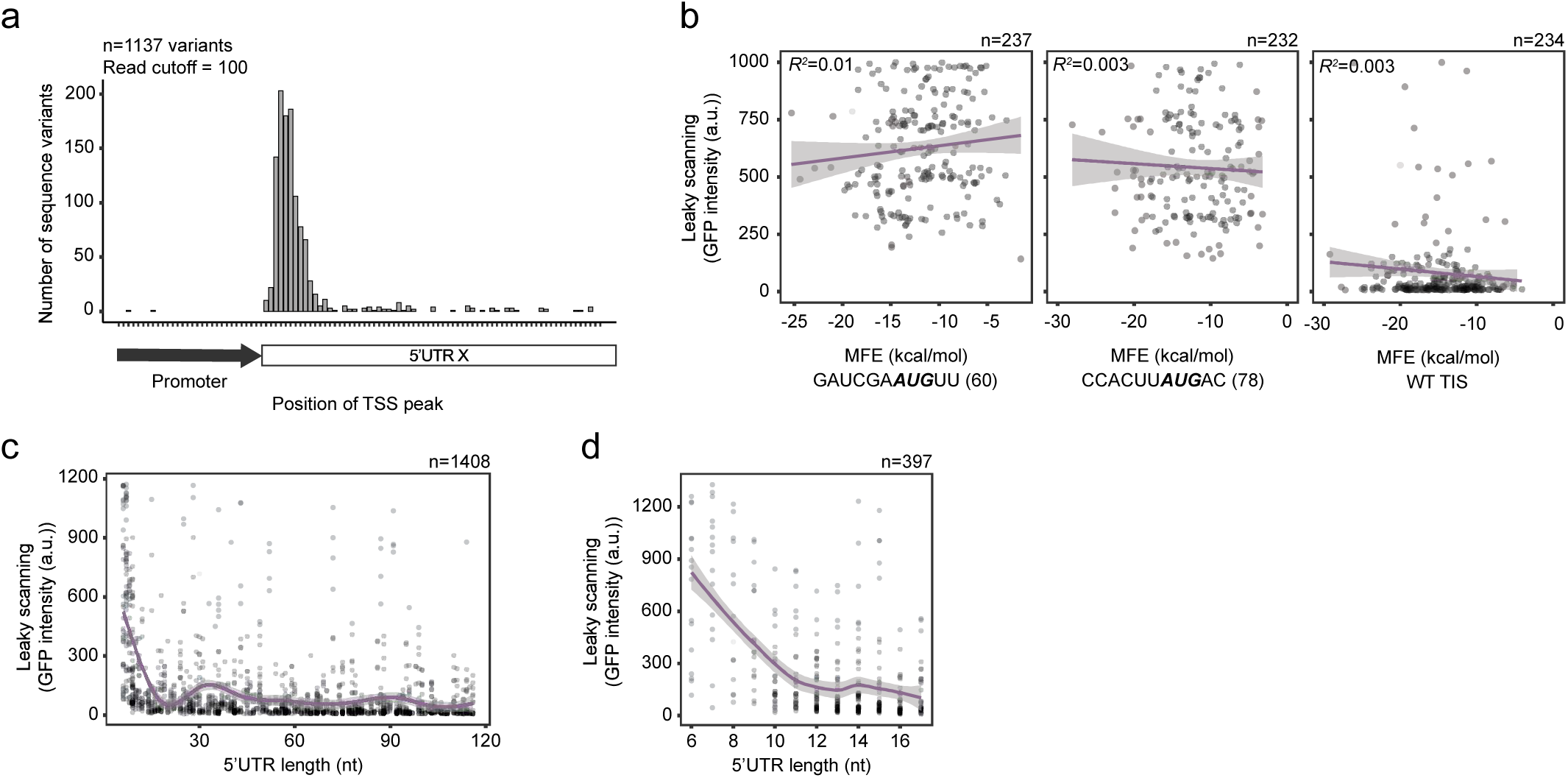
Impact of diverse mRNA features on start codon recognition. **a**, The TSS was mapped using 5’RACE and the most common TSS position across all RiboScan reporters is plotted. Note that the TSS is similar among different mRNAs. **b**, The mean free energy (MFE), a measure of RNA secondary structure, of the variable sequences (i.e. sequences spanning 15 nucleotides upstream and downstream of the start codon) was determined for each RiboScan reporter. MFE values are plotted against GFP expression for each mRNA. The TIS and TIS strength (between brackets) are indicated below each scatterplot. **c**, **d**, GFP expression of the RiboScan reporter plotted as a function of 5’UTR length. No correlation was observed between 5’UTR length and GFP expression, except at 5’UTR lengths of <12 nts, consistent with previous studies showing that start codon recognition efficiency is low if the start codon is very close to the 5’ end of the mRNA. Purple lines in **b** represent linear regression fits, with grey shaded regions indicating the 95% confidence intervals. Purple lines in **c**, **d** represent the smoothed trend lines, with grey shaded regions indicating the 95% confidence intervals.

**Supplementary Figure 3.**
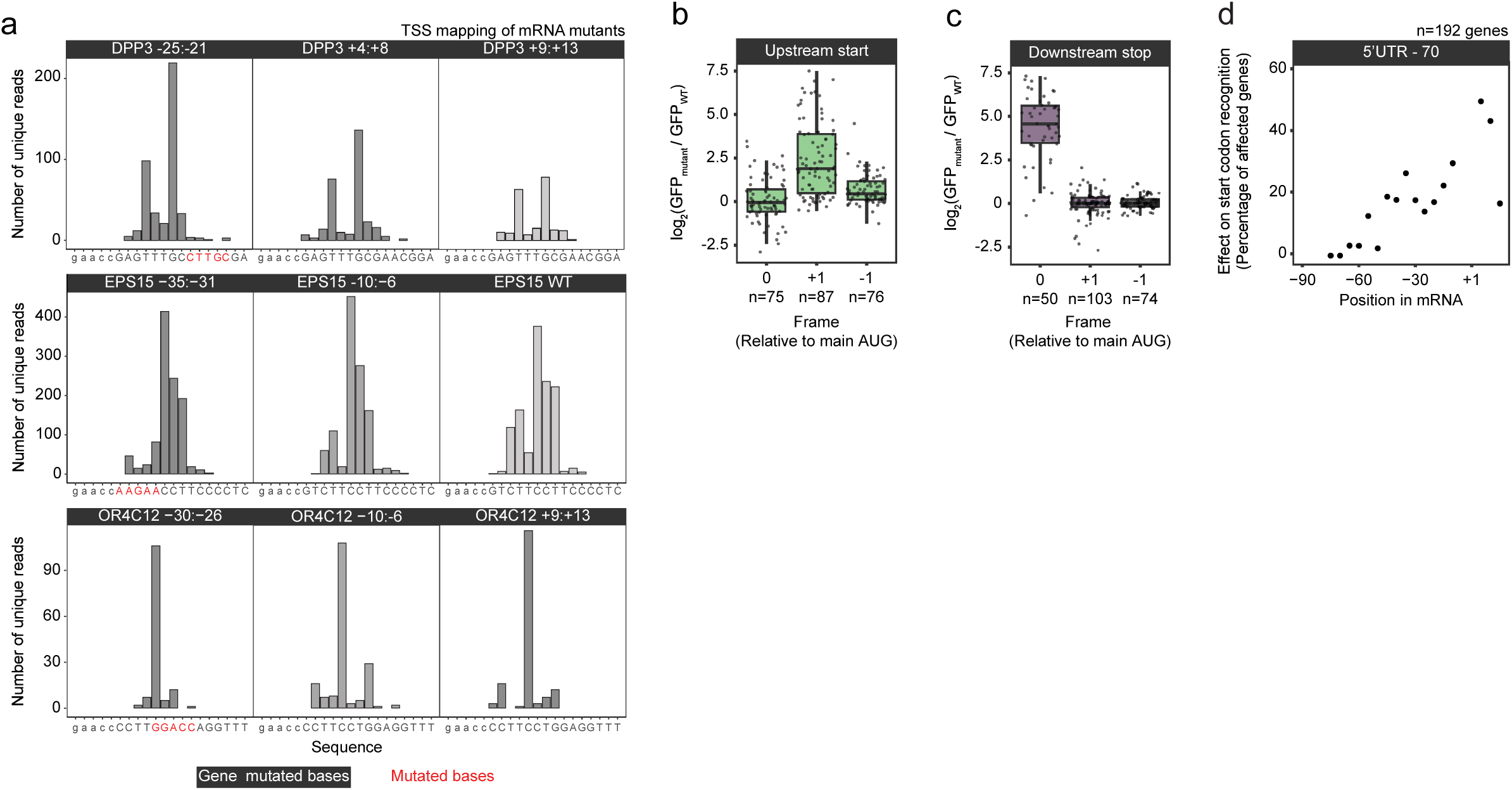
Scanning mutagenesis identifies sequence regions involved in start codon recognition. **a**, Examples of TSS mapping for different mRNAs of the scanning mutagenesis approach. Mutated bases are highlighted in red. Lowercase letters denote sequences derived from the CMV promoter, whereas uppercase letters indicate the RiboScan test sequence. TSSs are not affected by 5’UTR mutations. **b**, Impact of the introduction of an AUG in the GFP frame (F1) on GFP expression of the RiboScan reporter. **c**, Impact of the introduction of a stop codon in frame 0 downstream of the main start codon on GFP expression of the RiboScan reporter. The observed increase in GFP expression is likely due to termination at the stop codon of ribosomes that initiated at the main start codon, followed by re-initiation at the downstream AUG in-frame with GFP. **d**, Scanning mutagenesis was performed on mRNAs with a 5’UTR length of 70 nts in which blocks of 5 consecutive nucleotides were mutated. The fraction of mRNAs in which a mutation caused a change in GFP expression of >1.25 fold is plotted for each position relative to the start codon.

**Supplementary Figure 4.**
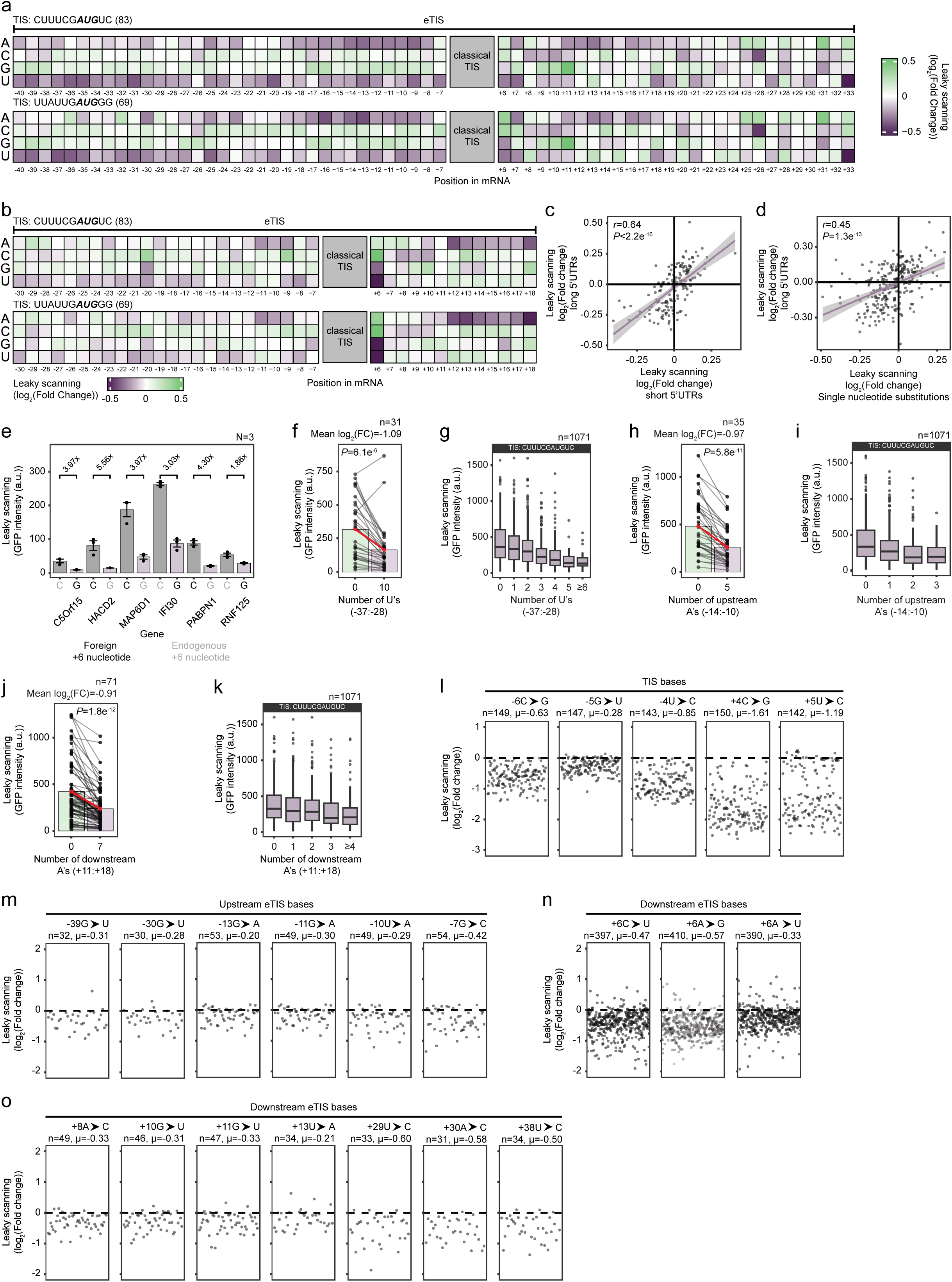
Start codon recognition is modulated by the eTIS. **a**, Heatmap representing the average log₂(Fold Change) in GFP expression of the RiboScan reporter for all mRNAs containing the indicated nucleotide at the indicated position relative to the start codon, relative to the average GFP expression across all genes (See Methods). Two distinct TISs were inserted in each gene. Note that the library used for this analysis only included mRNA coding sequences up to the +33 position. **b**, Same as **a**, but for mRNAs with short 5′ UTRs (<40 nucleotides). Note that the library used for this analysis only included mRNA coding sequences up to the +18 position. **c**, Comparison of the average log₂(Fold Change) in leaky scanning of each nucleotide of each position of the eTIS between mRNAs with long 5′UTRs (**a**) and short 5′UTRs (**b**). **d**, Comparison of the log₂(Fold Change) in leaky scanning per nucleotide per position of the eTIS obtained from single-nucleotide substitutions (Fig. 2b) and transcripts with long 5′ UTRs (**a**). Linear regression fits are shown in purple, with grey shaded regions indicating the 95% confidence intervals. Pearson correlation coefficients (*r*) and corresponding *P* values are indicated for each panel to assess the strength and significance of the linear relationships. **e**, Example genes in which the +6 nucleotide was mutated to either an optimal G or a non-optimal C. GFP intensities of the RiboScan reporter are shown. The endogenous +6 nucleotide is indicated in gray. **f–k**, Effects of indicated homopolymeric nucleotide stretches on start codon recognition efficiency. GFP intensities were compared for multiple individual mRNA sequences with or without the homopolymeric stretches (**f**, **h**, **j**), or for sequential reductions in the number of nucleotides in the homopolymeric stretch (**g**, **i**, **k**). A fixed TIS (CUUUCG***AUG***UC) was used in all mRNAs (**g**, **i**, **k**). Bars and red lines in (**f**, **h**, **j**) indicate mean values. *P* values were determined using a paired Wilcoxon signed-rank test. Box plots in (**g**, **i**, **k**) display the median (center line) and interquartile range (25th–75th percentiles; box), with whiskers extending to 1.5x the interquartile range. **l-o**, log_2_(Fold Change) in GFP expression of the RiboScan reporter when comparing the optimal nucleotide with the least optimal nucleotide for indicated positions within the TIS (**l**) and distal eTIS (**m**-**o**). Each dot corresponds to an mRNA sequence carrying the specified substitution at the indicated position.

**Supplementary Figure 5.**
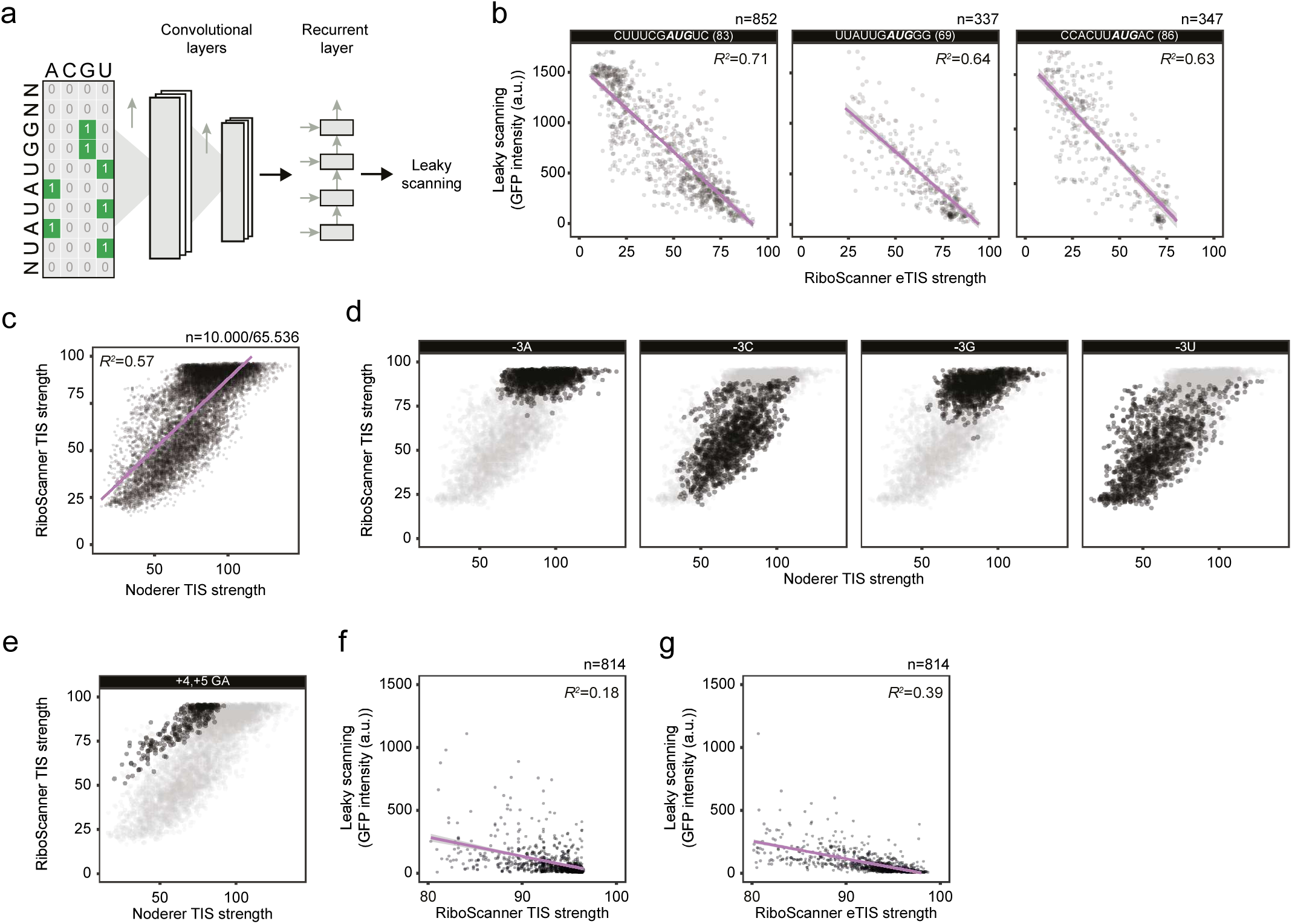
RiboScanner: a deep convolutional neural network for eTIS strength predictions. **a**, Overview of the RiboScanner model. The model consists of a convolutional neural network followed by a multi-layer recurrent neural network. It takes a one-hot–encoded RNA sequence as input and predicts its GFP (leaky-scanning) value. Early layers learn simple, motif-like features, whereas deeper layers capture more complex combinations of sequence elements. **b**, Comparison between predicted eTIS strength and measured GFP intensities using the RiboScan reporter assay. Reporters with three distinct TISs were examined. **c**, **d**, Comparison of the two indicated TIS strength metrics. Data was plotted separately for the different nucleotides at the -3 position (**d**). **e**, The TIS strength obtained with the eTIS model shows a distinct prediction for sequences containing a +4,+5 GA sequence compared to the Noderer TIS strength (Noderer *et al*., 2014). **f**, **g**, Zoom-in of bottom right part of the graph of Fig. 2i, j. TIS strength (this study) and eTIS strength are plotted as a function of GFP expression of the RiboScan reporter. All Linear regression fits (**b**, **c**, **f**, **g**) are shown in purple, with grey shaded regions indicating the 95% confidence intervals around each regression line.

**Supplementary Figure 6.**
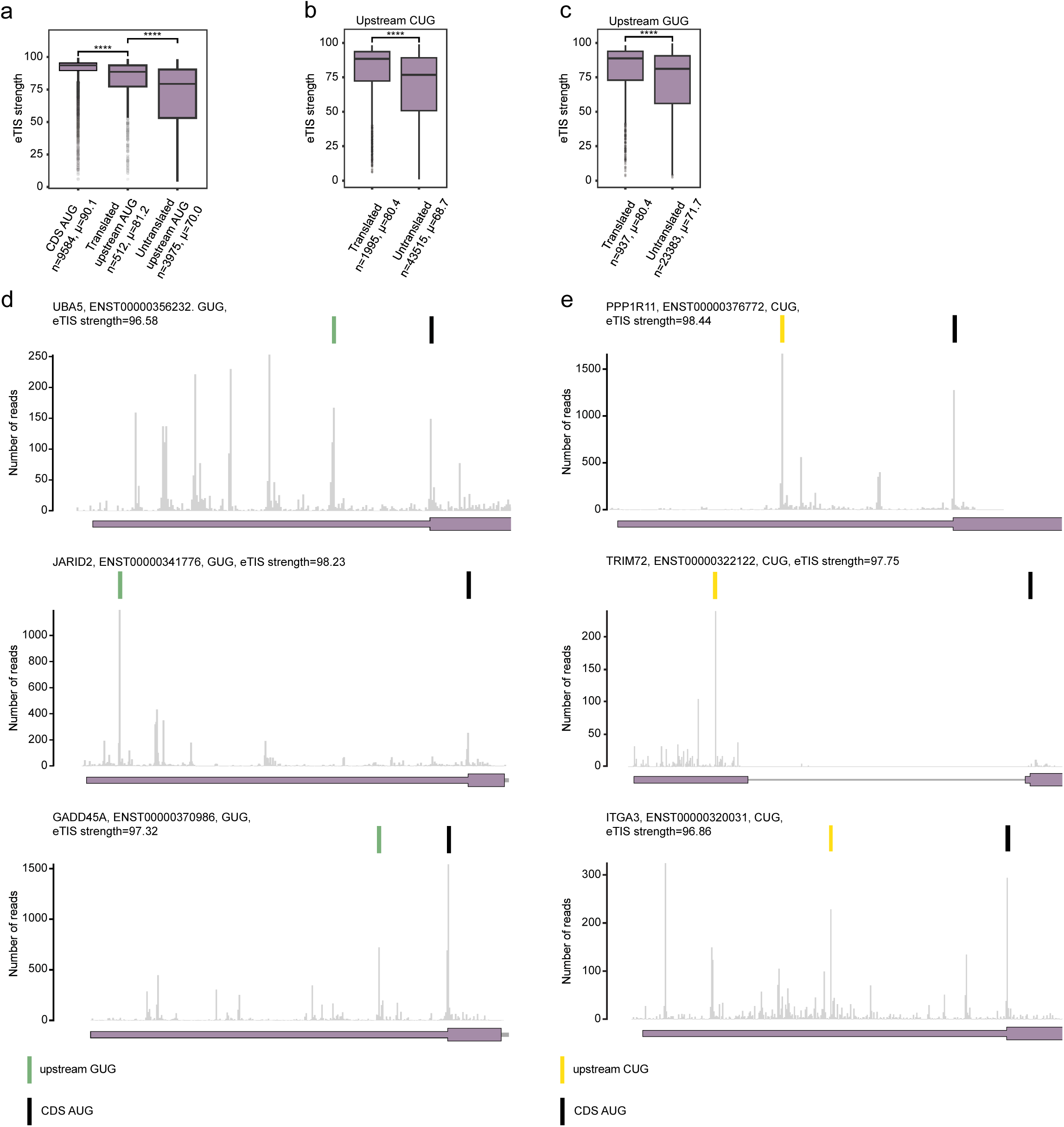
The influence of eTIS strength on start codon recognition in endogenous mRNAs. The RiboScanner model was used to predict eTIS strengths for different classes of start codons. **a**, Boxplot representing the eTIS strengths of CDS AUGs, experimentally validated translated upstream AUGs, as well as AUGs with no detectable translation (Mudge *et al*., 2022). **b**, **c**, Same as (**a**) for near-cognate CUG (**b**) and GUG (**c**) start codons. Boxplots display the median (center line) and interquartile range (25th–75th percentiles; box), with whiskers extending to 1.5x the interquartile range. Data points beyond the whiskers are plotted as outliers. Significance was calculated using a Wilcoxon rank-sum test. **** denotes *P* < 0.0001. **d**, **e**, Examples of upstream near-cognate start codons that show experimental evidence of translation in translation initiation profiling experiments (Michel *et al*., 2018) (See Methods). The canonical protein coding start codon is indicated by a black line, CUG and GUG start codons are indicated by yellow and green lines, respectively. The eTIS strength as determined by the RiboScanner model of each near-cognate start codon is indicated above the corresponding plot.

**Supplementary Figure 7.**
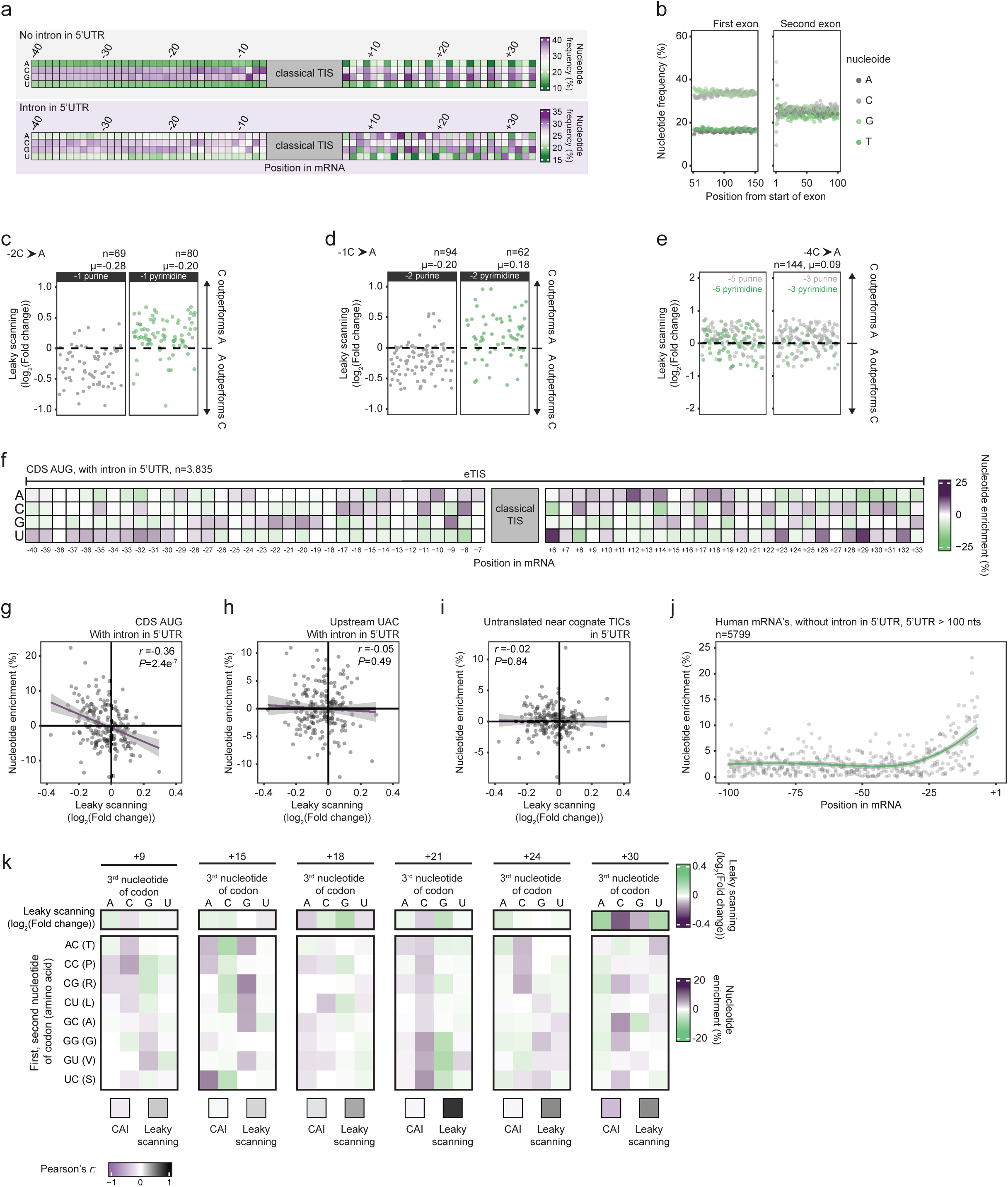
Genome-wide nucleotide enrichments in the eTIS. **a**, Heatmap of the frequency of each nucleotide at indicated positions relative to the TIS. Nucleotide frequencies for genes without (Top) or with (Bottom) intron in the 5’UTR is shown. Values are re-scaled from Fig. 3b to better illustrate small differences. **b,** Nucleotide frequency in the first exon (left) and second exon (right). Nucleotide frequencies remain constant across both exons. The first exon is strongly enriched for GC content, whereas the second exon does not show a pronounced GC bias. The spike at the left side of the second exon plot likely reflects nucleotide bias introduced by the splice site. Frequencies were calculated using only 5′ UTR sequences. The first 50 nucleotides of the first exon were excluded from the plot because they display strong nucleotide bias, likely due to enrichment of mRNA cap-associated sequences. **c**, The effect of the -2C to A substitution on leaky scanning is dependent on the identity of the nucleotide at the -1 position. When -1 is a purine, adenine at -2 reduces leaky scanning relative to cytosine (left), when -1 is a pyrimidine, cytosine at -2 reduces leaky scanning relative to adenine (right). Each dot corresponds to an mRNA molecule carrying the specified substitution at the indicated position. **d**, Same as (**c**) for a -1C to A substitution, with data separated by nucleotide type (purine, pyrimidine) at the –2 position. **e**, same as (**c**) for a -4C to A substitution, with data colored by nucleotide type at the –5 (left) and –3 (right) positions. In the context of the −4C to A substitution, nucleotide identity at neither the −5 nor the −3 position influences leaky scanning. **f**, Heatmap representing the enrichment of each nucleotide at indicated positions relative to the start codon, normalized to local nucleotide frequencies and position within the codon (for nucleotides downstream of the start codon) (See Methods) for canonical CDS AUGs in transcripts with an intron in the 5’UTR. **g**, **h**, Comparison between the measured effect on leaky scanning and nucleotide enrichment for canonical CDS AUGs (**g**) and upstream UAC codons (**h**) for transcripts with an intron in the 5’UTR. **i**, Comparison between the measured effect on leaky scanning and nucleotide enrichment for upstream near-cognate start codons, for which translation could not be validated. Pearson correlation coefficients (*r*) and corresponding *P* values are indicated for each panel to assess the strength and significance of the linear relationships. **j**, Scatterplot depicting the absolute nucleotide enrichment for the first 100 nucleotides upstream of the start codon. For each position, all four nucleotides are plotted. Nucleotide enrichment is highest in the region of the eTIS. TIS bases (nucleotides -6 to -1) are excluded from the plot. The green line indicates a smoothed trend, and the grey shaded region denotes the 95% confidence interval. **k**, Heatmap depicting nucleotide enrichments for the third nucleotide of the codon (wobble base) for amino acids whose codons permit any of the four nucleotides at the wobble base. Position relative to the start codon is shown on top. The effect of each nucleotide on start codon recognition is shown in the top row (replotted from Fig. 2b). The two lower boxes report Pearson correlation coefficients between nucleotide enrichment and CAI (left), and between nucleotide enrichment and the effect on leaky scanning (right). Nucleotide enrichments correlate better with start codon recognition efficiency than with codon optimality.

**Supplementary Figure 8.**
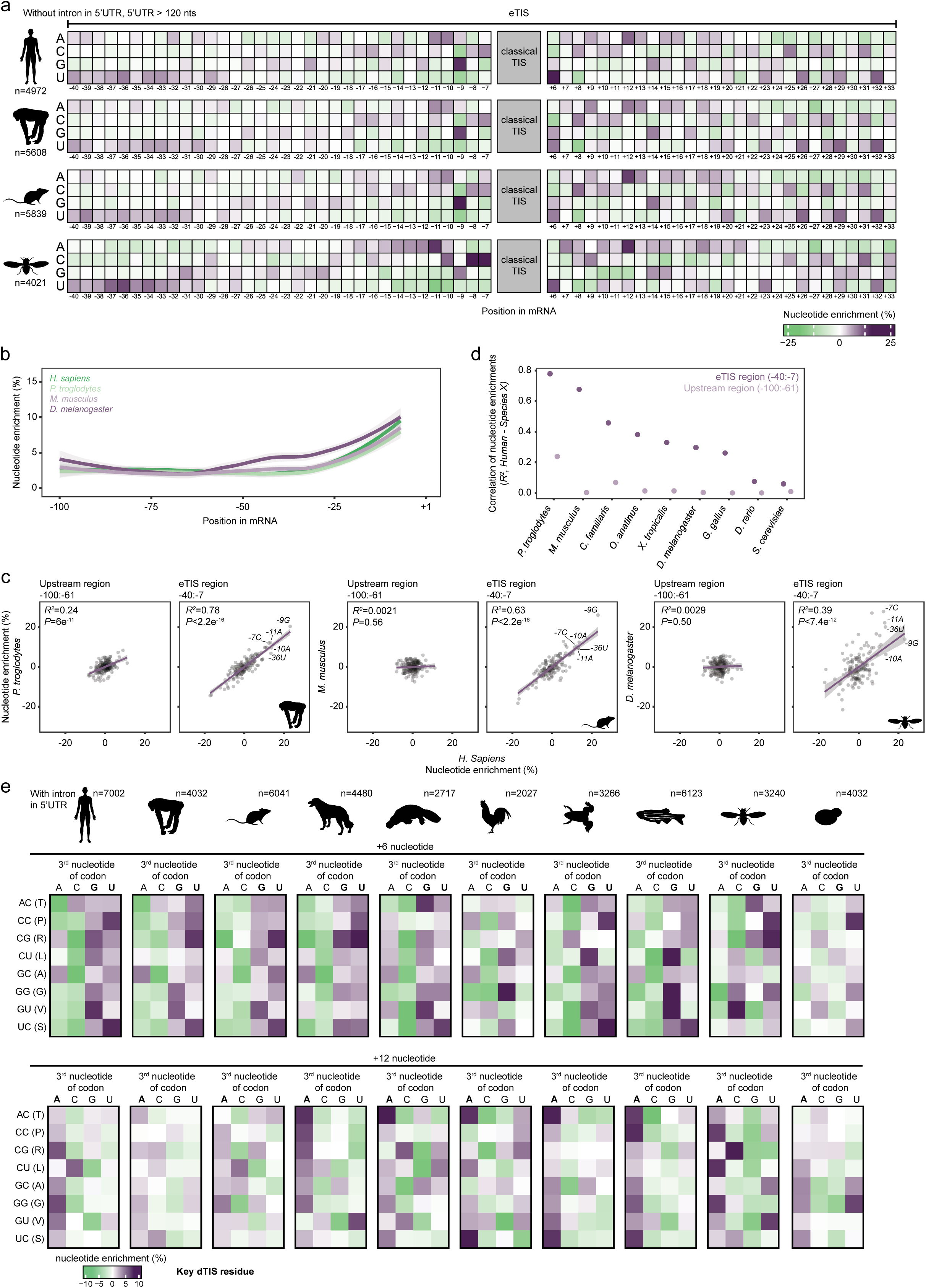
Enrichment of eTIS residues is evolutionary conserved. **a**, Heatmap representing the enrichment of each nucleotide at indicated positions relative to the start codon, normalized to local nucleotide frequencies and position within the codon (for nucleotides downstream of the start codon) (See Methods) for canonical CDS AUGs in transcripts without a 5’UTR intron for *H. Sapiens*, *P. troglodytes*, *M. musculus* and *D. Melanogaster*, respectively. **b**, Plot depicting the absolute nucleotide enrichment for the first 100 nts upstream of the start codon for all transcripts without a 5’UTR intron and a minimal 5’UTR length of 100 nts for the different indicated species. Nucleotide enrichment is highest in the region of the eTIS. TIS bases (nucleotides -6 to -1) are excluded from the plot. The lines indicate the smoothed trend lines, with the grey shaded regions indicating the 95% confidence intervals. **c**, Scatterplots illustrating the correlations between nucleotide enrichment in *H. sapiens* and the indicated species. Nucleotide enrichment values were calculated for the eTIS region (nucleotides -40 to -7, right panels) and as a control for the upstream 5’UTR region not involved in start codon recognition (see Fig. 2a) (nucleotides -100 to -61, left panel). Linear regression fits are shown in purple, with grey shaded regions indicating the 95% confidence intervals. Coefficients of determination (R²) and corresponding *P* values are indicated for each panel to assess the strength and significance of the linear relationships. Examples of highly enriched nucleotides in the eTIS region are indicated. **d**, Comparison of the coefficient of determination (R^2^) for nucleotide enrichment between *H. sapiens* and the indicated species in the eTIS region or upstream 5’UTR region. **e**, Heatmaps depicting nucleotide enrichments for the third nucleotide of the codon (wobble base) for amino acids whose codons permit any of the four nucleotides at the wobble base, across the indicated species (*H. sapiens*, *P. troglodytes, M. musculus*, *C. lupus familiaris*, *A. anatinus*, *G. gallus*, *X. laevis*, *D. rerio*, *D. melanogaster, S. cerevisiae*). Nucleotide position relative to the start codon is shown on top, key distal eTIS residues are highlighted in bold.

**Supplementary Figure 9.**
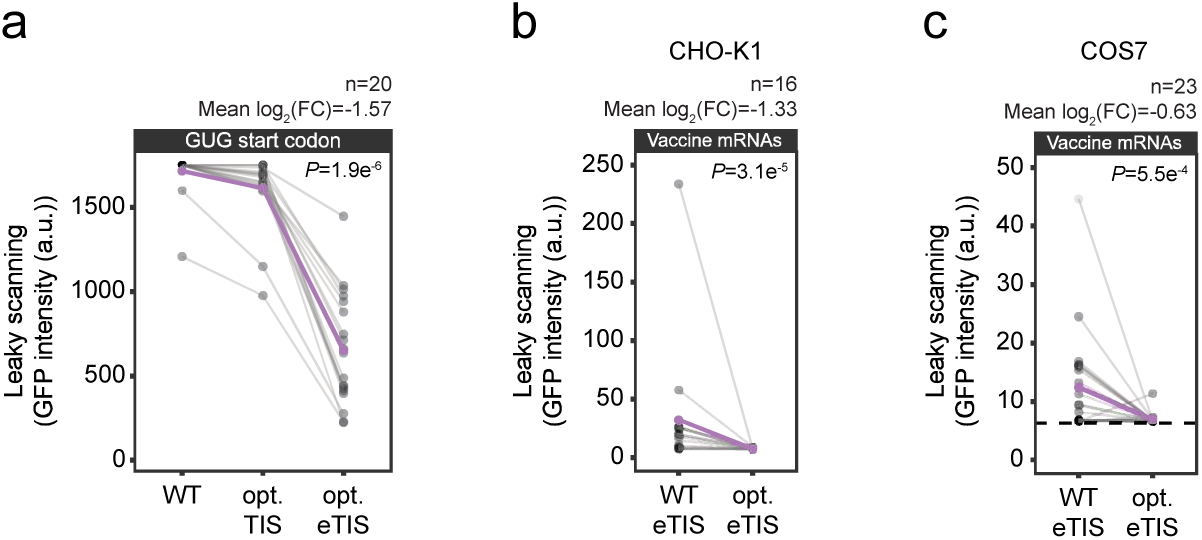
eTIS optimization for therapeutic mRNAs. **a**, Either TIS or eTIS optimization for mRNAs with a GUG start codon. Efficient recognition of the near-cognate start codon requires optimization of the full eTIS. **b**, **c**, eTIS optimization of vaccine mRNAs reduces leaky scanning in CHO-K1 (**b**) and COS-7 (**c**) cells. The purple lines in **a-c** represent the mean changes in leaky scanning upon (e)TIS optimization. *P* values were determined using a paired Wilcoxon signed-rank test. Dashed line indicates the median fluorescence intensity of the lowest FACS gate, representing the lowest possible GFP expression value in the RiboScan assay.

**Supplementary Figure 10.**
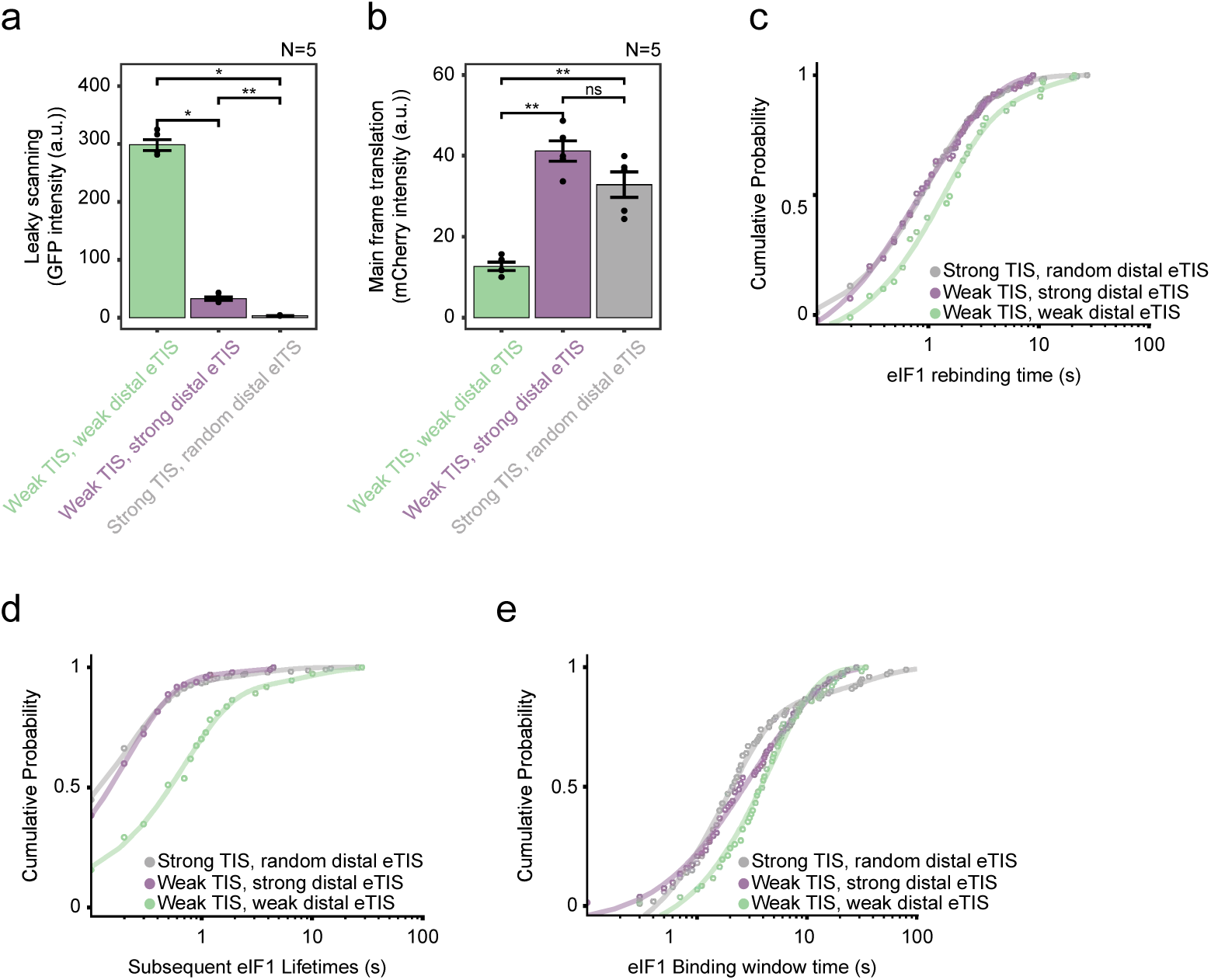
The eTIS modulates the kinetics of start codon recognition. **a**, GFP expression of the RiboScan reporter, representing leaky scanning, *in vivo* in HEK293T cells for mRNA sequences used for *in vitro* single-molecule imaging experiments. **b**, mCherry expression of the RiboScan reporter, representing main frame start codon recognition, in HEK293T cells for mRNA sequences used for *in vitro* single-molecule imaging experiments. Bars represent the mean of five biological replicates (N = 5), with individual replicates shown as dots. Statistical significance was assessed using a Welch’s two-sample t-test. * denotes *P* < 0.05, ** *P* < 0.01; ns, not significant. **c**, Time from initial departure of eIF1 until eIF1 rebinding is show for indicated mRNAs. **d**, eIF1 binding times of all eIF1 binding events after the first eIF1 release. **e**, Time from initial eIF1 binding until final eIF1 departure.

**Supplementary Figure 11.**
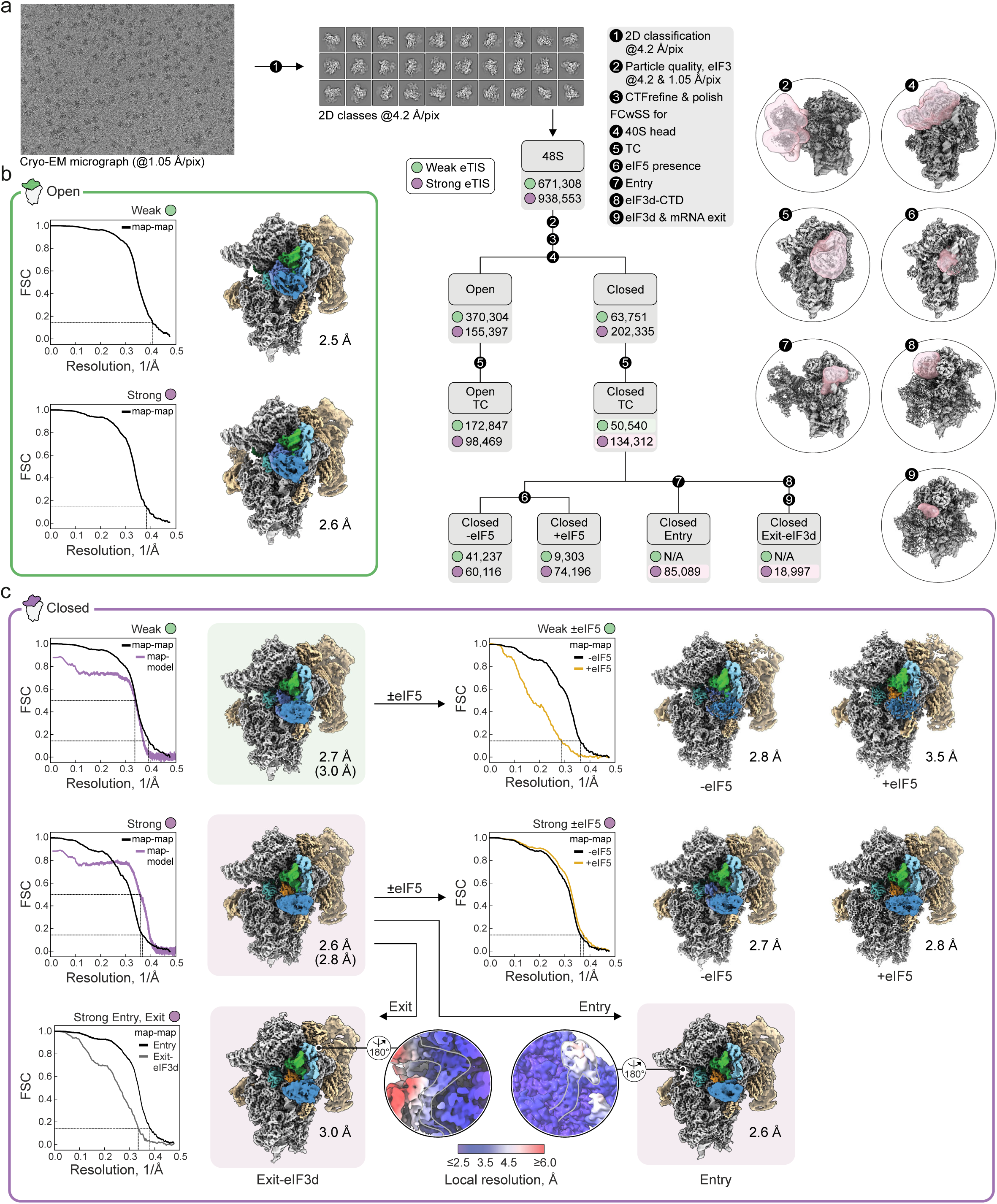
Cryo-EM analysis of distal eTIS function. **a**, Cryo-EM data processing workflow with representative micrograph and 2D class averages. Classification steps and masks (transparent pink) used for local sorting are indicated; particle numbers are shown for 48S complexes assembled on weak and strong distal eTIS mRNAs. Data processing was performed at a final pixel size of 1.05 Å per pixel unless stated otherwise (see Methods). **b**, Half-map Fourier shell correlation (FSC) curves and cryo-EM maps of open 48S complexes. **c**, FSC curves and cryo-EM maps of closed 48S states, showing half-map FSCs (black and yellow), and, where applicable, model-map FSCs for atomic models (light purple). Close-ups show local resolution estimates for the exit (left) and entry (right) regions from the “exit-eIF3d” and “entry” maps obtained by local classification; grey contours indicate the path of eIF3d at the exit and mRNA-eIF3g at the entry. In the closed 48S + eIF5 with weak eTIS mRNA, eIF5 density remained weak after focused classification, suggesting an even lower fraction of eIF5-bound particles in the weak eTIS dataset than estimated from the classification. Maps in **b** and **c** were rendered using model-free LocScale. For closed 48S±eIF5 with weak eTIS mRNA, sharpened maps are shown with flexible elements low-pass filtered. Map resolutions are given in Å, with model resolutions in parentheses; particle populations and maps used for atomic modelling are highlighted by light green and light purple background shading.

**Supplementary Figure 12.**
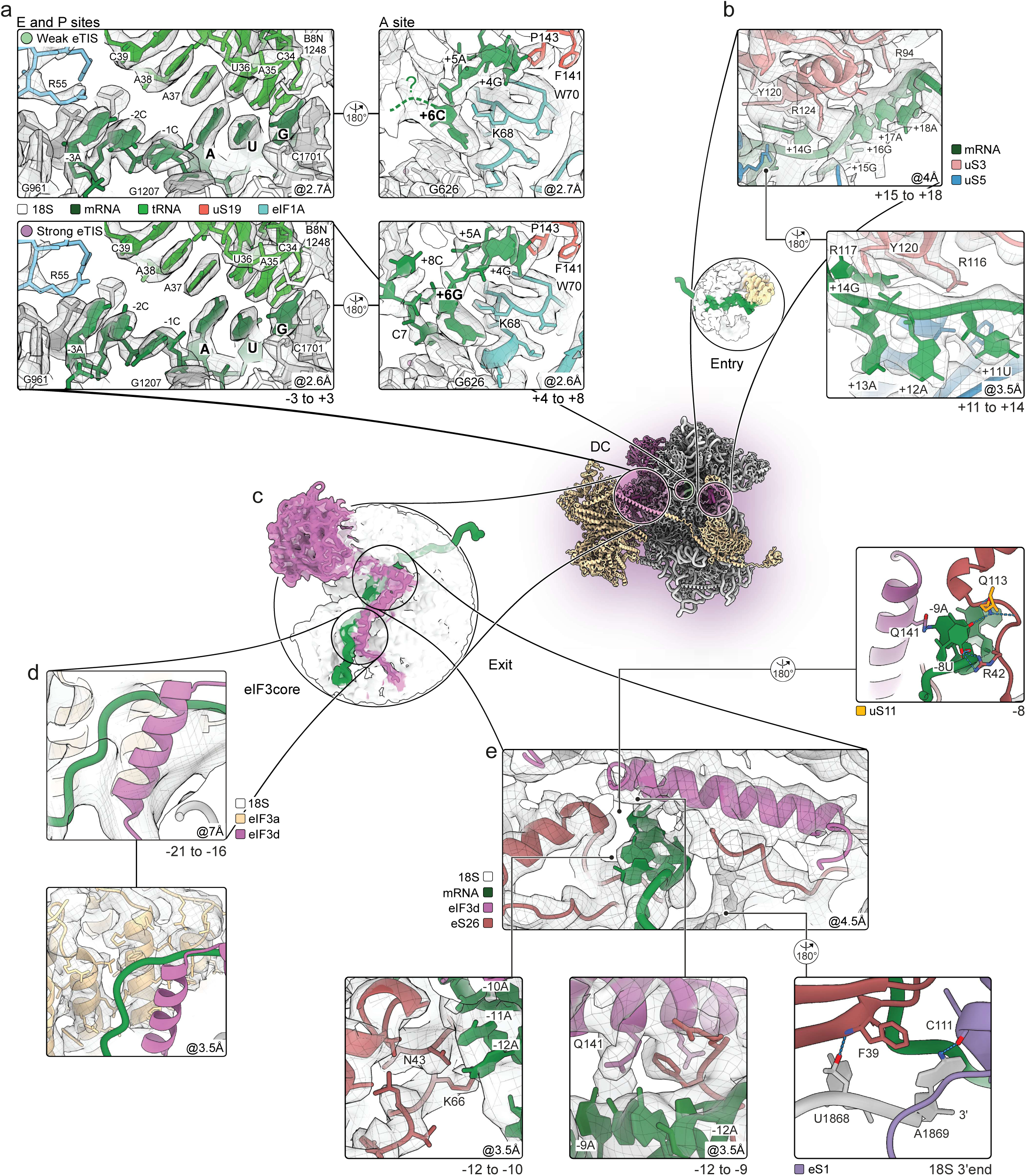
Details of 48S–eTIS interactions. **a**, Experimental densities at the E and P sites (left) and A site (right) of closed 48S complexes assembled on weak (top) and strong eTIS mRNAs (bottom). DC, decoding center; 18S, 18S rRNA. **b**, Densities at mRNA entry channel from locally classified map (top) and the overall map (bottom). **c**, Density of the eIF3core-exit region obtained by local classification, low-pass filtered to emphasize the path of eIF3d. **d**, Close-ups of density in the eIF3core region. Top, low-pass filtered map used to model a flexible eIF3d α-helix (residues 89-103); the mRNA backbone was traced at 5-6 Å resolution in this region, resulting in local uncertainty in register (±1 nucleotide). Bottom, higher resolution density for eIF3a. **e**, Close-ups of the mRNA exit channel with major eIF3d-eTIS contacts. mRNA was interpreted at 4.5 Å in this region (center), with protein side-chains resolved at higher resolution (bottom left and center). Top right, details of the +8U interaction network. Bottom right, contacts of the 18S 3’ terminus with eS1 and eS26. Resolution cut-offs used for low-pass filtering of maps for visualization and model building are indicated in Å.

**Supplementary Table 1.** Reporter overview, DNA primers and single reporter sequences.

**Supplementary Table 2.** GFP intensity values underlying individual figure panels.

**Supplementary Table 3.** Source data for (e)TIS heatmaps.

**Supplementary Table 4.** Data related to the RiboScanner model.

**Supplementary Table 5.** Sequences and eTIS strengths underlying Fig. 3d,g and Supplementary Fig. 6a–c.

**Supplementary Table 6.** Datasets from external sources and genome assemblies used in this study.

Ribosome initiating profiling datasets
| Publication | Cell line |
| --- | --- |
| Thousands of novel translated open reading frames in humans inferred by ribosome footprint profiling (Raj et al. 2016) | LCL |
| Positional proteomics reveals differences in N-terminal proteoform stability (Gawron et al. 2016) | Jurkat |
| Pervasive functional translation of noncanonical human open reading frames (Chen et al. 2020) | iPSC |
| Genome-wide identification and differential analysis of translational initiation (Zhang et al. 2017) | HEK |
| Quantitative profiling of initiating ribosomes in vivo (Xiangwei et al. 2014) | HEK |
| PROTEOFORMER: deep proteome coverage through ribosome profiling and MS integration (Crappe et al. 2015) | HCT116 |
| eIF1 modulates the recognition of suboptimal translation initiation sites and steers gene expression via uORF (Fijalkowska et al. 2017) | HTC116 |
| Many lncRNAs, 5'UTRs, and pseudogenes are translated and some are likely to express functional proteins (Ji et al. 2015) | BJ/MCF10A |

ORF annotations
| Publication |  |
| --- | --- |
| Standardized annotation of translated open reading frames (Mudge et al. 2022) | Identification of upstream CUG/GUG codons used as start codons |

TSS mapping
| Publication |  |
| --- | --- |
| Extensive transcriptional heterogeneity revealed by isoform profiling (Pelechano et al. 2013) | 5'end mapping using TIF-seq in *S. cerevisiae* |

List of genomes
| Species | Ensembl Assembly (Accession) |
| --- | --- |
| *Canis lupus familiaris* | ROS_Cfam_1.0 (GCA_014441545.1) |
| *Danio rerio* | GRCz11 (GCA_000002035.4) |
| *Drosophila melanogaster* | BDGP6.54 (GCA_000001215.4) |
| *Gallus gallus* | bGalGal1.mat.broiler.GRCg7b (GCA_016699485.1) |
| *Homo sapiens* | GRCh38.p14 (GCA_000001405.29) |
| *Mus musculus* | GRCm39 (GCA_000001635.9) |
| *Ornithorhynchus anatinus* | mOrnAna1.p.v1 (GCA_004115215.2) |
| *Pan troglodytes* | Pan_tro_3.0 (GCA_000001515.5) |
| *Saccharomyces cerevisiae* S288c | R64-1-1 (GCA_000146045.2) |

**Supplementary Table 7.** Kinetics summary of single-molecule imaging experiments.

| mRNA | # of traces | # of binding events | First eIF1 lifetime |  | Subsequent eIF1 lifetime |  |  | eIF1 rebinding |  |  |
| --- | --- | --- | --- | --- | --- | --- | --- | --- | --- | --- |
| | | | $k_{\text{off, fast}}$ (s<sup>-1</sup>) | $t_{1/2}$ (s) | $k_{\text{off, fast}}$ (s<sup>-1</sup>) (%) | $k_{\text{off, slow}}$ (s<sup>-1</sup>) | $t_{1/2}$ (s) | $k_{\text{on, fast}}$ (s<sup>-1</sup>) (%) | $k_{\text{on, slow}}$ (s<sup>-1</sup>) | $t_{1/2}$ (s) |
| Strong TIS, random distal eTIS | 101 | 331 | 0.54 ± 0.02 | 1.9 ± 0.1 | 4.9 ± 0.5 (92%) | 0.39 ± 0.19 | 0.4 ± 0.1 | 1.1 ± 0.1 (78%) | 0.27 ± 0.08 | 1.5 ± 0.2 |
| Weak TIS, strong distal eTIS | 103 | 229 | 0.61 ± 0.05 | 1.6 ± 0.1 | 4.8 ± 0.6 (95%) | 0.49 ± 0.59 | 0.3 ± 0.1 | 2.4 ± 0.9 (40%) | 0.50 ± 0.09 | 1.4 ± 0.2 |
| Weak TIS, weak distal eTIS | 64 | 101 | 0.35 ± 0.01 | 2.9 ± 0.1 | 1.4 ± 0.3 (88%) | 0.12 ± 0.16 | 1.6 ± 0.7 | 0.74 ± 0.23 (81%) | 0.12 ± 0.13 | 2.7 ± 1.0 |

| mRNA | eIF1 binding window |  | eIF5B binding after eIF1 binding window |  |
| --- | --- | --- | --- | --- |
| | $k_{\text{off, fast}}$ (s<sup>-1</sup>) | $t_{1/2}$ (s) | $k_{\text{on, fast}}$ (s<sup>-1</sup>) | $t_{1/2}$ (s) |
| Strong TIS, random distal eTIS | 0.44 ± 0.04 | 2.3 ± 0.2 | 0.29 ± 0.02 | 3.4 ± 0.2 |
| Weak TIS, strong distal eTIS | 0.26 ± 0.02 | 3.8 ± 0.2 | 0.37 ± 0.04 | 2.7 ± 0.2 |
| Weak TIS, weak distal eTIS | 0.23 ± 0.02 | 4.3 ± 0.3 | 0.17 ± 0.01 | 5.9 ± 0.3 |

**Supplementary table 8.** Cryo-EM structure determination – closed 48S.

| Ribosomal complex | Closed 48S, weak eTIS | Closed 48S, strong eTIS |
| --- | --- | --- |
| EM Data Bank and PDB identifiers | EMD-YYYYY<br>XXXX | EMD-YYYYY<br>XXXX |
| **Data collection and processing** |  |  |
| Microscope | Titan Krios | Titan Krios |
| Camera | Gatan K3 | Gatan K3 |
| Magnification | 81,000 | 81,000 |
| Voltage (kV) | 300 | 300 |
| Electron exposure (e<sup>-</sup> / Å<sup>2</sup>) | 40 | 40 |
| Defocus range (μm) | 0.5-1.5 | 0.5-1.5 |
| Pixel size (Å) | 1.05 | 1.05 |
| Symmetry imposed | C1 | C1 |
| Initial particles (no.) | 671,308 | 938,553 |
| Final particles (no.) | 50,540 | 134,312 |
| Map resolution* (Å) | 2.7 | 2.6 |
| FSC threshold | 0.143 | 0.143 |
| **Refinement** |  |  |
| Initial models used (PDB code) | 8PJ2, 8QOI, 7A09 | 8PJ2, 8QOI, 7A09 |
| Model resolution (Å) | 3.0 | 2.8 |
| FSC threshold | 0.5 | 0.5 |
| Model resolution range (Å) | 3.0-6 | 2.8-6 |
| Map sharpening B factor (Å<sup>2</sup>) | -42.1 | -45.8 |
| **Model composition: non-hydrogen atoms/residues/RSCC** |  |  |
| Total | 119,602/12,709/0.62 | 120,522/12,813/0.66 |
| RNA | 40,023/1,875/0.82 | 40,087/1,877/0.82 |
| Proteins | 79,488/10,743/0.59 | 80,285/10,837/0.63 |
| Cumulative RSCC (%) >0.8/>0.6/>0.4 | 0.50/0.63/0.71 | 0.54/0.66/0.74 |
| **B factors (Å<sup>2</sup>)** |  |  |
| Protein | 70.50 | 70.47 |
| Nucleotide | 53.09 | 53.09 |
| Ligands and ions | 39.18 | 39.18 |
| *R. m. s. deviations* |  |  |
| Bond lengths (Å) | 0.006 | 0.006 |
| Bond angles (°) | 1.082 | 0.872 |
| **B factors (Å<sup>2</sup>)** |  |  |
| MolProbity score | 1.30 | 1.31 |
| Clashscore | 2.61 | 2.68 |
| Rotamer outliers (%) | 0.00 | 0.00 |
| *Ramachandran plot* |  |  |
| Favored (%) | 96.23 | 96.24 |
| Allowed (%) | 3.77 | 3.76 |
| Disallowed (%) | 0.00 | 0.00 |

\*Resolution used for final atomic model refinement; model building and interpretation were carried out using low-pass filtered and local maps as described in Methods.

**Supplementary Table 9.** Cryo-EM structure determination – open 48S and closed 48S ±eIF5.

| Ribosomal complex | Open 48S<br>weak eTIS | Closed 48S -eIF5<br>weak eTIS | Closed 48S +eIF5<br>weak eTIS | Open 48S<br>strong eTIS | Closed 48S -eIF5<br>strong eTIS | Closed 48S +eIF5<br>strong eTIS |
| --- | --- | --- | --- | --- | --- | --- |
| EM Data Bank identifier | EMD-YYYYY | EMD-YYYYY | EMD-YYYYY | EMD-YYYYY | EMD-YYYYY | EMD-YYYYY |
| **Data collection and processing** |  |  |  |  |  |  |
| Microscope | Titan Krios | Titan Krios | Titan Krios | Titan Krios | Titan Krios | Titan Krios |
| Camera | Gatan K3 | Gatan K3 | Gatan K3 | Gatan K3 | Gatan K3 | Gatan K3 |
| Magnification | 81,000 | 81,000 | 81,000 | 81,000 | 81,000 | 81,000 |
| Voltage (kV) | 300 | 300 | 300 | 300 | 300 | 300 |
| Electron exposure (e<sup>-</sup> / Å<sup>2</sup>) | 40 | 40 | 40 | 40 | 40 | 40 |
| Defocus range (μm) | 0.5-1.5 | 0.5-1.5 | 0.5-1.5 | 0.5-1.5 | 0.5-1.5 | 0.5-1.5 |
| Pixel size (Å) | 1.05 | 1.05 | 1.05 | 1.05 | 1.05 | 1.05 |
| Symmetry imposed | C1 | C1 | C1 | C1 | C1 | C1 |
| Initial particles (no.) | 671,308 | 671,308 | 671,308 | 938,553 | 938,553 | 938,553 |
| Final particles (no.) | 172,847 | 50,050 | 13,521 | 98,469 | 60,116 | 74,196 |
| Map resolution (Å) | 2.5 | 2.8 | 3.5 | 2.6 | 2.7 | 2.8 |
| FSC-threshold | 0.143 | 0.143 | 0.143 | 0.143 | 0.143 | 0.143 |
| Sharpening B factor (Å<sup>2</sup>) | -44.0 | -42.7 | -54.2 | -49.0 | -46.3 | -46.8 |

**Supporting Figure 1.**
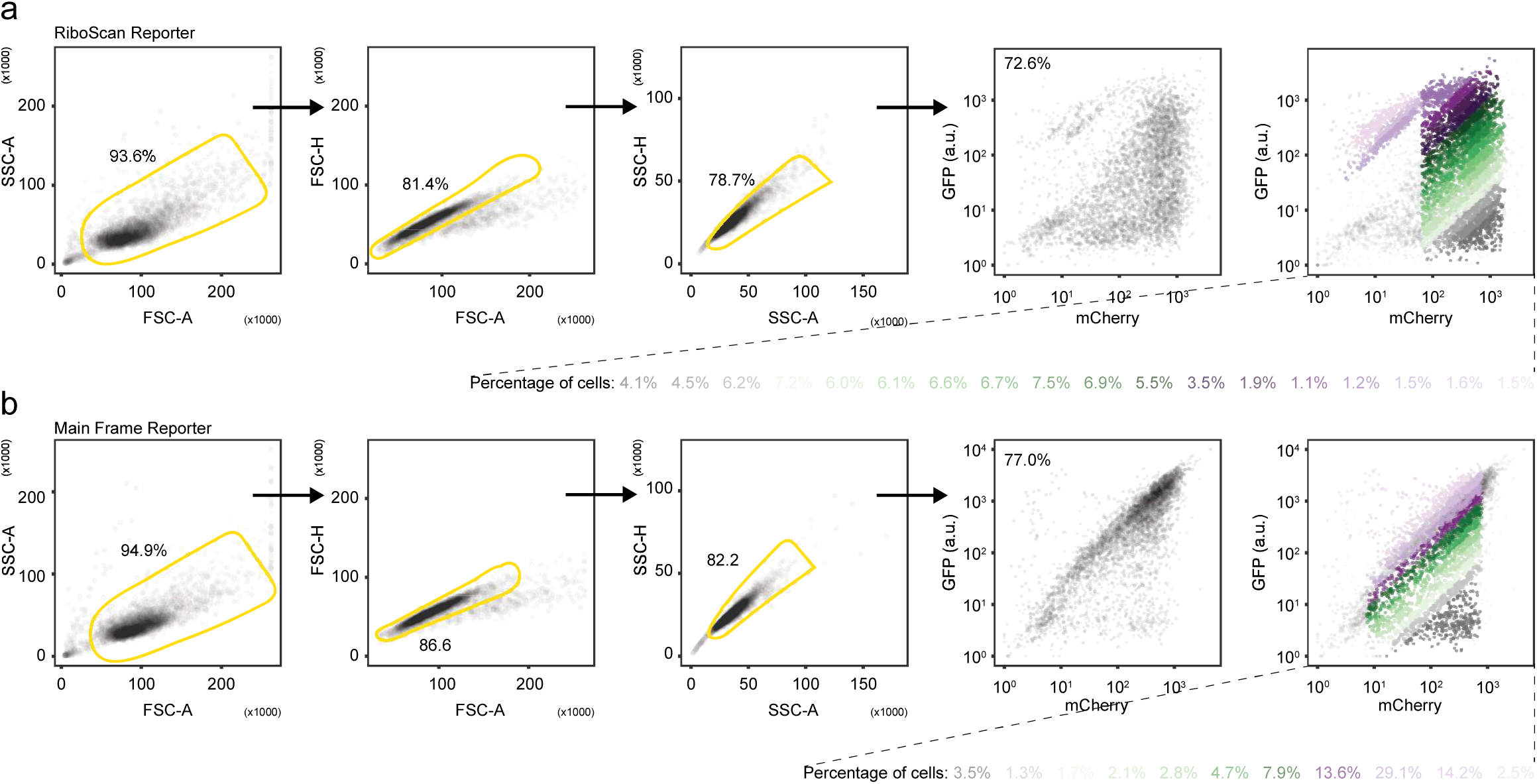
FACS gating strategy for RiboScan and Main Frame Reporter FACS-seq experiments. **a, b** FACS gating strategy used for the RiboScan (**a**) and Main Frame reporter (**b**) FACS-seq experiments. Representative FACS plots are shown. Cells were sorted into different bins based on their GFP/mCherry fluorescence ratio, as indicated by the different colors in the rightmost panels.

